# Adaptive-learning physics-aware light-field microscopy enables day-long and millisecond-scale super-resolution imaging of 3D subcellular dynamics

**DOI:** 10.1101/2023.03.15.532876

**Authors:** Lanxin Zhu, Jiahao Sun, Chengqiang Yi, Meng Zhang, Yihang Huang, Sicen Wu, Mian He, Liting Chen, Yicheng Zhang, Chunhong Zheng, Hao Chen, Yuhui Zhang, Dongyu Li, Peng Fei

## Abstract

Long-term and high-spatiotemporal-resolution 3D imaging of living cells remains an unmet challenge for super-resolution microscopy, owing to the noticeable phototoxicity and limited scanning speed. While emerging light-field microscopy can mitigate this issue through three-dimensionally capturing biological dynamics with merely single snapshot, it suffers from suboptimal resolution insufficient for resolving subcellular structures. Here we propose an Adaptive Learning PHysics-Aware Light-Field Microscopy (Alpha-LFM) with a physics-aware deep learning framework and adaptive-tuning strategies capable for highly-generalizable light-field reconstruction of diverse subcellular dynamics. Alpha-LFM delivers sub-diffraction-limit spatial resolution (∼120 nm) while maintaining high temporal resolution and low phototoxicity. It enables rapid (at hundreds of volumes per second), long-term (up to 60 hours) 3D super-resolution imaging of diverse intracellular dynamics with exceptional details. Using Alpha-LFM approach, we finely resolve the lysosome-mitochondrial interactions, capture rapid motion of peroxisome and the endoplasmic reticulum, and reveal the variations in mitochondrial fission activity throughout two complete cell cycles.

## Introduction

In living cells, different organelles work together to execute diverse and intricate physiological functions. Elucidating these fast dynamics / interactions of organelles across the cell cycle needs the microscopes to have sufficiently high spatiotemporal resolution in four dimensions (time +3D space) and low phototoxicity rate for long-term observation ^1–4^. This is highly challenging because current 3D microscopy techniques are constrained to the maximal photon budget a sample permits, which results in an inevitable trades-off among imaging speed, spatial resolution, and photon efficiency^5–12^. A series of 3D microscopy implementations including confocal microscopy^11, 13^, 3D structured illumination microscopy (3D SIM)^5, 7, 12^ and light sheet microscopy (LSM)^9, 14^, have been intensively developed for live-cell imaging beyond diffraction limit, e.g., ∼150 nm lateral and ∼280- nm axial resolutions by SIM mode of lattice light-sheet microscope (LLSM-SIM)^9^. However, these scanning-based super-resolution (SR) approaches often require recording hundreds of frames at different planes to reconstruct a super-resolved volume, thereby showing compromised temporal resolution limited to a few seconds and relatively high phototoxicity limited to hundreds of volumes acquisition^9^. Therefore, it’s difficult for these approaches to observe either long-term evolution during the cell cycle or instantaneous subcellular process occurred at milliseconds timescale.

Unlike scanning-based microscopy, light-field microscopy (LFM) provides photon-efficient direct volumetric imaging by encoding both position and angular information of 3D signals on single 2D camera snapshots without time-consuming axial scanning ^15–20^. Benefiting from the high-speed scanning-free 3D imaging, LFM has facilitated various biological studies on neural activities, cardiac hemodynamics and live cells^18, 19, 21^. In contrast to the superior temporal resolution and low phototoxicity well suited for live imaging, the spatial resolution of LFM is often unsatisfactory, owing to the insufficient pixel sampling and limited sub numerical aperture existed in each simultaneously captured light-field view. Many efforts have been made to improve this limitation of LFM^19, 22, 23^, for example, DAOSLIMIT can yield a spatial resolution of ∼220 nm after a 9-times aperture scanning that slightly lower the imaging speed^19^. Meanwhile, the use of iterative light-field deconvolution for 2D-3D image reconstruction is vulnerable to artifacts, and incapable of surpassing the diffraction limit. There always exists an “impossible performance triangle” which greatly limits the design space of current 3D fluorescence microscopy techniques towards high speed, high spatial resolution and high photon efficiency imaging.

The recent advent of deep neural networks for image reconstruction enlarges the design space of microscopy by introducing prior knowledge of high-resolution data for learning and inference^24,25^. We previously report view-channel-depth light-field microscopy (VCD-LFM), in which a VCD network model is trained to learn the nonlinear relationships between the 3D confocal ground truths and their 2D light-field projections and afterwards, can directly reconstruct high-resolution 3D volume from a single 2D light-field image through using the well-trained channels in the VCD model to transform the implicit features of the light-field views into depth information of a 3D stack. With deep-learning model combining the high-resolution advances of scanning microscopy into high-speed imaging of LFM, VCD-LFM holds the promise for high-speed and high-resolution 4D imaging of live samples^26–28^. However, current end-to-end supervised networks encounter constraints in term of both enhanced capabilities and generalization ability. When dealing with complex inverse problems, ensuring accuracy is challenging. For instance, reconstructing a 3D SR image from a 2D under-sampled light-field image with spatial bandwidth compressed by around 600 times, requires the recovery of various degradations brought by noise, resolution, and dimensionality reduction. This presents a highly intricate ill-posed problem with a huge solution space. The fitting ability of network depends on the model complexity, the amount of data prior available and loss constraints^29^. However, traditional one-stage end-to-end networks, with limited amount of label data and model complexity, is difficult to find precise SR solutions in such a huge space, resulting in either limited resolution or reduced fidelity. Another typical challenge that supervised learning has to face is the generalization. This arises from the network’s predominant focus on learning high-dimensional features of specific training samples, thereby limiting their applicability to the new samples.

Here we propose an Adaptive Learning PHysics-Aware Light-Field Microscopy (Alpha-LFM), capable of accurately super-resolved and highly-generalizable reconstruction of diverse subcellular structures. To enhance the network’s fitting capability, we established an adaptive-learning physics-aware network framework (Alpha-Net) to increase model complexity and the data constraint through decoupling the complex light field inverse problem into multiple subtasks with multi-stage data guidance. Instead of simply incorporating a 3D SR Net with VCD-Net, our decomposition strategy is designed to progressively denoise, de-aliase and reconstruct, facilitating a more precise 3D reconstruction by effectively leveraging the angular information in light-field (LF) views. To permit the implementation of this decomposition strategy, we developed a physics-embedded hierarchical data synthesis pipeline to introduce multi-stage data prior and decomposed-progressive optimization strategy to enable the convergence of multi-stage networks. We demonstrate that through carefully designing the model and training strategies, our Alpha-Net can notably narrow down the inversion space for seeking the correct solution more efficiently, enabling 3D SR reconstruction of diverse samples with high fidelity. To tackle unseen structures, we further developed an adaptive tuning strategy that allows for instant optimization for new live samples using 2D wide-field images. Alpha-LFM demonstrates imaging of the dynamics of intracellular structures in live cells at an isotropic spatial resolution of ∼120 nm and hundreds of hertz volume rate, facilitating the analysis of the lysosome-mitochondrial interactions as well as the rapid motion of peroxisome and the endoplasmic reticulum. With minimized phototoxicity of Alpha-LFM, we first achieve 3D live-cell imaging over 60 hours for the tracking of mitochondrial evolution across two complete cell cycles.

## Results

### Principle and Implementation of Alpha-LFM

Light-field microscopy encodes the spatial-angular patterns from 3D sample into a single 2D image^29, 30^. This procedure contains multiple imaging degradations including dimension compression from the diffraction-unlimited 3D objective to a diffraction-limited 2D LF projection, the frequency aliasing induced by the under-sampling of the microlens array (MLA) during the encoding of spatial-angular information and the noises mainly from the camera exposure (Fig. 1a). In our study, light-field reconstruction beyond diffraction limit needs to inverse this compression with a space-bandwidth product (SBP) expansion over 600 times (Method). This intricate and ill-posed inversion problem has resulted in an extensive solution space that maps the under-sampled LF measurement to the possible 3D SR solutions, thereby posing a big challenge to either deconvolution-based approaches without data priors or standard end-to-end DL models constrained by limited label data and model complexity^26^ (Supplementary Note 1). This leads to an unsatisfactory reconstruction performance especially when imaging the fine subcellular structures where both high spatial resolution beyond diffraction limit and high fidelity close to the ground truth are required for downstream tasks.

**Figure 1.**
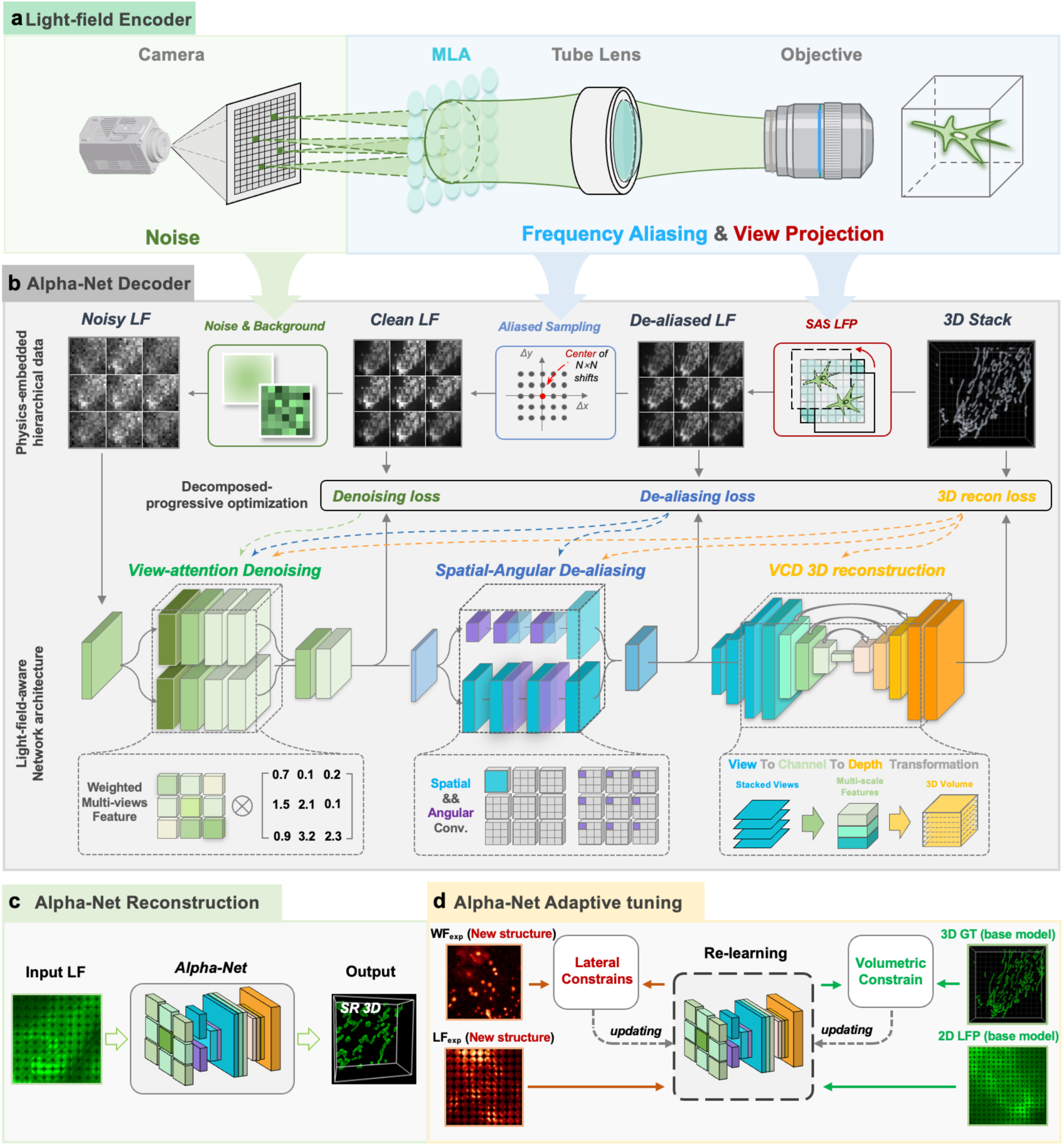
Design of Alpha-LFM. The principle and architecture of Alpha-LFM and its training strategy. **a,** The stepwise network is designed based on the physical model of the light-field imaging process. The light-field imaging process encodes multidimensional degradation mainly including: (i) the frequency aliasing caused by the limited NA of objectives and sub-apertures of microlens, (ii) dimension compression during light field encoding and (ii) the noise addition by the camera recording. **b,** Three light-field-aware sub-networks with view-attention denoising, spatial-angular de-aliasing and VCD 3D reconstruction restoration tasks are designed to disentangles and sequentially solves the complex light-field inversion problem. Multi-stage training data including De-aliased LF, Clean LF and Noisy LF are synthesized from the same 3D SR data using an “physics embedded hierarchical data synthesis”, to guide the training of the sub-networks. The decomposed-progressive optimization strategy ensures the collaboration of the sub-networks when training the multi-stage data. **c,** Direct Alpha-Net reconstruction when training and testing dataset are the same type of structures. **d,** The schematic illustrates the adaptive inference strategy of Alpha-LFM. When inferring previously unseen types of structures, experimentally-obtained 2D WF and corresponding LF images of the new structure are used to instantly tune the pre-trained model, making it adaptive to the unseen structures.

To improve the solving capability of the network and reduce the solution space, we established a physics-aware deep-learning framework to enhance the model complexity and the data constraint by disentangling the complex light field inverse problem into multiple subtasks with multi-stage data guidance and jointly optimization (Fig.1b, Supplementary Note 1, Supplementary Video 1, Method). Instead of simply using a 3D SR network to further improve the diffraction-limited resolution of VCD results, we devised a multistage network encompassing LF denoising, LF de-aliasing, and 3D reconstruction to progressively solve the light-field inverse problem. In this case, we are able to introduce more angular constraint in the de-aliasing network, therefore demonstrating notably enhanced reconstruction fidelity. Meanwhile, our decomposing strategy also achieves four-order-of-magnitude higher inference speed by avoiding the use of complex 3D blocks (Supplementary Fig. 1). To permit the implementation of this decomposed strategy, we firstly need to develop a sub-aperture shifted light-field projection (SAS LFP) strategy to generate light-field images without frequency aliasing (De-aliased LF) to guide the LF de-aliasing task, in which we projected the 3D SR images containing sub-aperture shifts into a series of clean light-field images (Clean LF) and rearrange them into a single one (Supplementary Fig. 2, Supplementary Note 1). This strategy serves as the cornerstone of our physics-embedded hierarchical data synthesis pipeline that allows semi-synthetic multi-stage data priors to be conveniently generated from the same 3D SR data based on the light-field model and then progressively guide the sub-networks (Fig. 1b, Method).

To achieve the SR light-field reconstruction that involves a 30-fold increase in sampling and transformation from 2D to 3D, we devised light-field-aware networks for each inversion task and a decomposed-progressive optimization strategy to efficiently facilitate the collaboration of multiple sub-networks, thus together ensuring the whole network can find an optimal solution with high resolution and high fidelity (Fig. 1b, Supplementary Note 1). To fully exploit the angular information from multiple views of LF image, we incorporated view-attention denoising modules, spatial-angular convolutional feature extraction operators and disparity constraints for high-fidelity (HiFi) denoising and de-aliasing of 4D (x, y, u, v) LF images, showing performance superior to modules that rely solely on spatial information (Supplementary Fig. 3, 4). We also optimized VCD 3D reconstruction sub-network by incorporating multi-res blocks to extract features from more dimensions, ensuring the high quality of the reconstruction results. Considering the different difficulty in each task, the three sub-networks are jointly optimized using our decomposed-progressive optimization (DPO) strategy, in which, each sub-network was optimized independently to ensure the high-quality solution for each sub-task at the beginning and then the sub-networks were grouped into “denoising”, “denoising and de-aliasing” and “denoising, de-aliasing and reconstruction” for being optimized progressively (Supplementary Fig. 5). DPO facilitates the collaboration of sub-networks while maintaining their independent training, thus achieving reconstructions with improved resolution and fidelity (Supplementary Fig. 6). It’s noteworthy that DPO strategy is also valuable to other image restoration networks. We also demonstrate the superior performance on other image restoration networks, such as RCAN networks^31^, for delivering SR results with higher fidelity (Supplementary Fig. 7). With these strategy, Alpha-Net can directly reconstruct high-fidelity SR images of diverse samples (Fig. 1c, Supplementary Fig. 8).

Another inherent limitation of supervised learning is the generalization ability. Networks are often unable to generalize to unseen types of structure because they primarily learn high-dimensional features of specific types of fluorescence signals during training, thereby limiting their applicability to other types of signals. Common solutions that either include training a large amount of data from diverse structures^22^ or use few 3D data for transfer learning^26^ need the acquisition of 3D high-resolution data on static sample in advance, suffering from low flexibility. We developed an adaptive-tuning strategy in Alpha-LFM to reconstruct the types of structure that were not included in the training datasets. When facing unseen types of structures, Alpha-LFM adopted few wide-field (WF) 2D images and corresponding LF measurements of new samples to rapidly tune the pre-trained model adaptive to the new structure (Fig.1d, Supplementary Note 1 and Method). In the adaptive-tuning phase, the WF images of the new samples were readily obtained using the regular port of the inverted microscope and performed deconvolution, serving as the lateral constraint of 3D reconstructions, which were used for calculating the mean-square-errors (MSE) with maximum projection and down-sampling of network’s 3D inferences. To maintain the mapping function from LF to 3D reconstructions and prevent the over-fitting by the lateral constraint, we employed an alternate training strategy to incorporate a small amount of raw data used for the base model into the training process, acting as the volumetric constraint. The lateral and volumetric constraint were alternatively optimized during the fine-tuning phase to together contribute the optimized results in 3D. Through this adaptive-tuning strategy, we fine-tuned the network trained on lysosomes for reconstructing light-field images of fluorescent bead, yielding a resolution of ∼120 nm (Supplementary Fig. 9). Additionally, we successfully transferred the model trained on the outer membrane of mitochondria to the outer membrane of lysosome and mitochondrial matrix, respectively (Supplementary Fig. 9). The finely-tuned Alpha-Net showed significantly reduced artifacts and enhanced fidelity, as compared with VCD-Net and Alpha-Net without fine-tuning yielding sharp but distorted reconstruction.

### Characterization of Alpha-LFM

We demonstrated the performance advances of Alpha-Net through a comparison with end-to-end VCD on both Argolight resolution board and organelle data. To quantitatively illustrate the performance of the network in the process of solving inverse problems, we developed a new network comprehensive performance pyramid (NCPP) method based on the comprehensive evaluation of the fidelity and resolution changes during the network training (Fig. 2a, Supplementary Video 1, Method). We quantified the fidelity and resolution of the reconstruction results during the network’s convergence process by calculating the differences in structural similarity (ΔSSIM with 0 indicating the best similarity) and cut-off frequency (ΔKc with 0 indicating the highest resolution) between the inference results of 196 regions of interest (ROIs) and the ground truths (GTs). The position of data point on the coordinate axes reflects the network fitting ability while the concentration of the distribution represents the robustness of the network across different datasets. Through NCPP metric, we verified that it’s indeed difficult for original VCD to find HiFi, SR solutions through a one-stage supervision, as evidenced by the premature convergence and halting at inferior resolution and fidelity (Fig. 2a, NCPP map, left). In contrast, Alpha-Net under multi-stage data supervision achieved significant improvements in fidelity and resolution right after the initial task optimization (Initial stage, Fig 2b). The progressive optimization strategy further reduced the unnatural high-frequency artifacts caused by such local optimization in the sub-networks. With the decomposed and progressive optimization of Alpha-Net, the network rapidly approached the global optimum (Fig. 2a, NCPP map, right), yielding HiFi and SR reconstructions in the last stage (Last stage, Fig 2b).

**Figure 2.**
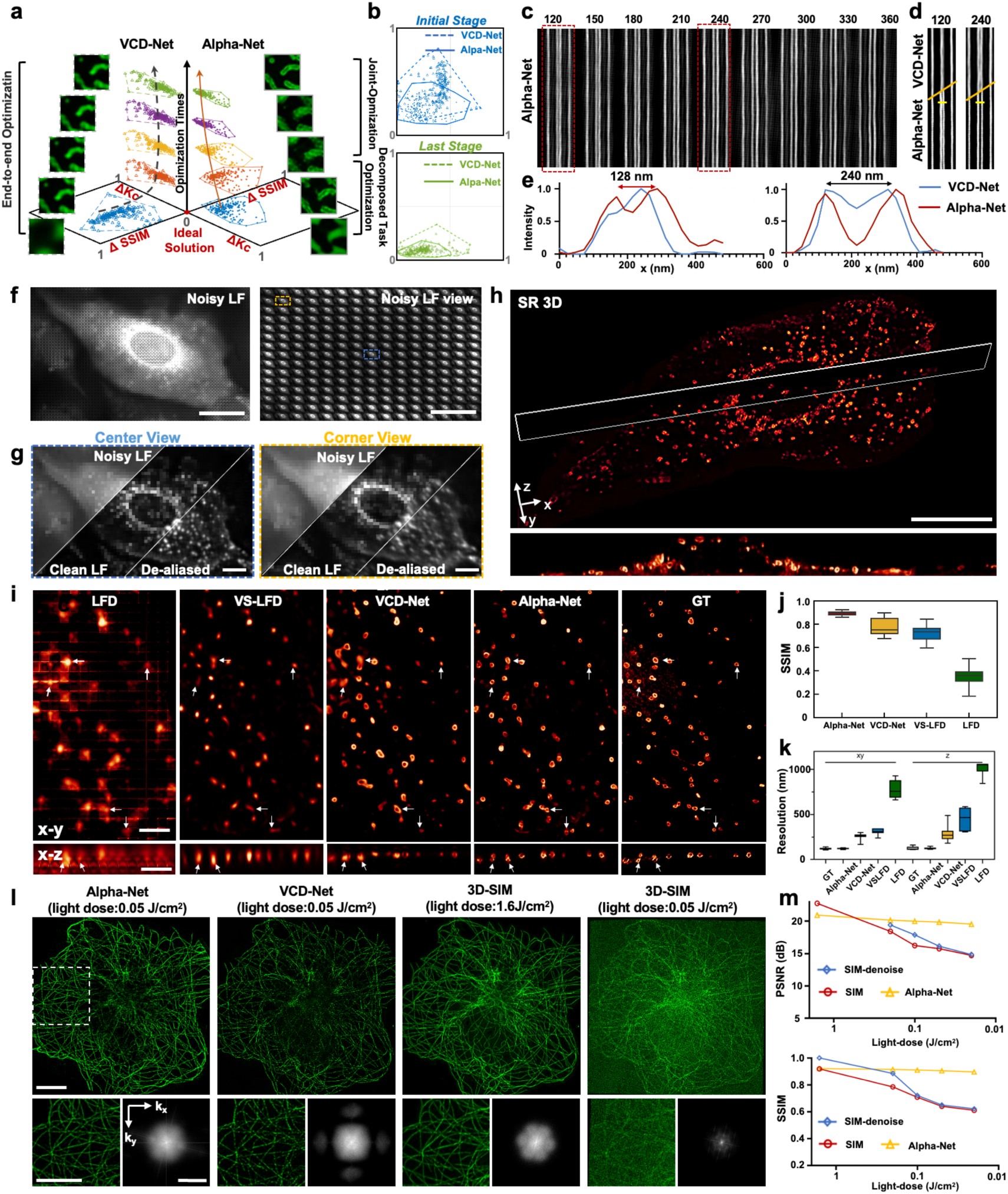
The resolution and structural fidelity of Alpha-LFM. **a**, The comparative NCPP map evaluating the fitting capability of previous end-to-end VCD-Net and Alpha-Net during the optimization process. The fitting ability of the network is comprehensively quantified by the fidelity (ΔSSIM, with 0 indicating the best fidelity) and spatial resolution (ΔKc, with 0 indicating the best resolution) of the 3D reconstructions throughout the entire training process, where ΔKc and ΔSSIM are the difference values of cut-frequency and multiscale structural similarity between the 3D reconstructions and ground truths, respectively. The ideal solution, reflecting a lossless 3D reconstruction, is positioned at (0,0). 196 ROIs are calculated for statistical analysis of K_c_ and SSIM (hollow triangle: VCD, solid circle: Alpha-Net). The shorter distance to the center axis represents the less quality difference as compared with ground truth. b, The 2 charts show the network performance at initial optimization stage (10th epoch) and final stage (well-convergence at 250th epoch). **c**, Resolution characterization of Alpha-LFM using Argolight resolution board featuring adjacent lines with known distances ranging from 120 nm to 360 nm. **d,** Regions of interest (ROIs) showing lines spaced 120 nm and 240 nm apart, imaged using VCD-Net and Alpha-Net. **e**, Intensity profiles along the lines indicated in **d**, quantifying the resolution of Alpha-Net and VCD-Net. **f,** The experimental Noisy LF image of lysosome outer membranes in fixed U2OS cell. Its separated views are shown in the right panel. **g,** The Noisy, Denoised, De-aliased LF views (indicated by dotted boxes in **f**) and **h,** the 3D SR results from three sub-networks of Alpha-Net. Scale bar, 5 μm. **i,** The comparison of reconstructions by Alpha-Net and current leading light-field reconstruction approaches including LFD, VS-LFD and VCD. The white arrows indicate the noticeable errors in LFD, VS-LFD and VCD results whereas being accurately resolved by Alpha-Net. Scale bar, 2 μm. **j,** Structure similarity (SSIM) metric quantitatively comparing the fidelity of Alpha-Net and other approaches with using enhanced Airyscan data (GT) as reference (n=15 volumes). **k,** Decorrelation analysis quantifying the lateral and axial resolution of reconstructions by Alpha-Net, LFD, VS-LFD, VCD and GT. The center line indicates the median, the box limits denote the lower and upper quartiles, and the whiskers represent the minimum and maximum values. **l,** MIPs of microtubules in a fixed COS-7 cell obtained by 3D-SIM under high (1.6 J/cm^2^) and low (0.02 J/cm^2^) light dose, as well as by VCD-LFM and Alpha-LFM under a low light dose (0.05 J/cm^2^) used for volumetric acquisition. The magnified view of ROI indicated by white box and corresponding Fourier spectrums are shown at the bottom. Scale bar, 10 μm, 1/100 nm^-1^. **m,** SSIM and PSNR metrics quantifying the fidelity of 3D-SIM and Alpha-LFM across varying light doses, with using 3D-SIM with AI denoising under high light dose as reference.

Through a simple retrofit of commercial inverted microscope (Olympus IX73) using a designed compact light-field add-on (∼220 mm×140 mm in size, full design in Supplementary Fig. 10 and Method), we conducted LF imaging of the Argolight resolution board, which features adjacent lines with known distances, to quantify our resolution. Alpha-Net successfully resolved adjacent lines with distances ranging from 120 nm and 360 nm (Fig. 2c-e). In contrast, VCD-Net could only resolve lines spaced 240 nm apart and produced ambiguous reconstructions for lines spaced 120 nm apart (Fig. 2d, e). These results validate the capability of Alpha-LFM to achieve a resolution of 120 nm.

Furthermore, we imaged the lysosomes in a fixed U2OS cell and reconstructed their 3D distributions using Alpha-LFM (Fig. 2f-h). The raw LF image of lysosome was effectively denoised, de-aliased and finally transformed into fine 3D structures (Fig. 2g, Supplementary Fig. 11, 12). To validate the resolution enhancement and fidelity by Alpha-LFM, we also obtained the *in-situ* Airyscan^13, 32^ images of lysosomes in the same fixed U2OS cell and enhanced them through an axial-to-lateral isotropic learning (Method). As verified by calculating the structure similarity (SSIM) using enhanced Airyscan microscope’s results as references, the fine structures of lysosomes were accurately reconstructed throughout the 3D volume (Fig. 2i). The fidelity of Alpha-Net reconstruction was quantified to be significantly higher when compared to current leading light-field reconstruction techniques, including LFD, virtually-scanning LFD (VS-LFD)^22^ and VCD (Fig. 2j, Method). Additionally, while LFD, VS-LFD, VCD yielded average lateral resolutions of 780 nm, 315 nm, 261 nm, and average axial resolution of 1021 nm, 440 nm, 277 nm, respectively, Alpha-Net achieved a near isotropic resolution of 120 nm, which is far superior to all the alternative approaches and close to the resolution of GT (Fig. 2k, Method).

We further evaluated the performance of Alpha-Net across different LFM configurations, including various spatial LFM and Fourier LFM systems^18, 21^. To validate its adaptability, we first constructed an LFM system using a microlens array (MLA) with a pitch size of 45.5 µm and a focal length of 1.6 mm (Methods). When reconstructing LF images of lysosomes acquired with this setup, Alpha-Net still demonstrated superior resolution and fidelity compared to state-of-the art light-field reconstruction algorithms, including LFD, VS-Net and VCD-Net (Supplementary Fig. 13). In addition to spatial LFM, we built a Fourier LFM system by incorporating a Fourier lens to perform an optical Fourier transform of the image at the native image plane and placing the MLA (pitch = 3.25 mm, f = 120 mm) at the back focal plane of Fourier lens (Methods). By adapting the data synthesis strategy and view extraction method in the network to align with the wave optics model of Fourier LFM, Alpha-Net accurately reconstructed fine microtubules structures, using 3D-SIM results acquired under high light dose as GT (Fig. 2l, Methods). In contrast, VCD-Net introduced high-frequency artifacts and produced discontinuous signals, resulting in low fidelity.

The high fidelity achieved by Alpha-LFM under low light dose required for volumetric acquisition highlights its advantages in imaging speed and reduced photobleaching. To evaluate these benefits, we assessed the imaging performance of Alpha-LFM across varying total light doses used for volumetric acquisition and compared it to the state-of-the-art live-cell imaging technique, 3D SIM^7, 33^, under identical light dose conditions (Fig. 2l, Method). While 3D-SIM achieves high-quality reconstructions under high light dose conditions, its reconstruction fidelity deteriorates as the light dose decreases. In contrast, Alpha-LFM fully utilizes photons emitted from the entire volume, achieving a higher signal-to-noise ratio (SNR) under the same low light dose. The SSIM and PSNR metrics presented in Fig. 2m confirm the superior and stable reconstruction fidelity of Alpha-LFM compared to SIM and SIM-denoise^34^ under low light dose conditions (Fig. 2m, Supplementary Fig. 14). The combination of high photo-efficiency and robust denoising capabilities enables Alpha-LFM to deliver superior resolution and reconstruction fidelity, making it particularly well-suited for high-speed or long-term imaging applications.

### Comparative performances of Alpha-LFM and other approaches

Alpha-LFM enabled 4D imaging of large-scale mitochondrial dynamics in dozens of cells with a volumetric imaging rate up to 333 Hz and a field of view (FOV) of ∼ 220 × 220 × 10 µm^3^ (Fig. 3a). We compared the reconstruction of Alpha-LFM with those from the alternative LFM reconstruction approaches including VS-LFD ^22^ and VCD^26^ based on end-to-end network model (Fig. 3b, Supplementary Video 2). While Alpha-Net reconstruction clearly showed the outer membranes of mitochondria, these structures remained unresolvable in LFD, VS-LFD and VCD reconstructions, owing to their suboptimal resolutions and noticeable artifacts. Alpha-LFM yielded a near isotropic lateral and axial resolutions of ∼120 nm, which were compared with 230 nm and 370 nm by VS-LFD, 180 nm and 350 nm by VCD, respectively (Fig. 3c). The ultrafast light-field imaging rate together with subcellular-resolution reconstruction thus allowed the visualization of the fast morphology changes of the mitochondria, such as fission and fusion, occurred in three dimensions and at milliseconds time scale (Fig. 3d). Since LFM only requires light exposure once per volume, its photobleaching is notably lower than the scanning-based 3D microscopy (Fig. 3e). Meanwhile, the inclusion of denoising module in Alpha-LFM ensures stable light-field reconstruction from low-exposure, noisy measurements (Supplementary Fig. 15). As a result, low-photobleaching imaging combined with low-exposure reconstruction capability together permitted high spatiotemporal resolution live-cell imaging in long term, yielding over 40000 SR volumes with less than 50% photobleaching (Supplementary Video 3). In contrast, plane-scanning-based light-sheet fluorescence microscopy (LSFM)^34^, 3D SIM^7, 33^ (15 exposures for each plane) and point-scanning-based Airyscan microscopes^13^ suffered from noticeable photobleaching after imaging merely 500, 6 and 50 volumes, respectively (Fig. 3e, f, Method). We compared the resolution and volumetric speed of our Alpha-LFM with current leading 3D fluorescence microscopy techniques for live-cell imaging, including scanning-based SR microscopes such as Airyscan microscope, Instant SIM (iSIM)^8^ and LLSM-SIM^9^ and LFM modalities such as sLFM^19^, Fourier light-field microscopes (FLFM)^21^ and VCD-LFM^26^ (Fig. 3g). While scanning-based SR microscopes exhibit trades-off among spatial resolution and imaging speed, Alpha-LFM has apparently enlarges this limitation, showing spatial resolution as well as volumetric speed far superior to not only SR microscopes, but also other LFM modalities (Fig. 3g).

**Figure 3.**
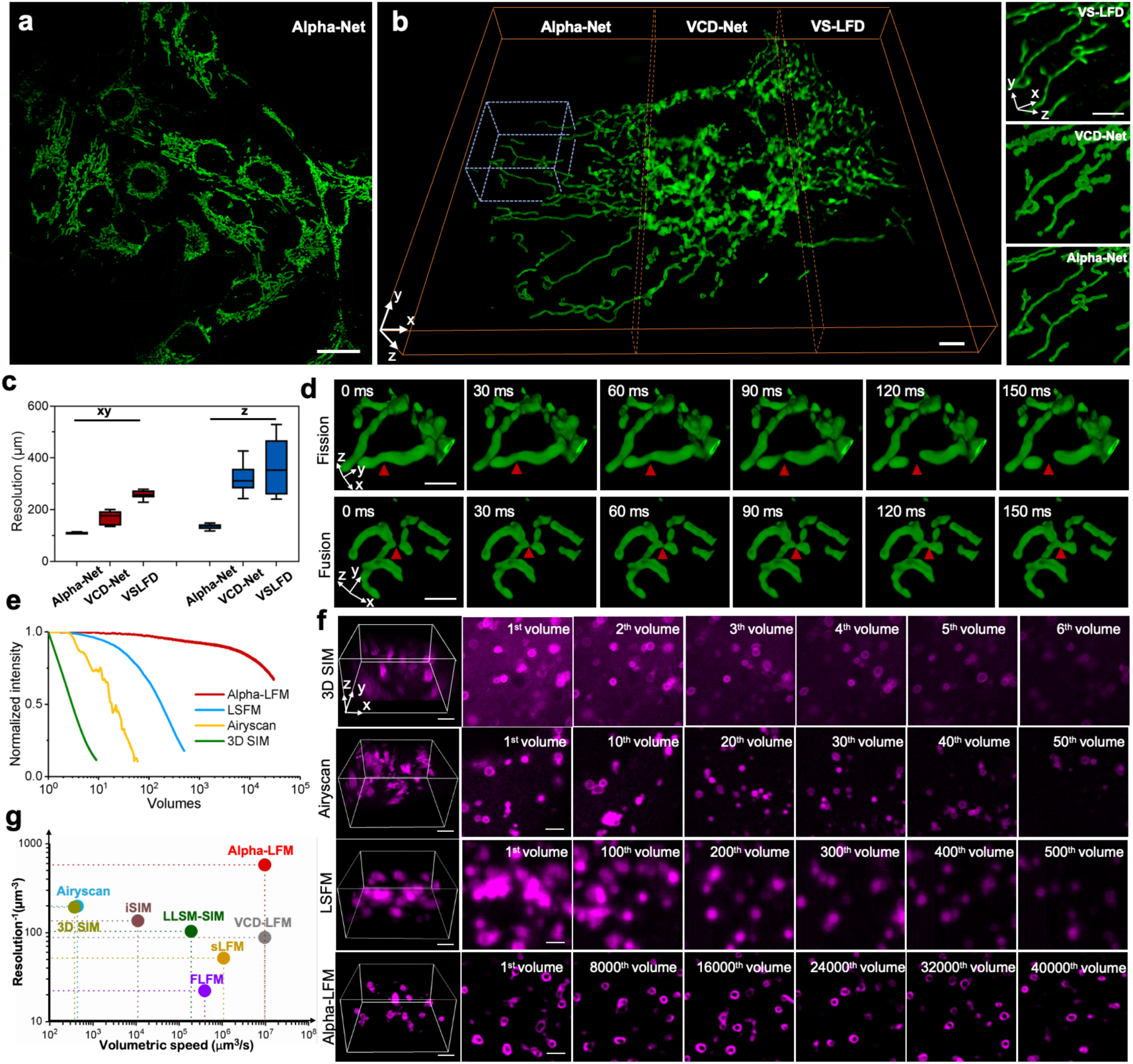
The performance of Alpha-LFM. a, The volume rendering of the large-scale mitochondrial dynamics in dozens of live U2OS cells reconstructed by Alpha-Net within a large FOV of 220×220×10 μm^3^ using a 60×/1.3 NA objective. Scale bar, 20 μm. **b,** The volume rendering of the reconstructed results of mitochondrial outer membranes in live U2OS cells by Alpha-Net, VCD-Net, VS-LFD. Insets show the magnified volume renderings of the ROI indicated by blue dotted box. Scale bar, 5 μm. **c,** Decorrelation analysis quantifying spatial and axial resolution of VS-LFD, VCD-Net and Alpha-Net. n= 20 volumes were analyzed at both xy and xz planes. The center line represents the median, the box limits represent the lower and upper quartiles, and the whiskers represent the min and max value. **d,** Time-lapse 3D visualization captures the rapid morphological transformations of the mitochondrial outer membrane occurred at milliseconds time scale, illustrating both the processes of mitochondrial fission and fusion. Scale bar, 2 μm. **e,** Comparisons of the photobleaching rates between Alpha-LFM, LSFM, Airyscan and 3D SIM. **f,** Max intensity projection of images of lysosomes in live U2OS cells imaged via 3D SIM (30 s every volume for a whole cell), Airyscan (4 minutes every volume for a whole cell), LSFM (4 seconds every volume for a whole cell) and Alpha-LFM (3 ms every volume for at least a whole cell). Scale bar, 2 μm. **g,** The comparisons of the performance in resolution and volumetric imaging speed between Airyscan, iSIM, LLSM-SIM, FLFM, sLFM, VCD-LFM, and Alpha-LFM.

### High-speed 3D imaging of peroxisomes and ER enabled by Alpha-LFM

The ultrafast volumetric imaging rate combined with subcellular-resolution reconstruction provided by Alpha-LFM enabled the visualization of rapid dynamics of peroxisomes and the endoplasmic reticulum (ER) in three dimensions.

Without requiring scanning, Alpha-LFM successfully demonstrated its capability to capture peroxisomes (tagged with SKL-mApple) in live U2OS cells at 100 volumes per second (vps) over a 1-minute duration, yielding 6000 volumes (Fig. 4a, Supplementary Video 4). This high imaging rate allowed the extraction of spatiotemporal patterns of peroxisome motion in 3D on a millisecond timescale (Fig. 4b). In contrast, current scanning-based 3D microscopy techniques^9, 35^ achieve imaging speeds up to 10 vps. To evaluate the impact of imaging speed on peroxisome analysis, the 100-vps data was downsampled to 10 vps, and peroxisome tracking was performed on both datasets. While analysis of the 10-vps data revealed a velocity limited to 1μm/s, the 100-vps data captured velocities as high as 10 μm/s (Fig. 4c, d). This discrepancy arose from missed movements and reduced trajectory accuracy caused by the lower imaging speed (Fig. 4e, f).

**Figure 4.**
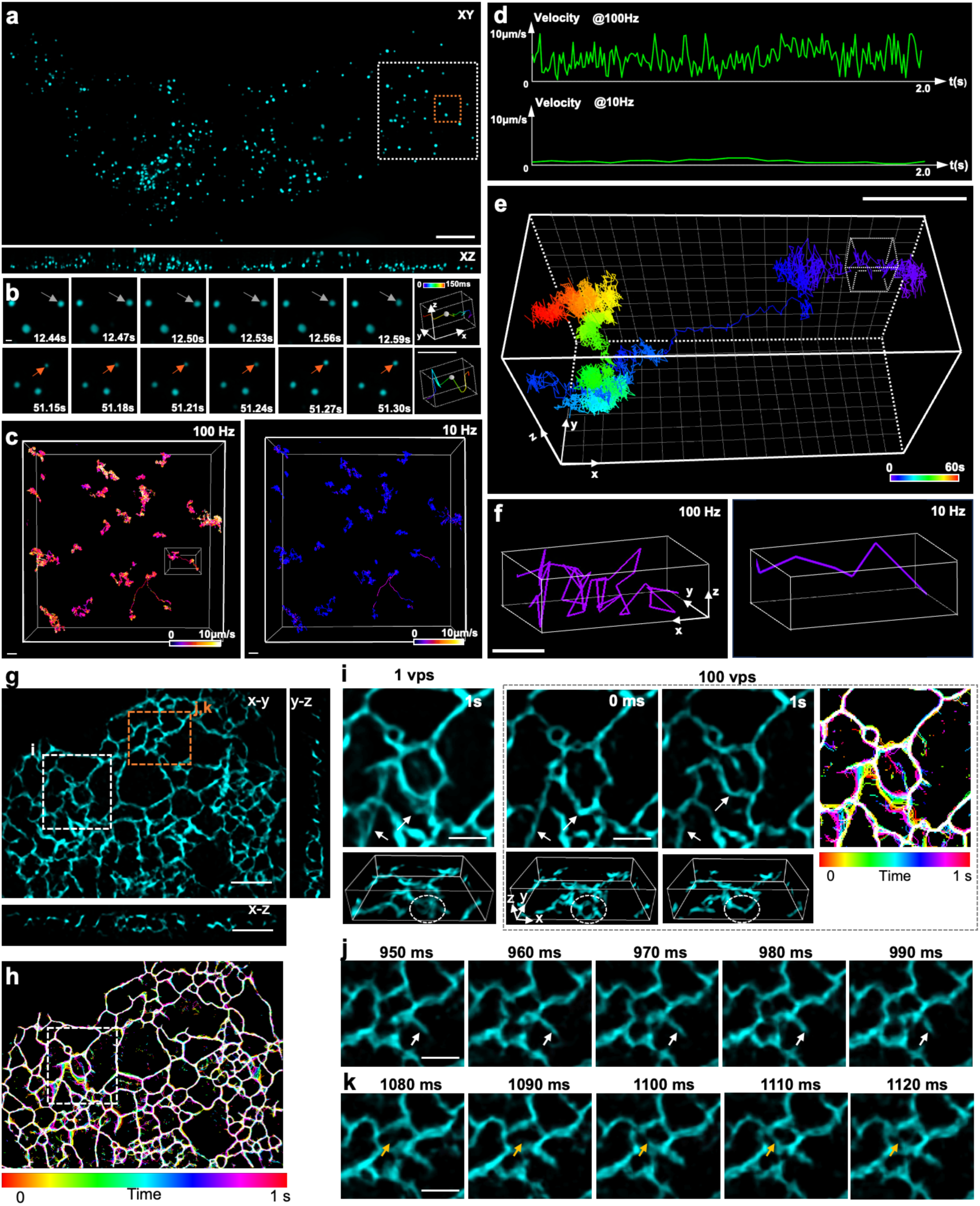
Imaging of peroxisomes and ER in live cells at a 100 volumes per second (vps) using Alpha-LFM. a, Volume rendering of peroxisomes (tagged with SKL-mApple) in a live U2OS cell, acquired with Alpha-LFM. **b,** Time-lapse MIPs of the ROI indicated by orange boxes in a. Arrows indicate the rapid motion of peroxisomes. **c,** 3D tracking of peroxisomes imaged at 100 vps and those down-sampled to 10 vps, with velocity encoded by color. **d,** Velocity plots of the peroxisome highlighted by a box in **c**. **e,** Trajectory of the peroxisome highlighted by box in **c**. **f,** Comparison of trajectories tracked using the 100- vps and 10-vps results. **g,** x-y MIP and x-z, y-z slices of the ER (tagged with Sec61β-EGFP) in a live COS- 7 cell captured by Alpha-LFM. **h,** Projection of skeletonized images over 1 s, with time encoded by color, visualizing ER dynamics. Scale bar, 5 μm. **i,** Comparison of ER dynamics of the ROI (marked by white boxes in **g** and **h)** acquired at 100 vps versus 1 vps (downsampled). White arrows and circles highlight dynamics resolved at 100 vps but blurred at 1vps. Scale bar, 2 μm.. **j, k,** Magnified time-lapse images of the ROI marked by orange box in **g**. White arrows in **j** indicate the stretching of an ER tubule within 40 ms, while orange arrows in **k** show the formation of a new tubule within 40 ms. Scale bar, 2 μm.

Additionally, we recorded the dynamics of the ER (tagged with Sec61β-EGFP) in a live COS- 7 cell at 100 vps using Alpha-LFM (Fig. 4g, Supplementary Video 5). The rapid growth and remodeling of the ER, with velocities reaching ∼3.5 μm/s, have previously been observed in 2D using 2D SIM microscopes^8, 25, 36^. Alpha-LFM’s volumetric imaging capability at high resolution provided a clear visualization of ER tubules in 3D. The 100-vps speed allowed us to resolve the remodeling of individual ER tubules within milliseconds, which appeared blurred at 1 Hz, the maximum volumetric imaging rate achievable by current 3D SR microscopes^12, 25^ (Fig. 4h, i). Leveraging this high speed, we observed the stretching of an ER tubule and the formation of a new ER tubule, both occurring in less than 50 ms (Fig. 4j, k). These experiments underscore the essential of Alpha-LFM in capturing highly dynamic biological processes.

### Dual-color Alpha-LFM for 5D *in-toto* imaging and quantification of lysosome-mitochondria interactions

The high spatiotemporal resolution of Alpha-LFM allowed the visualization of rapid morphological changes in mitochondria, such as fission and fusion, in three dimensions. We validated the fidelity of dynamic Alpha-LFM imaging for identifying mitochondrial activities using *in-situ* WF images (Supplementary Fig. 16) and co-expression of Drp1 (Supplementary Fig. 17). Notably, we observed 48 mitochondrial fission events, with ∼96% of these events (n=46) were marked by Drp1, consistent with previous studies^37^. These results confirm that the fission events identified by Alpha-LFM are genuine.

Using Alpha-LFM, we conducted simultaneous 5D (3D space + time + spectrum) SR imaging of the outer membranes of lysosomes (tagged with Rab7-mCherry2) and mitochondria (tagged with Tomm20-EGFP) in live U2OS cells (Fig. 5a, Supplementary Video 6). The isotropic subcellular resolution presented in five dimensions then permitted *in-toto* visualization of the Lysosome-Mitochondria (Lyso-Mito) interactions (Fig. 5b, c). While 2D microscopes merely provided projection images of Lyso-Mito contacts, our ability to reconstruct 3D processes eliminated false judgments and yielded more accurate analytic results (Fig. 5d). By measuring the distance between lysosome and mitochondria in 2D and 3D (Method), respectively, we identified a 24 % margin of inaccuracy in 2D results (Fig. 5e, 11 false cases in 41 events from 17 cells), proving the significance of Alpha-LFM for investigating organelle interactions in 4D. Recent research has revealed that lysosome-mitochondria contact sites may serve as a markers for mitochondrial fission^38^. We scrutinized 35 fission cases from 17 cells reconstructed by Alpha-Net to measure the proportion of Lyso-Mito contact in the mitochondrial fission events. We found that lysosomes contacted mitochondria at 48 % of mitochondrial fission sites, which was significantly lower than the rate obtained via 2D analysis of our own images (71%) (Fig. 5f). This was presumably due to the tendency of 2D imaging methods to misidentify the Lyso-Mito contact. We also analyzed the Lyso-Mito contacts at mitochondrial fusion sites and the proportion was 41% in our statistics. With the high spatiotemporal resolution of Alpha-LFM, we also perform 3D tracking of the endpoints of mitochondrial fission and fusion and analyzed whether Lyso-Mito contact would affect the speed of mitochondrial fission and fusion (Fig. 5g, Supplementary Video 7). While there is no significant difference in fission speed was found when 2D analysis was performed (P>0.05, one-way ANOVA), the results based on our 3D images showed a positive correlation between lysosome contacts and fission (P<0.05) / fusion velocity (P<0.001) (Fig. 5h-i). These findings suggest that interactions with lysosomes might accelerate mitochondrial fission and fusion. However, this hypothesis requires further validation through systematic physiological and biochemical experiments.

**Figure 5.**
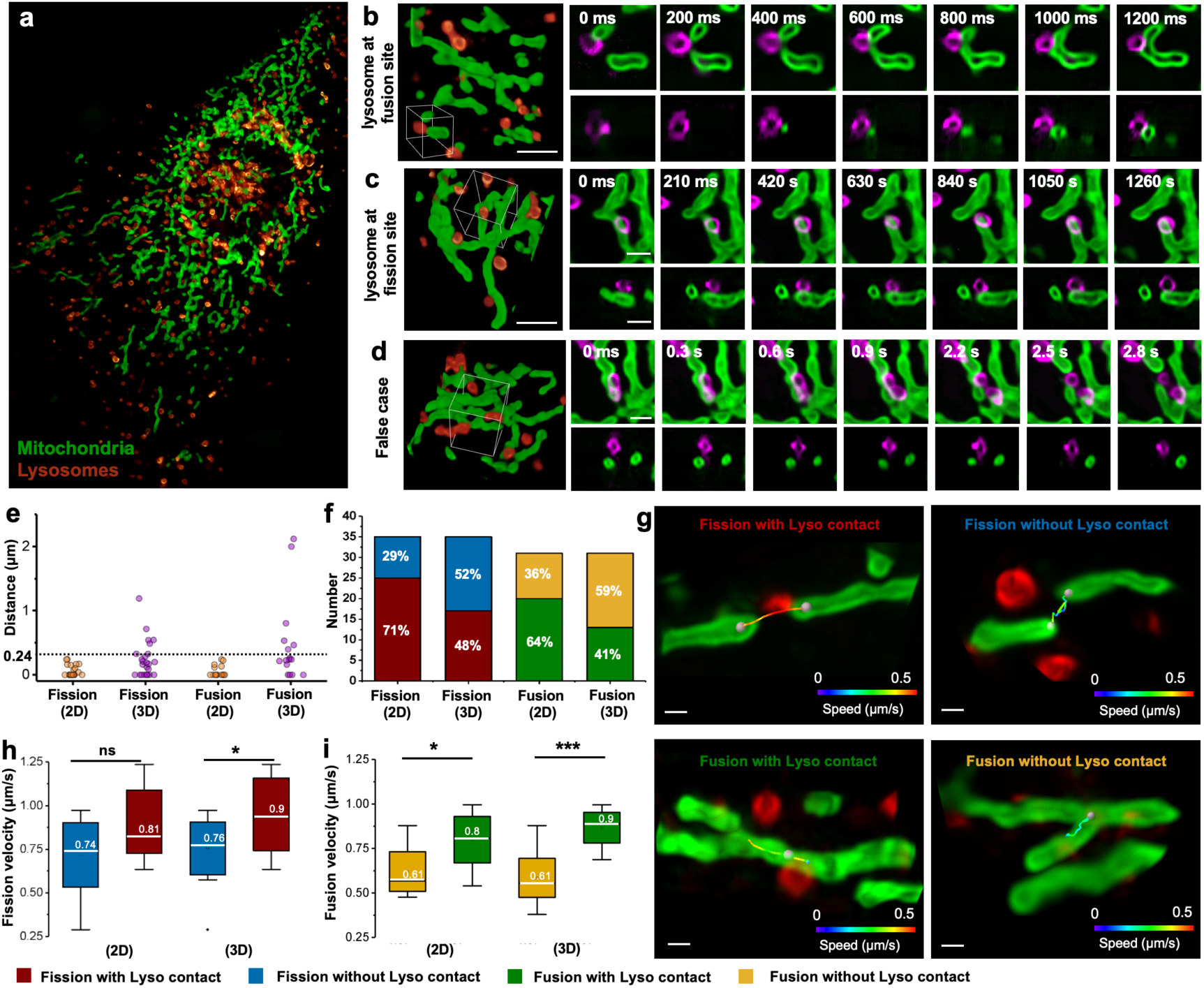
Alpha-LFM enabling 5D *in-toto* imaging and quantitative analysis of mitochondria-lysosomes interactions at high spatiotemporal resolution. a, A dual-color volume rendering of the outer membranes of mitochondria (tagged with Tomm20-EGFP) and lysosomes (tagged with Rab7-mCherry2) in a live U2OS cell. Scale bar, 10 μm. **b, c,** Time-lapse images of typical lysosomes-mediated mitochondrial fusion and fission events, respectively. Volume rendering shows the first volume of the events in 3D. The xy and xz slices of the ROI indicated by white boxes in the volume rendering accurately show the rapid interactions between the lysosome and the fission / fusion sites of mitochondrial. **d,** An example of a false positive case where the lysosome appears to be in contact with the fission sites in 2D projection, but in fact there is a significant distance between them in 3D observation. **e,** The 3D distances between the lysosomes and mitochondrial contact sites reveal the inaccuracies in the identification of Lyso-Mito contacts in 2D. The dashed line indicates the close proximity (<240 nm) between lysosomes and mitochondrial constriction sites that can be identified as Lyso-Mito contacts. **f,** The proportion of mitochondrial fission and fusion events occurred with and without Lyso-Mito contacts identified in 3D and 2D projection, respectively. The results of 3D indicate lower contact rate during the mitochondrial fission and fusion than the rate obtained via 2D measurements (n= 35 fissions and 31 fusions from 17 cells). **g,** 3D tracking of mitochondrial fission and fusion with and without Lyso-Mito contact. The fission and fusion velocity are represented by color. The hotter color when in contact with lysosome indicate the higher fission and fusion velocity compared to without lysosome’s contact. The center line represents the median, the box limits represent the lower and upper quartiles, and the whiskers represent the min and max value. Scale bar, 500 nm. **h, i,** The velocity of mitochondrial fission and fusion with or without Lyso-Mito contact via 2D and 3D analysis. The velocity is significantly higher when in contact with lysosomes via 3D analysis while there is no significant difference of fission velocity based on 2D results. Data are shown as mean ± s.d. ns P >0.05, *P<0.05, ***P < 0.001 (One-Way ANOVA).

### Long-term imaging and quantitative analysis of mitochondrial fates across cell cycles

The low phototoxicity and robust denoising capability of Alpha-LFM enable 3D imaging of mitochondria evolution (tagged with Cox4-EGFP) in live *U2OS* cells across a long timescale up to 60 hours. Considering both rapid mitochondrial dynamics and their long-term fates need to be studied, we specifically designed an automatic imaging strategy to track the high-speed fission and fusion process throughout the entire cell cycle with minimal phototoxicity (Fig. 6a, Method). During each cycle of 4 hours, we continuously imaged the mitochondria for 30 minutes across 8 FOVs, with exposure time of 500 ms and a temporal resolution of 10 seconds (Fig. 6b). During this 30-min mid-term observation window, the imaging pipeline included the following steps: (i) Implementing an Alpha-Net-based focusing strategy to correct sample drift along z axis. Mitochondrial were imaged and quickly reconstructed by Alpha-Net to calculate the z-drift distance, which would be further corrected by moving the objective. This process typically takes merely 1 second. (ii) Imaging the mitochondria dynamics at fixed depth of interest for 30 minutes, with an exposure time of 500 ms and interval time of 10 seconds. Meanwhile, we imaged *in-situ* chromosomes of the same FOV for identifying the related cell stage with an exposure time of 500 ms and interval time of 100 seconds (Supplementary Video 8). After 36-hours observation containing 9 cycles of 30-minture continuous recording, we discovered diverse viabilities of the cells within one batch, identifying both inertia without cell division and active 2-generation divisions across one entire cell cycles (Fig. 6c).

**Figure 6.**
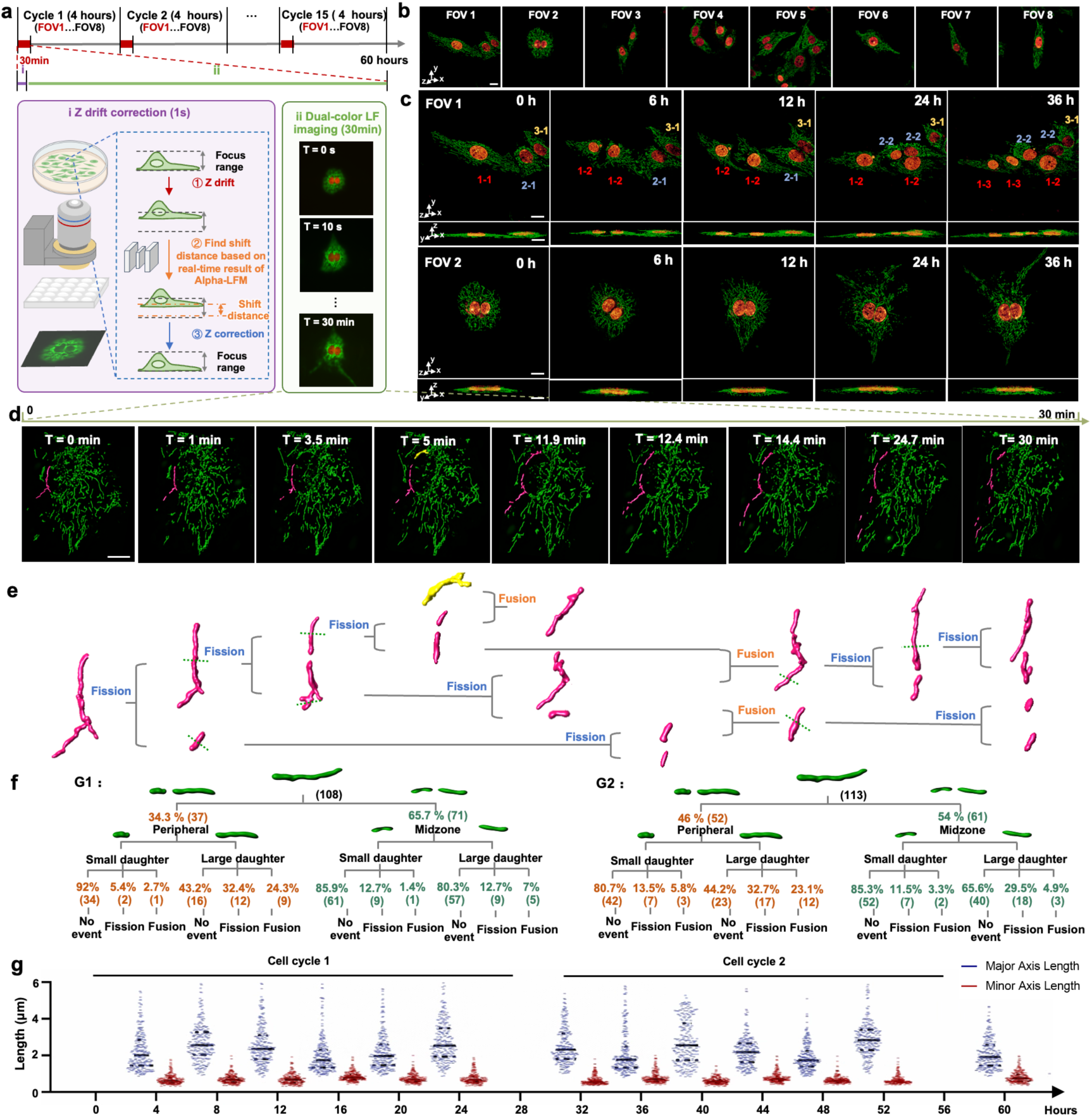
Long-term imaging of mitochondrial fates and morphology changes across cell cycles using Alpha-LFM. a, Automatic Alpha-LFM imaging pipeline designed for the long-term imaging (up to 60 hours) of mitochondria (tagged with Cox4-EGFP) and chromosomes (tagged with H2B-mCherry). In each cycle of 4 hours, the cells were continuously imaged for 30 minutes. The 30-min observation majorly includes the following steps: (i) An instant VCD-based auto-focusing strategy to correct sample drift along the z axis. Quick light-field imaging and reconstruction of mitochondria were performed to calibrate the z- axis drift of the system. The drift was then corrected by moving the objective using piezo scanner. (ii) Alpha-LFM imaging of the mitochondria dynamics and chromosomes in cell nuclei for 30 minutes. 8 FOVs were sequentially imaged in each cycle with an interval time of 10 seconds for mitochondria and 100 seconds for chromosomes and an exposure time of 500 ms for both. **b,** The MIPs of 8 FOV imaged in the first cycle. **c,** The volume renderings of time-lapse images visualizing diverse cell viabilities with one undergoing 2-generation divisions (the first FOV in b) and another showing inertia without cell division (the second FOV in b) during the same 60-hours imaging. The inset numbers indicate three cells and their later generations. **d,** The 3D visualization of a representative mitochondrion during 30 minutes. The mitochondria are highlighted in the 3D rendering of whole cell. **e,** Detailed lineage tracing of the selected single mitochondrion with 7 generations traced. The mitochondria daughters rendered by magenta were from the same source mitochondrion (leftmost one) while the yellow rendering was a foreign mitochondrion. **f,** Schema depicting the different fates of the daughter mitochondria in different stages of interphase G1 and G2 in the cell cycle. Mitochondria were tracked for 30 minutes. Both the quantities and portions of the specific mitochondria events were calculated. **g,** The morphology changes (Major axis length and minor axis length) of mitochondria during two entire cell cycles throughout 60 hours. The data are shown as mean with standard deviation (SD), and the spots indicates all calculated values. Scale bar, 10 μm.

Smart Alpha-LFM imaging of live cell as such included capture of instantaneous mitochondrial fission / fusion events (500 ms), lineage tracing of single mitochondrion’s change in middle term (30 minutes), and evolution mapping of all mitochondrial in long term (60 hours), thereby enabling *in-toto* investigation of the mitochondrial fates during entire cell cycle. For example, up to seven generations of the fissions / fusions of an individual mitochondrion were recorded during the 2^nd^ 30-minute observation. Each fissions / fusions activities of the selected mitochondrion and their resulting later generations were successfully visualized in three dimensions (Fig. 6d). We casted lineage tracing of these mitochondrion activities (Fig. 6e, Supplementary Video 9). In this case, we observed that majority of subsequent generations of the source mitochondrion (the Magenta ones in Fig. 6e) fused with their relatives with only one exception being fused with a foreign mitochondrion (the yellow one in Fig. 6e). Furthermore, we followed the fates of the daughter mitochondria from peripheral and midzone fissions at whole-cell scale. After analyzing 108 / 113 mitochondrial at G1 / G2 stages, we validated that most of the small peripheral daughter mitochondria were excluded from further fusions or divisions, consistent with previous study^39^. Also, we observed a significant difference in the rates of no event occurring in these small peripheral daughters between the G1 (92%) and G2 (80.7%) phases (Fig. 6f, Supplementary Fig. 18). This is probably because the G2 phase is the phase before the cell division, in which the mitochondria are more active to prepare for the cell division^40^. Furthermore, Alpha-LFM results also allowed us to study the morphology changes of mitochondria, including variations in major and minor axis length, throughout two entire cell cycles (Fig. 6g, Supplementary Fig. 19). The significant variations in mitochondria morphology and event rates highlight the importance of long-term imaging facilitated by Alpha-LFM in advancing biological research and applications.

## Discussion

Sustained observation of the subcellular biological dynamics at their physiological status is essential to investigating diverse organelle functions and their interactions in cell biology. This is difficult for either scanning-based SR microscopes with suboptimal speed and phototoxicity, or high-speed light-field microscopes with limited single-cell resolution. Alpha-LFM circumvents this compromise between speed, resolution and photon efficiency based on the new development of deep-learning VCD pipeline now with efficient solution seeking and strong generalization abilities. We’d like to summarize the following technical advances of Alpha-LFM enabled by our new developments of physics-aware network framework and adaptive tuning strategy. First, Alpha-LFM includes new physics-aware model design, hierarchical data synthesis procedure and DPO strategy to maximize the network’s solving ability to light-field inversion problem with huge solution space, thereby efficiently pushing the resolution of LFM beyond diffraction limit. Meanwhile, the strong denoising by the network in conjunction with photon-efficient LFM modality together enables ultra long-term cell observation under physiological status, wiping off the phototoxicity issue existed commonly in scanning-based SRM with allowing over 10-fold more measurements. We also demonstrated the broad adaptability of the ALPHA procedure to various LFM configurations, including spatial LFM excelling in imaging large fields of view and Fourier LFM being well-suited for resolving denser signals. Furthermore, the intrinsically weak generalization ability of a supervised network is also improved in Alpha-LFM by using readily-accessible WF and LF measurements of live sample to instantly tune the model adaptive to new types of signals. It is worth noting that Alpha-LFM has been wrapped into a fully opensource program with a user-friendly GUI provided. With Alpha-LFM, we four-dimensionally imaged rapid motion of peroxisome and the endoplasmic reticulum at 100 vps and highly dynamic interactions between lysosomes and mitochondrial membranes at isotropic 120-nm spatial resolution. The high quality We also demonstrated SR fluorescent imaging of live cells with probably longest observation time of 60 hours and up to 40000 volumes recorded. The image results support lineage tracing of both short-term changes and long-term fates of mitochondria for the first time at single-mitochondrion resolution and across an entire cell cycle.

While Alpha-LFM demonstrates improved performance compared to existing LFM approaches, it still faces inherent limitations stemming from supervised deep-leaning approaches. Although the adaptive-tuning strategy enhances the generalization ability of the supervised network, it is envisaged that the incorporation of more data, especially those with higher resolution and volumetric prior, will further improve the performance of fine-tuning. Meanwhile, it’s noted that Alpha-LFM still requires a considerable amount of high-resolution label data to initiate the training of base model. Additionally, these deep-learning-based super-resolution method cannot guarantee 100% fidelity, particularly in regions with highly-dense signals (Supplementary Fig. 14).

We anticipate that the generalization issue and dependency on external data will be circumvented by a full-cycle self-supervision reconstruction strategies, in which the *in-situ* WF measurements of the samples can be used as internal training labels to further improve the resolution of LF views and lead to reconstruction with higher quality. A completely unsupervised deep-learning light-field reconstruction could be also made possible by using implicit network representation (INR)^41, 42^. It’s promising because the intrinsic view synthesis capabilities in INR is indeed helpful to the enhancement of axial resolution and reduction of reconstruction artifacts. But the learning-and-representing mode for each single LF image is very time consuming and computationally demanding. INR-based reconstruction needs to solve this efficiency problem before it becomes as practical as DL and deconvolution approaches. Given that a lot of *in vivo* biological dynamics occur in the deep tissues, we also expect the combination of Alpha-LFM with more advanced LFM techniques, e.g., LFM with AO or two photon excitations, or even a system working at NIR-II window, to *in-toto* observe the subcellular dynamics inside living animals. Taken together, we believe Alpha-LFM has strongly pushed the spatiotemporal limit of current SRM / 3D fluorescence microscopy and could be a powerful and accessible tool that helps a broad range of biology research to demystify the worlds inside the cell. The paradigm shift it shows could be also beneficial to propelling other microscopy techniques towards deeper, faster and clearer imaging.

## Methods

### LFM optical setup

An add-on device was designed to provide easy light-field imaging of live cells on a commercial inverted fluorescent microscope (Olympus IX73), thus enabling wider applications for biology researchers without optics background. The device only requires the assembling of off-the-shelf lenses into a customized mounting base without any complex alignment of the optical path. To precisely position MLA at the native image plane of the microscope, a customized lens tube 1 was designed to fix the MLA (RPC Photonics, MLA-S100-f28) at the Flange focal distance away from the camera port. Then a pair of relay lens (Thorlabs, TTL100-A) was integrated into another customized lens tube 2 to 1:1 relay the back focal plane of MLA (light-field image plane) onto the camera sensor. A high-precision zoom housing (SM1ZM, Thorlabs) was used to interconnect the 2 lens tubes and allow fine focusing of the light-field image plane (Supplementary Fig. 10). The complete add-on device containing the two lens tubes and the zoom housing was then mounted between the microscope’s camera port and the sCMOS camera (Photometrics Prime BSI Express), converting the ordinary inverted microscope into an advanced light-field microscope. This setup contains 15×15 views (Nnum=15). An alternative LFM configuration was implemented by replacing MLA with one featuring a pitch size of 45.5 µm and a focal length of 1.6 mm, containing 7×7 views (Nnum=7) (Supplementary Fig. S13).

Additionally, a Fourier LFM system was constructed by adding a Fourier lens (Thorlabs, AC508-250-A) to perform Fourier transform of the native image plane, placing microlens array (pitch = 3.25 mm, f = 120 mm) at the back focal plane of Fourier lens, and using a 100×/NA1.5 objective lens (Olympus UPL, APO100XOHR). This setup was used for imaging microtubules and ER in Fig. 2, 4 and Supplementary Fig. 14.

### Training data processing pipeline

A data processing pipeline was developed to generate all training data (including Noisy LF, Clean LF, De-aliased LF and 3D SR) from the 3D SR data. It’s noteworthy that the 3D SR data could be acquired from any SR microscopy with a resolution of around 120 nm. In our study, we provide an easily accessible solution that acquiring from commercial Airyscan confocal microscopy^13, 32^ (Zeiss, LSM900) by imaging fixed cell and enhanced through applying a well-established self-learning network^12, 24, 43^, thereby achieving an isotropic resolution of 120 nm. We also demonstrate that our strategy can perform well on raw Airyscan data (Supplementary Fig. 20), 3D SIM data acquired from commercial SIM microscope (HIS-SIM, Guangzhou Computational Super-resolution Biotech) (Fig. 2l, Supplementary Fig. 14) or SR data acquired from 4-beam SIM^12^ (Supplementary Fig. 21). A SAS LFP pipeline was designed to create De-aliased LF images to guide the LF de-aliasing task. The 3D SR volumes were shifted by a distance less than the size of the microlens along the horizontal (both x and y respectively) and then were projected by convolving with 5D PSF of LFM to yield multiple light field projections that included more spatial information through multiple sampling. The shifted times depended on the under-sampling rate of the microlens, which is 5×5 times with a step of 3 pixels for the setup of Nnum = 15 and 3×3 times with a step of 2 for the setup of Nnum =7 in this paper. The SAS LF projections were then realigned to generate the De-aliased LF images according to the arrangement of the light.

Then, Clean LF images (the center ones with shift = 0 of the De-aliased LF images) were used to guide the denoising task. To generate Noisy LF with accurate SNR matched with the experimental LF, the Clean LF images was normalized to the same range of experimental LFs and various noises in the range of the highest and lowest noise in the experimental LFs during long-term observation were added to the Clean LFs to generate Noisy LFs. The SNR was calculated by:SNR 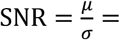 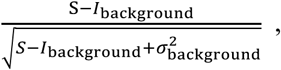 where S is the average signal intensity value in the image, *I_background_* and *σ_background_* are the mean and the standard deviation of the background, respectively. Finally, we obtained De-aliased LF images, Clean LF images, Noisy LF images, which were all conveniently generated from the same 3D SR images, for the network training of Alpha-LFM.

For Fourier LFM, the LF projections were generated by using the PSF of FLFM. The high-resolution (HR) LF images used to guide the LF de-aliasing network were generated by squaring the PSF of Fourier LFM and convolving it with 3D SR data.

### The network design of Alpha-LFM

Previous deep-learning-based light-field reconstruction network enabled resolution enhancement with training the light-field projections and high-resolution ground truths in an end-to-end manner. This one-step network produces hallucination or artifacts when solving hard inverse problem. Similar with the physical process of light-field image generation, our Alpha-Net disentangles this inverse problem into four sub-problems: denoising, de-aliasing, 3D reconstruction. The three sub-problems are divided and conquered by jointly optimizing task-specific networks with weighted objective function.

The architecture of the denoising sub-network was based on attention mechanism^44, 45^, which consisted of dual branches (channel-attention and view-attention) to extract the view-wise features from input views with different SNR. The denoising module can be formulated as

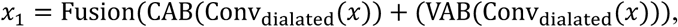

where Conv_dialated_ means dilated convolution operation that enlarges receptive field. Fusion is feature element-wise adding. CAB and VAB are the channel-attention branch and view-attention branch. In order to suppress the noise fluctuation, L1 and L2 loss is used as loss function during denoising stage:

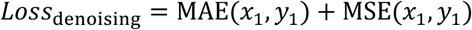

For LF de-aliasing, how to effectively utilize the angular and spatial information encoded in LFs poses a challenge when using conventional 2D super-resolution networks, such as ResNet^46^ and RCAN^47^, directly on the extracted views. These approaches often fail to fully leverage the rich angular features inherent in LFs. In FLFM, we used the dual-attention network structure as denoising sub-network to extract angular features from FLFM views, taking advantage of the distinct 4D representation in the Fourier domain^48^. To increase the sampling rate of LFs, an additional upsampling layer was integrated at the end of the dual-attention network, forming the FLFM de-aliasing network. In spatial LFs de-aliasing, considering the interleaved arrangement of angular and spatial information in spatial light-field images, we employed two types of dilated convolution operations to disentangle 2D spatial information and 2D angular information^49^. These two convolution and series of activation function and feature fusion layers form Spatial-Angular block (SAB). According to sequential connection of four SAB and upsampling convolution, the aliased spatial information caused by sparse sampling rate of microlens array can be de-aliased:

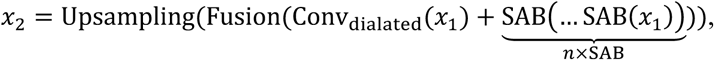

where Fusion is channel concatenate operation. Considering the potential parallax information across different views, epipolar constraint (EPI_consistency_) on de-aliasing prediction is proposed to keep geometry relationship among various views. The objective function of de-aliasing can be formulated as:

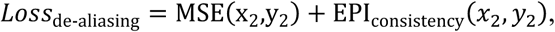

where MSE means mean squared error between network output and de-aliased LF.

For 3D reconstruction model, we extended original VCD to handle super-resolution reconstruction. According to adding MultiRes blocks^50^ and modifying activation function, the capability of VCD network is enhanced achieving lower fitting errors when solving SR reconstruction:

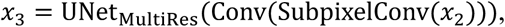

where SubpixelConv refers to the subpixel convolution operation of the input.

Correspondingly, for 3D reconstruction task, we adopt MSE loss and lateral gradients loss to preserve the structure information of ground truth.

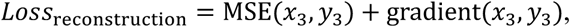

These three networks are optimized synchronously based on designed weighted objective function:

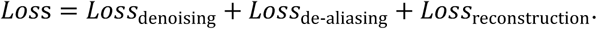

For Alpha-Net training, all models were trained on NVIDIA GeForce RTX 3090 or 4090 with Python version 3.8 and Tensorflow version 1.15.0. The training process of a Alpha-Net model was composed of two stages. First, pretrain 3 sub-networks, including denoising network, de-aliasing network and 3D reconstruction network. Specifically, denoising network was trained with a learning rate of 1×10^-4^ and a training epoch of 51. For de-aliasing network and 3D reconstruction network, they used the same learning rate (5×10^-4^) but different training epoch (51 and 151, respectively). The pretraining process lasted for 16 hours. Secondly, train Alpha-Net with these pretrained model under the designed weighted loss. During this decomposed-progressive optimization, we adopted step-based learning rate schedules with a decay factor of 0.5 and a decay step of 25 to enhance model performance. The initial learning rate was set to 1×10^-4^ and the total training epoch was 101. The training time of second phase was roughly 2 hours.

Once the model was trained, the captured LFs could be reconstructed to 3D volumes through network inference. The inference time was determined by the computational capacity of the device and the voxel count of the reconstructed volume, for example, reconstructing a 3D volume with a size of 2040×2040×161 (height×width×depth) from captured LF (1020×1020, height×width) took ∼0.540 s (∼0.062 s for LF denoising, ∼0.080 s for LF de-aliasing and ∼0.398 s for 3D Reconstruction). For more details about Alpha-Net implementations, see Supplementary Note 1 and our open-source code.

### Adaptive-tuning strategy for reconstruction of unseen samples

Unlike previous supervised learning strategy^26^ only solving the inverse problem for specific samples similar to the training data, Alpha-LFM adopted an adaptive-tuning strategy to overcome the hallucination and abnormal structure features when facing unseen data for trained model. We developed an adaptive-tuning strategy to reconstruct HiFi 3D signals from LF captures of unseen types of structure that not included in training data via the fine-tuning model, in which only an *in-situ* 2D WF image captured together with the LF imaging was required instead of the supervision of 3D label data from another scanning microscope. During the adaptive-tuning phase, we adopted an alternative optimization approach to re-update parameters of the trained model. The paired synthetic training data (e.g. the outer membrane of mitochondria) containing 3D stack were used to provide the volumetric priors to ensure the fidelity of 3D prediction of original model while the experimental LF captures and corresponding WF images act as prior knowledge of new domain (e.g. the outer membrane of lysosome) to prevent data bias brought by trained model. For synthetic data training, the designed weighted loss of original model was adopted:

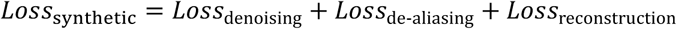

For the new domain transfer, reprojection loss was to provide constraints on the different structure of unseen samples:

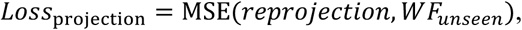

where *reprojection* was computed from the maximum projection and down-sampling of network 3D prediction of *LF_unseen_*. The WF images were processed by deconvolution algorithm to suppress out-of-focus signals. To ensure the accuracy of this loss computation, the two detection modalities (LF and WF) were pre-aligned with the aid of fluorescent beads to yield paired LF images and registered WF images. The paired LF-WF dataset in fine-tuning phase contained ∼3 cells. In this work, *Loss_synthetic_* and *Loss_projection_* were calculated every two consecutive iterations N and N+1, respectively. The training time of fine-tuning phase was typically 70∼100 folds less than the consumption of building a new model, with a total mini-batch iteration of ∼30,000 finished 10-15 minutes.

### Network comprehensive performance pyramid (NCPP)

The NCPP visualized the model ability by quantifying the resolution and fidelity of network inference results between Alpha-LFM and VCD-LFM during model optimization. To clarify the fitting ability of these two models on label data, such resolution and fidelity metrics were derived from the cut-off frequency difference (ΔKc) and the structural similarity discrepancy (ΔSSIM=1-SSIM) between the 3D reconstruction of the network and corresponding GT, where Kc is computed by decorrelation analysis^51^. Both ΔKc and ΔSSIM range between 0 and 1, where values close to 0 indicate a favorable combination of resolution and fidelity, while values close to 1 signify a low-quality image reconstruction. In our study, 196 LF patches (Size: 360×360) of mitochondrial outer membrane data were reconstructed by VCD and Alpha-Net during whole model training process. Specifically, we chose five different epochs (10, 100, 200, 210, and 300) to track this comprehensive performance of two models: For Alpha-Net, 10∼100 epochs denoted the decomposed optimization on each sub-network while 200∼300 epochs represented the progressive optimization process with multi-stage data (Noisy LFs, Clean LFs, De-aliased LFs and 3D SR); For VCD, 10∼300 denoted the optimization process under SR stacks and corresponding Noisy LFs. By calculating ΔKc and Δ SSIM of network inferences under the epochs, the performance scattering plots in network optimization process were produced (Fig. 2a). Besides, the convex hull computation of these scattering plots was used to generate the boundary line of statistics distribution and evaluate the performance deviation.

### Assessment of the resolution, fidelity and SBP of Alpha-LFM

In Fig. 2, we used the structural similarity (SSIM) and peak signal to noise ratio (PSNR) function in Matlab to assess the fidelity of our Alpha-Net reconstructions and other LFM techniques including LFD, VS-LFD, VCD utilizing the Airyscan data as reference in both the xy and yz planes. In Supplementary Fig. 9, the resolution-scaled Pearson coefficient (RSP) was quantified by SQUIRREL analysis^52^ using WF as reference. We applied decorrelation analysis to quantify the resolution of Alpha-Net results and other 3D microscopy implementations. The analysis was conducted using MATLAB. The axial resolution was measured by the sectorial resolution mode of the decorrelation analysis^51^, in which only the resolution in a narrow sectorial region along z direction was calculated. The space-bandwidth product (SBP) was calculated by SBP=FOV/(0.5δ)^3^ with δ as the system’s resolution, the factor 0.5 stemming from the Nyquist-Shannon sampling theorem and factor 3 representing three dimensions.

In Fig. 2l and Supplementary Fig. 14, the fidelity of Alpha-LFM was validated under varying light dose by adjusting the optical power and exposure time. Comparison between Alpha-LFM and 3D-SIM were conducted using the same objective (Olympus UPL, APO100XOHR) and identical light doses from 1.6 to 0.02 J/cm^2^ for obtaining the same volume with a depth of 4 μm.

### Cell culture and fluorescence labeling

U2OS cells were grown in culture medium containing McCoy’s 5A medium (Thermo Fisher Scientific) supplemented with 1% antibiotic-antimycotic (Thermo Fisher Scientific) and 10% fetal bovine serum (Thermo Fisher Scientific) at 37°C with 5% CO_2_ in a humidified incubator.

For labeling lysosomes in fixed U2OS cells, cells were first transfected with EGFP-Rab7A using Lipofectamine 2000 according to the standard protocol and cultured at 37 °C with 5% CO_2_ for an additional 24 h. Before imaging, the cells were fixed with 2% glutaraldehyde for 20 min.

For labeling tubulin in fixed U2OS cells, cells were seeded onto coverslips at 37°C with 5% CO_2_ for 12 h. Before fixation, the cells were washed with phosphate buffered saline (PBS, Thermo Fisher Scientific) at 37°C and then treated with fixing buffer (containing 3% paraformaldehyde (Electron Microscopy Sciences), 0.1% glutaraldehyde (Electron Microscopy Sciences), 0.2% Triton X-100 (Sigma-Aldrich)) for 15 min, then incubated with 0.2% Triton X-100 for 15 min and blocked with blocking buffer (3% bovine serum albumin (Sigma-Aldrich) and 0.05% Triton X-100 (Sigma-Aldrich)) for 20 min at room temperature. After that, cells were incubated with an anti-alpha tubulin antibody (Abcam, 1:500 dilution) overnight at 4 ° C. Subsequently, the primary antibody was removed and the cells were washed twice with PBS. Next, the cells were incubated with a secondary antibody (Abcam, labeled with Alexa Fluor 488, 1:400 dilution) for another 2 h at room temperature. The antibody was then removed and the cells were washed three times with PBS.

For labeling outer membranes of mitochondria and lysosomes in live U2OS cells, cells were first transfected with Tomm20-EGFP and Rab7A-mCherry2 using Lipofectamine 2000 according to the standard protocol and cultured at 37°C with 5% CO_2_ for an additional 24 h. Before imaging, remove old media and add fresh media.

For labeling mitochondrial matrix and chromosomes in live U2OS cells, cells were first transfected with Cox4-EGFP and H2B-mCherry2 using Lipofectamine 2000 according to the standard protocol and cultured at 37°C with 5% CO_2_ for an additional 8 h. After 6-8 hours of transfection, the cells were digested with 0.25% trypsin, seeded on cell culture dishes (20 mm diameter), and incubated for 36 hours at 37°C in 5% CO_2_.

For labeling peroxisomes in live U2OS cells, cells were first transfected with SKL-mApple using Lipofectamine 2000 according to the standard protocol and cultured at 37°C with 5% CO_2_ for an additional 24 h. Before imaging, remove old media and add fresh media. For ground truth images acquisition, cells were fixed with 4% paraformaldehyde (PFA) before imaging.

For labeling ER in live COS-7 cells, cells were first transfected with Sec61β-EGFP using Lipofectamine 2000 according to the standard protocol and cultured at 37°C with 5% CO_2_ for an additional 24 h. Before imaging, remove old media and add fresh media. For ground truth images acquisition, cells were fixed with 2% glutaraldehyde before imaging.

During the imaging process, cells were cultivated in phenol red free McCoy’s 5A medium (customized, Boster Biological Technology) within the confocal dishes. To ensure a stable environment, the cell in confocal dishes were cultured in the live cell microscope incubation system (TOKAIHIT) to maintain a consistent temperature of 37°C and a 5% CO_2_ atmosphere.

### Live-cell imaging

Light-field imaging of the lysosome outer membrane in live U2OS cells was implemented using our add-on light-field device with a × 60/1.3NA objective (Olympus UPlanSApo60XS2) at a volumetric imaging rate up to 333 Hz. This high-speed light-field imaging of live cells continued for 2 minutes with less than 50 % photobleaching, yielding 40000 SR reconstructions using Alpha-LFM. As a result, the fine deformation of lysosome outer membranes occurred in milliseconds could be observed by our Alpha-LFM in three dimensions. To compare the photon bleaching rate with other 3D microscopy implementations, we also imaged the lysosome outer membrane by Airyscan and LSFM microscope. The laser intensity and exposure time was carefully adjusted to ensure that the image SNRs by these microscopes were similar (SNR = 5.98, 5.87, 5.74, 5.99 for LSFM, Airyscan, 3D SIM and Alpha-LFM). To maintain sufficient SNR, LSFM, Airyscan and 3D SIM required 4 s, 4 minutes and 30 s for imaging a whole cell, respectively. The bleaching rates were then calculated using Matlab. Multiple areas were cropped and a threshold was used to discriminate the signal areas and background areas automatically. The photon bleaching rate over time was finally calculated using the following equation: *I*(*t*) = *I*_signal_(*t*) – *I*_background_(*t*))/max(*I*_signal_(*t*) – *I*_background_(*t*)), where *I*_signal_ and *I*_background_ are the mean intensity value of the multiple regions of signal and background images, respectively.

### Quantitative analysis of the interaction between mitochondria and lysosome

Interaction of mitochondrial and lysosomes was identified when the distance between a certain lysosome and the constriction site of a certain mitochondrion was measured smaller than 240 nm (close proximity) in at least three consecutive frames. We four-dimensionally captured such interaction between mitochondria and lysosomes in 17 live cells, where 41 ROIs that met the Lyso-Mito contact criteria in 2D MIP mode were identified, with 24 ROIs containing lysosome-mediated mitochondrial fission and other 17 ROIs containing mitochondrial fusion. Then, the real distance between lysosomes and mitochondria in these ROI was measured in true 3D mode using the commercial Imaris software. As a result, 10 false contacts were found in 2D results. We further calculated the fission and fusion velocity using self-written MATLAB code that included the following steps:

(i) Obtaining the coordination of the mitochondrial endpoints. For each frame of the time-series data: first, use the ‘imbinarize()’ function in Matlab to binarize the image. Then, employ the ‘bwskel()’ function to skeletonize the binary image. Finally, use the ‘endpoints’ method in the ‘bwmorph3()’ function to mark the endpoints of the skeletonized image and record the endpoint coordinates of all mitochondria.

(ii) Tracking the endpoints and identify the trajectories related to mitochondrial fission and fusion events. The code uses Matlab‘s ‘assignDetectionsToTracks’ function to track the sequence of the recorded endpoint coordinates along the time axis for the cropped ROIs, providing the motion trajectory of all endpoint along the time axis. Then, two longest motion trajectories were filtered as the trajectories related to the fission or fusion events while the discrete trajectories were removed. As long as the discontinuities occurred between frames in tracking, the ‘interp1()’ function is applied to complete the trajectories.

(iii) Velocity Acquisition. For each motion trajectory, the code calculates the distance between endpoints in all adjacent frames. Since frame interpolation has been performed, the instantaneous velocity were calculated through dividing the distance by the time interval. The calculated sequence of instantaneous velocity along the time corresponding to each motion trajectory were then averaged to obtain the overall fission or fusion velocity of this ROI.

### Experimental setup for long-term live-cell imaging

To observe the mitochondrial dynamics and using chromosomes to identify the cell stages, we used a 488/561 nm dual-band optical filter block (Chroma, 59904) in the microscope. Given that the cell division is highly sensitive to the phototoxicity, the power of LED (CoolLED, PE-800) was adjusted to 1% in 470 nm (30 μw) and 1% in 550 nm (30 μw). Since the tracking of high-speed fission and fusion event requires at least 10-seconds imaging rates and the entire cell cycle needs at least 48 hours observation, we designed a non-uniform imaging strategy for this long-term experiment. During this 30-min mid-term observation window, the imaging pipeline included the following steps:

(i) Implementing a Alpha-Net-based focusing strategy to correct sample drift along z axis. Then the 3D distribution of the mitochondria was quickly reconstructed by Alpha-Net and compared with the original distribution to calculate the z-drift distance, which would be further corrected by moving the objective using piezo scanner (COREMORROW, P73.Z). This process typically takes merely 1 second. (ii) Imaging the mitochondria dynamics at fixed depth of interest for 30 minutes, with an exposure time of 500 ms and interval time of 10 seconds. Meanwhile, identifying the related cell stage through the imaging of chromosomes in cell nuclei with an exposure time of 500 ms and interval time of 100 seconds.

### Tracking of the fates and morphology changes of mitochondria

We applied Mitometer^53^ to track the mitochondria dynamics reconstructed by Alpha-Net. Then the confident tracks and the corresponding morphology parameters can be exported as a mat file. To obtain the morphology changes of mitochondria throughout entire cycles, we first extracted the morphology parameter including major axis length, minor axis length, Volume, Solidity from the mat file in each timepoint. Then we mapped these parameters along the time and observed the changes throughout entire cycles. To further track the fates of each mitochondrion, we used the self-written Matlab code to extract the fission and fusion events in the mat file of the confident tracks. Then the mitochondria that exhibited the first fission were extracted with the fission sites being identified. We compared the volumes of two divided mitochondria in the next frame after fission and the smaller one will be defined as the small daughter. If the volume of the smaller one is less than 25% of the total length of two divided mitochondria, the fission will be defined as a peripheral fission, otherwise it is a midzone fission. Then the number of fission and fusion events occurred in these daughters were extracted and the daughters that undergo no fission or fusion were defined as no event. As a result, the event rates of these peripheral or midzone daughters were calculated and mapped.

## Data availability

The datasets generated and analyzed in this study are available from the corresponding authors upon request.

## Code availability

Customized Alpha-Net program and codes for quantitative analyses implemented in current study are either available at https://github.com/feilab-hust/Alpha-LFM or from the corresponding authors upon request.

## Acknowledgements

We are grateful to X. Duan for providing us the fluorescent cell samples and S. Mao for discussing the biological applications with us. This work was supported by the funding from National Natural Science Foundation of China (T2225014, 62375095, 21927802), National Key Research and Development Program of China (2022YFC3401100, 2023ZD0519900).

## Author contributions

P.F., L.Z. and C.Y. conceived the idea. P.F. and D.L. oversaw the project. L.Z., and J.S. developed the optical setups and acquired the experimental images. L.Z., J.S. and C.Y. developed the programs. L.Z. and J.S. processed the images. M.Z. prepared all the biological samples. L.C., Y.Z., C.Z., Y.Z., L.Z., C.Y., J.S., M.H., Y. H., S. W., H. C. and D.L. analyzed the data. L.Z., D.L, and P.F discussed and wrote the paper.

## References

1. Valm, A.M. et al. Applying systems-level spectral imaging and analysis to reveal the organelle interactome. Nature 546, 162–167 (2017).

2. Sahl, S.J., Hell, S.W. & Jakobs, S. Fluorescence nanoscopy in cell biology. Nat Rev Mol Cell Biol 18, 685–701 (2017).

3. Choquet, D., Sainlos, M. & Sibarita, J.B. Advanced imaging and labelling methods to decipher brain cell organization and function. Nat Rev Neurosci 22, 237–255 (2021).

4. Schermelleh, L., et al. Super-resolution microscopy demystified. Nat Cell Biol 21, 72–84 (2019).

5. Schermelleh, L., Carlton, P. M., Haase, S., Shao, L., Winoto, L., Kner, P.,…&Sedat, J. W. Subdiffraction multicolor imaging of the nuclear periphery with 3D structured illumination microscopy. Science 320.5881, 1332–1336 (2008).

6. Huang, B., Wang, W., Bates, M. & Zhuang, X. Three-dimensional super-resolution imaging by stochastic optical reconstruction microscopy. Science 319, 810–813 (2008).

7. Shao, L., Kner, P., Rego, E.H. & Gustafsson, M.G. Super-resolution 3D microscopy of live whole cells using structured illumination. Nat Methods 8, 1044–1046 (2011).

8. York, A.G. et al. Instant super-resolution imaging in live cells and embryos via analog image processing. Nat Methods 10, 1122–1126 (2013).

9. Chen, B.C. et al. Lattice light-sheet microscopy: imaging molecules to embryos at high spatiotemporal resolution. Science 346, 1257998 (2014).

10. Li, D., et al. ADVANCED IMAGING. Extended-resolution structured illumination imaging of endocytic and cytoskeletal dynamics. Science 349, aab3500 (2015).

11. Wu, Y. et al. Multiview confocal super-resolution microscopy. Nature 600, 279–284 (2021).

12. Li, X. et al. Three-dimensional structured illumination microscopy with enhanced axial resolution. Nature Biotechnology 41, 1307–1319 (2023).

13. Huff, J. The Airyscan detector from ZEISS: confocal imaging with improved signal-to-noise ratio and super-resolution. Nature Methods 12, i-ii (2015).

14. Gao, L. et al. Noninvasive imaging beyond the diffraction limit of 3D dynamics in thickly fluorescent specimens. Cell 151, 1370–1385 (2012).

15. Broxton, M. et al. Wave optics theory and 3-D deconvolution for the light field microscope. Optics express 21, 25418–25439 (2013).

16. Cohen, N. et al. Enhancing the performance of the light field microscope using wavefront coding. 22, 24817–24839 (2014).

17. Prevedel, R. et al. Simultaneous whole-animal 3D imaging of neuronal activity using light-field microscopy. Nature methods 11, 727–730 (2014).

18. Guo, C., Liu, W., Hua, X., Li, H. & Jia, S. Fourier light-field microscopy. Opt Express 27, 25573–25594 (2019).

19. Wu, J. et al. Iterative tomography with digital adaptive optics permits hour-long intravital observation of 3D subcellular dynamics at millisecond scale. Cell 184, 3318–3332 e3317 (2021).

20. Zhang, Z. et al. Imaging volumetric dynamics at high speed in mouse and zebrafish brain with confocal light field microscopy. Nat Biotechnol 39, 74–83 (2021).

21. Hua, X., Liu, W. & Jia, S. High-resolution Fourier light-field microscopy for volumetric multi-color live-cell imaging. Optica 8, 614–620 (2021).

22. Lu, Z. et al. Virtual-scanning light-field microscopy for robust snapshot high-resolution volumetric imaging. Nat Methods (2023).

23. Han, K., et al. 3D super-resolution live-cell imaging with radial symmetry and Fourier light-field microscopy. Biomedical Optics Express 13 (2022).

24. Weigert, M. et al. Content-aware image restoration: pushing the limits of fluorescence microscopy. Nature methods 15, 1090–1097 (2018).

25. Qiao, C. et al. Rationalized deep learning super-resolution microscopy for sustained live imaging of rapid subcellular processes. Nat Biotechnol (2022).

26. Wang, Z. et al. Real-time volumetric reconstruction of biological dynamics with light-field microscopy and deep learning. Nat Methods 18, 551–556 (2021).

27. Zhu, L., Yi, C. & Fei, P. A practical guide to deep-learning light-field microscopy for 3D imaging of biological dynamics. STAR Protoc 4, 102078 (2023).

28. Zhu, T. et al. High-speed large-scale 4D activities mapping of moving C. elegans by deep-learning-enabled light-field microscopy on a chip. Sensors and Actuators B: Chemical 348 (2021).

29. Guo, Y. et al. in Proceedings of the IEEE/CVF conference on computer vision and pattern recognition 5407–5416 (2020).

30. Lempitsky, V., Vedaldi, A. & Ulyanov, D. in 2018 IEEE/CVF Conference on Computer Vision and Pattern Recognition 9446–9454 (IEEE, 2018).

31. Chen, J. et al. Three-dimensional residual channel attention networks denoise and sharpen fluorescence microscopy image volumes. Nat Methods 18, 678–687 (2021).

32. Huff, J. et al. The new 2D Superresolution mode for ZEISS Airyscan. Nature Methods 14, 1223–1223 (2017).

33. Gustafsson, M.G. et al. Three-dimensional resolution doubling in wide-field fluorescence microscopy by structured illumination. Biophysical journal 94, 4957–4970 (2008).

34. Sapoznik, E. et al. A versatile oblique plane microscope for large-scale and high-resolution imaging of subcellular dynamics. Elife 9 (2020).

35. Voleti, V. et al. Real-time volumetric microscopy of in vivo dynamics and large-scale samples with SCAPE 2.0. Nat Methods 16, 1054–1062 (2019).

36. Guo, Y. et al. Visualizing Intracellular Organelle and Cytoskeletal Interactions at Nanoscale Resolution on Millisecond Timescales. Cell 175, 1430–1442 e1417 (2018).

37. Boutry, M. & Kim, P.K. ORP1L mediated PI(4)P signaling at ER-lysosome-mitochondrion three-way contact contributes to mitochondrial division. Nat Commun 12, 5354 (2021).

38. Wong, Y.C., Ysselstein, D. & Krainc, D. Mitochondria-lysosome contacts regulate mitochondrial fission via RAB7 GTP hydrolysis. Nature 554, 382–386 (2018).

39. Kleele, T. et al. Distinct fission signatures predict mitochondrial degradation or biogenesis. Nature 593, 435–439 (2021).

40. Mishra, P. & Chan, D.C. Mitochondrial dynamics and inheritance during cell division, development and disease. Nat Rev Mol Cell Biol 15, 634–646 (2014).

41. Chen, A. et al. in Proceedings of the IEEE/CVF International Conference on Computer Vision 14124–14133 (2021).

42. Mildenhall, B. et al. Nerf: Representing scenes as neural radiance fields for view synthesis. Communications of the ACM 65, 99–106 (2021).

## References

43. Zhao, F., et al. Deep-learning super-resolution light-sheet add-on microscopy (Deep-SLAM) for easy isotropic volumetric imaging of large biological specimens. Biomedical Optics Express 11 (2020).

44. Zhang, Y. et al. in Proceedings of the European conference on computer vision (ECCV) 286–301 (2018).

45. Mo, Y., Wang, Y., Xiao, C., Yang, J. & An, W. Dense dual-attention network for light field image super-resolution. IEEE Transactions on Circuits and Systems for Video Technology 32, 4431–4443 (2021).

46. He, K., Zhang, X., Ren, S. & Sun, J. in Proceedings of the IEEE conference on computer vision and pattern recognition 770–778 (2016).

47. Qiao, C. et al. Evaluation and development of deep neural networks for image super-resolution in optical microscopy. Nat Methods 18, 194–202 (2021).

48. Guo, C., Liu, W., Hua, X., Li, H. & Jia, S. Fourier light-field microscopy. Opt Express 27, 25573–25594 (2019).

49. Wang, Y., et al. in Computer Vision – ECCV 2020. (eds. A. Vedaldi, H. Bischof, T. Brox & J.- M. Frahm) 290-308 (Springer International Publishing, Cham; 2020).

50. Ibtehaz, N.&Rahman, M.S. MultiResUNet: Rethinking the U-Net architecture for multimodal biomedical image segmentation. Neural Networks 121, 74–87 (2020).

51. Descloux, A., Grußmayer, K.S. & Radenovic, A. Parameter-free image resolution estimation based on decorrelation analysis. Nature Methods 16, 918–924 (2019).

52. Culley, S. et al. Quantitative mapping and minimization of super-resolution optical imaging artifacts. Nat Methods 15, 263–266 (2018).

53. Lefebvre, A.E.Y.T., Ma, D., Kessenbrock, K., Lawson, D.A. & Digman, M.A. Automated segmentation and tracking of mitochondria in live-cell time-lapse images. Nature Methods 18, 1091–1102 (2021).

