## Supplementary information for "Adaptive-learning physics-aware light-field microscopy enables day-long and millisecond-scale super-resolution imaging of 3D subcellular dynamics"

### Supplementary Figures

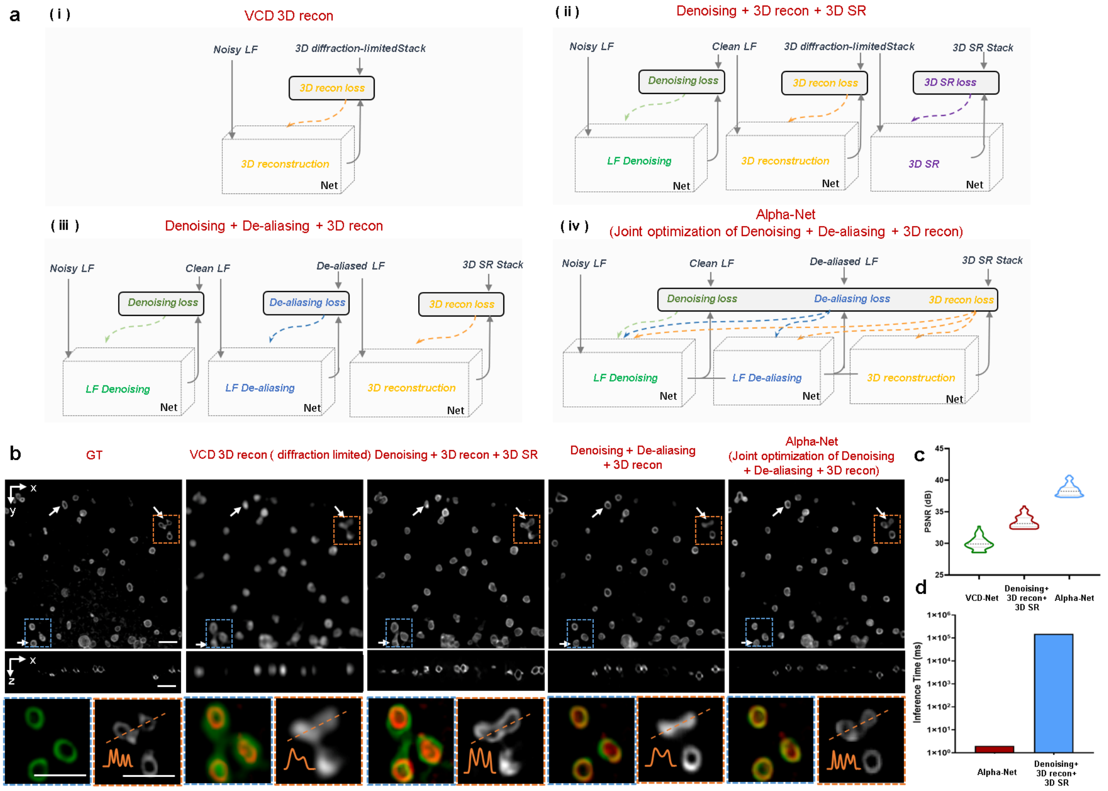

**Supplementary Figure 1. The verification of different model design strategy in model design. a,** The schematic illustrating the different decomposed and training strategy. (i) The VCD network are employed end-to-end training using training pairs of Noisy LF and 3D diffraction-limited stack. (ii) The conventional strategy that decompose the network into Denoising, 3D reconstruction and 3D SR sub-works are trained using Noisy LF, 3D diffraction-limited stack and 3D SR stack and optimized independently with the input training data for each subsequent module being based on the output of the preceding one. (iii) Our model design strategy decomposes the network into LF Denoising, LF De-aliasing and 3D reconstruction and employs Noisy LF, Clean LF, De-aliased LF and 3D SR stack for training. The sub-networks are optimized independently with the input training data for each subsequent module being based on the output of the preceding one. (iv) Our Alpha-Net decomposes the network into LF Denoising, LF De-aliasing and 3D reconstruction decomposed strategy and employs decomposed-progressive optimizing strategy for network optimizing. **b**, The xy and xz MIP of the lysosome results reconstructed by four decomposed and training strategy. The merged results of the ROI indicated by the blue box in top row with GT are shown in the bottom left corner in each panel. The magnified view of the ROI indicated by the orange box in upper row is shown in bottom right corner in each panel. The inset plot shown in bottom right of each panel shows the intensity profiles indicated by the orange dashed line. **c,** The peak signal-to-noise ratio (PSNR) metrics indicating the signal fidelity of VCD, Denoising + VCD+3D SR and Alpha-Net using GT as reference (n=30). **d,** The inference time of Alpha-Net and Denoising + VCD + 3D SR. Scale bar, 2μm

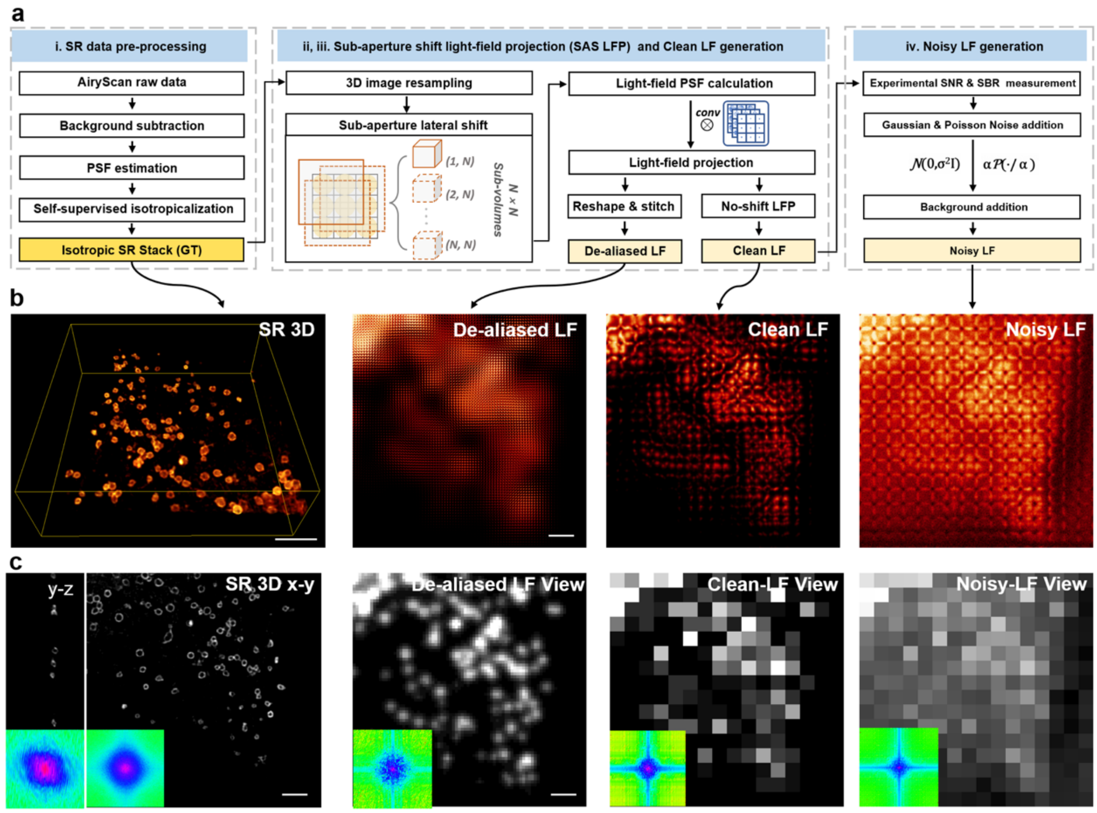

**Supplementary Figure 2. Physics-embedded hierarchical training data generation procedure.** **a,** Three main steps to generate the hierarchical De-aliased LF, Clean LF and Noisy LF from the same 3D SR image: (i) The Airyscan data is firstly enhanced through applying axial-to-lateral self-supervised isotropic learning to create 3D SR data with isotropic resolution. (ii) The 3D SR volume is shifted by a distance smaller than the size of single lenslet along the horizontal (x axis) and vertical (y axis) direction. The shift in SAS LFP operation covers 3 pixels during the 5×5 times scan . Such shifted 3D volumes are then projected based on wave optics model, yielding 25 LFs. The shifted light field projections are then stitched to generate the De-aliased LF images according to the arrangement of the light. (iii) The central LF without shift in step (ii) is used as the Clean LF data. (iv) The Clean LF in step (iii) is normalized to the same grayscale range of experimental LFs, and then added with diverse noise (Gaussian noise with different deviation and Poisson Noise) common in the experimental LFs to generate Noisy LF. **b,** The 3D SR image of lysosomes (by airy scan confocal microscope) and the semi-synthetic De-aliased LF, Clean LF and Noisy LF all derived from it. **c,** The corresponding projection views of the 3D SR, De-aliased LF, Clean LF and Noisy LF with the insets comparing their spectrum in Fourier domain. Scale bar, 5μm.

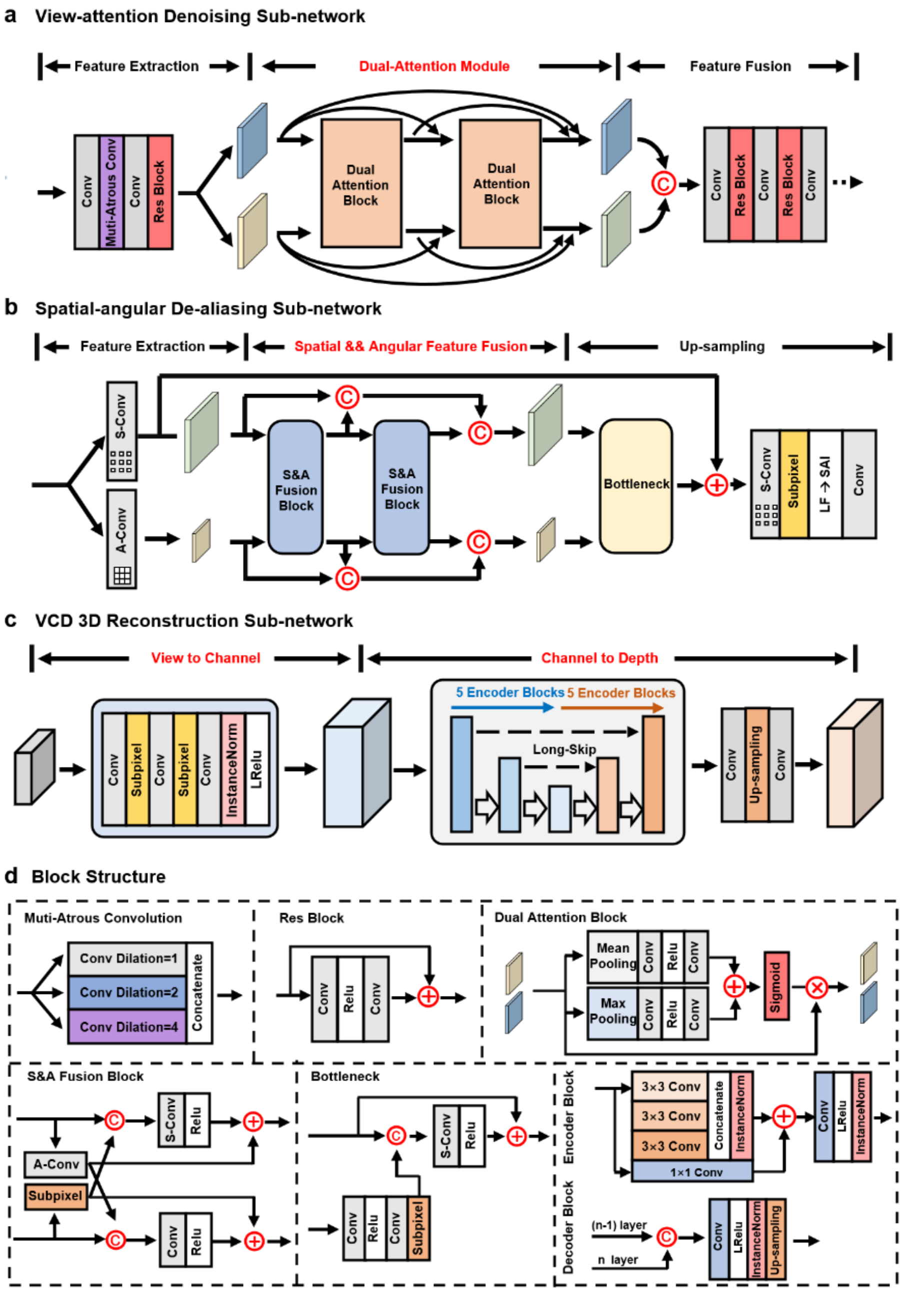

**Supplementary Figure 3. Network architecture of Alpha-Net model.** Our proposed Alpha-Net network is composed of three sub-networks: **a,** View-attention denoising network. We used 2 dual-attention blocks (contains "view-attention" branch and "channel attention" branch) to achieve deep feature extraction and adaptive view-wise processing. Before these attention blocks, dilate convolution and res-blocks consist of PFE (primary feature extraction) module. At the end of this module, plain convolution and res-blocks were used to fuse previous features and achieve denoised output. **b,** Spatial-angular de-aliasing network. Similar with the topology of denoising network, we adopted three modules: PFE, Spatial-angular feature fusion module[1] and Upsampling module. At the feature fusion part, both spatial convolution and angular convolution processed the extracted features. **c,** VCD 3D reconstruction network. This network follows the design of previous VCD structure[2]: PFE module and U-net module. The difference lies in the utilization of multi-res blocks to replace the vanilla convolution layers in VCD encoder blocks. **d,** The structure of blocks used in (a), (b) and (c).

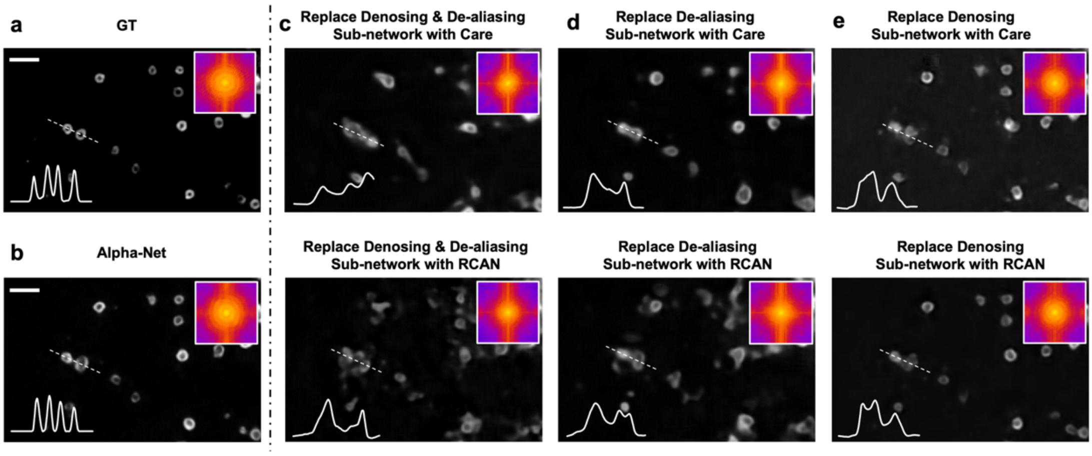

**Supplementary Figure 4. Ablation study evaluating the replacement of Alpha-Net sub-networks with state-of-the-art image enhancement networks.** An ablation study was conducted to compare the performance of the View-attention denoising and Spatial-angular de-aliasing sub-networks in Alpha-Net with alternative modules, CARE[3] and RCAN[4]. The MIPs of a lysosome-labelled cell captured with Airyscan are presented for comparison (GT, **a**), Alpha-Net reconstruction (**b**), and the results obtained from networks with substituted modules (**c-e**). The insets in the upper left of each image display the corresponding spectra in Fourier domain, which the insets in the bottom left show the normalized intensity profiles along the white dotted lines. The comparative analysis highlights the efficacy of Alpha-Net sub-networks in reconstructing high-fidelity images. Scale bar: 2 μm.

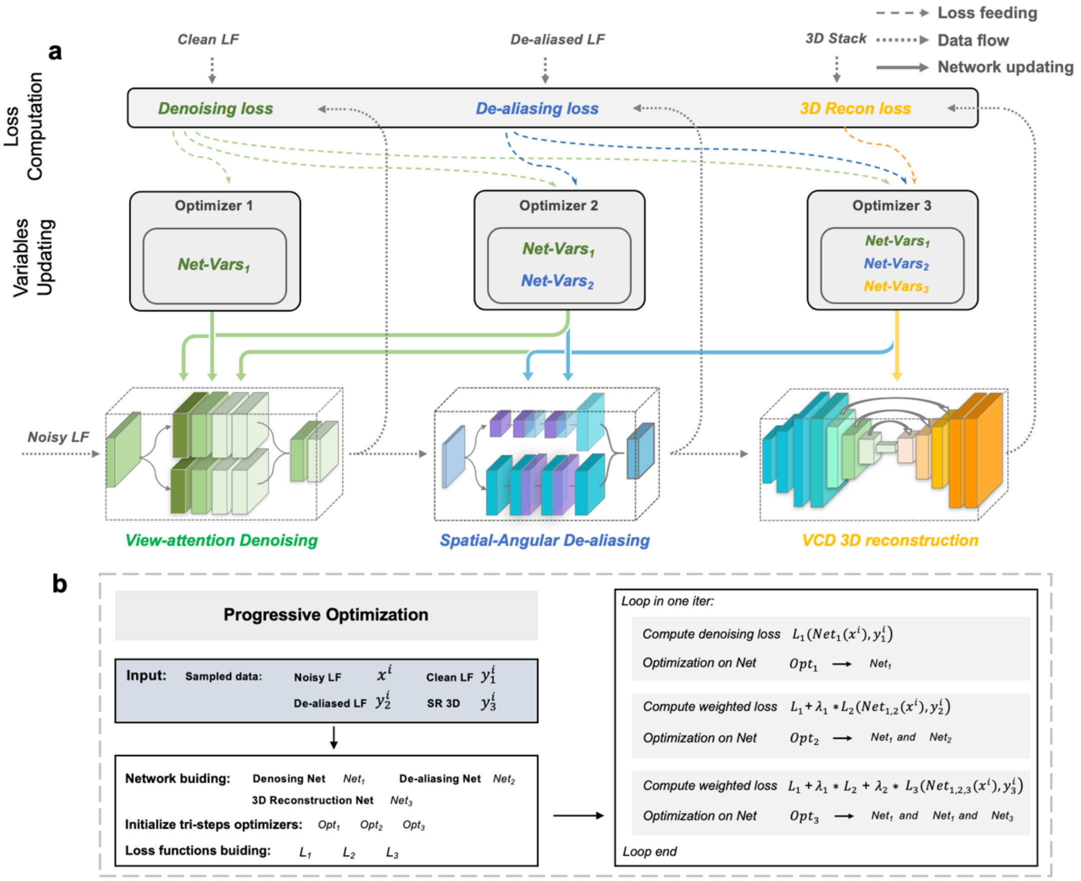

**Supplementary Figure 5. The principle of decomposed-progressive optimization strategy. a,** The process schematic illustrating the cooperation of three network optimizers in progressive Alpha-Net optimization. The whole model is optimized as the following orders: 1) Compute the denoising loss and update the parameters of LF denoising sub-network; 2) Compute the weighted denoising & de-aliasing loss and update the parameters of LF denoising and LF de-aliasing sub-networks; 3) Triple sub-networks are updated simultaneously with the guidance of weighted denoising & de-aliasing & 3D reconstruction loss. **b,** Pseudo codes describe the workflow of progressive optimization in one iteration. In the $i^{th}$ iteration, the paired data ($x^i$, $y_1^i$, $y_2^i$, $y_3^i$) sampled from training dataset were used to compute the denoising loss ($L_1$), de-aliasing loss ($L_2$) and 3D reconstruction loss ($L_3$). Such computed losses were then combined according to the pre-defined coefficients ($\lambda_1$, $\lambda_2$, $\lambda_3$), and passed to three optimizers ($Opt_1$, $Opt_2$, $Opt_3$) to update three sub-networks ($Net_1$, $Net_2$, $Net_3$) progressively.

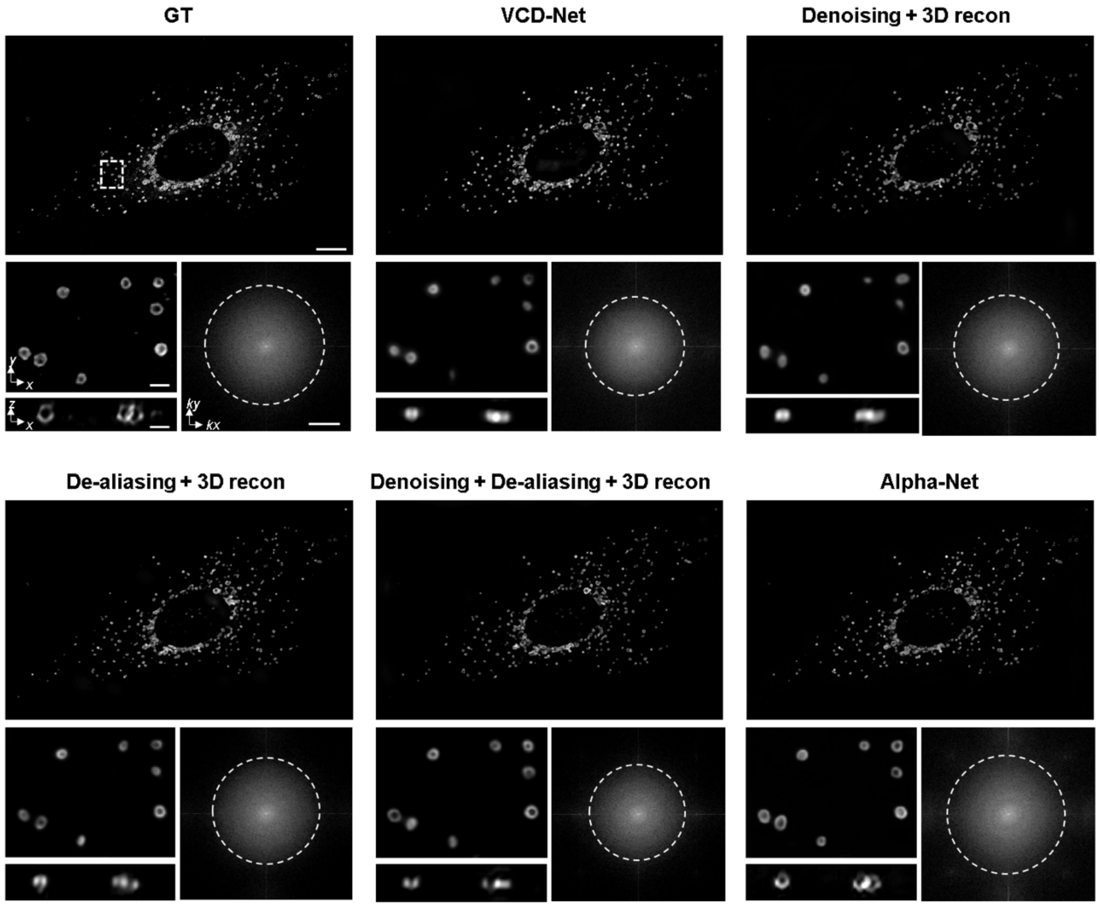

**Supplementary Figure 6. Evaluation of individual module in Alpha-Net.** An ablation study is conducted to assess the effect of each module in the Alpha-Net. The 3D SR image of a lysosome-labelled cell (GT), the corresponding light-field 3D reconstructions by standard VCD, Denoising + VCD recon (two-step network), De-aliasing + VCD recon (two-step network), Denoising + De-aliasing + VCD recon (three-step network without joint optimization) and full Alpha-Net are comparatively shown from **a** to **f**, respectively. The x-y and x-z MIPs of the same small regions (indicated by the white box in **a)** are magnified to clearly show the reconstruction quality from different conditions. Their corresponding Fourier spectrums are also shown to compare the resolution. In this evaluation, two or three modules are trained independently, with the input training data for each subsequent module being based on the output of the preceding one. Only Alpha-Net includes the decomposed-progressive optimizing strategy and trained by simultaneously optimizing the loss function of three module based on designed weight. Scale bar, 10 μm (top in each panel), 1 μm (bottom left), 10 $\mu m^{-1}$ (bottom right).

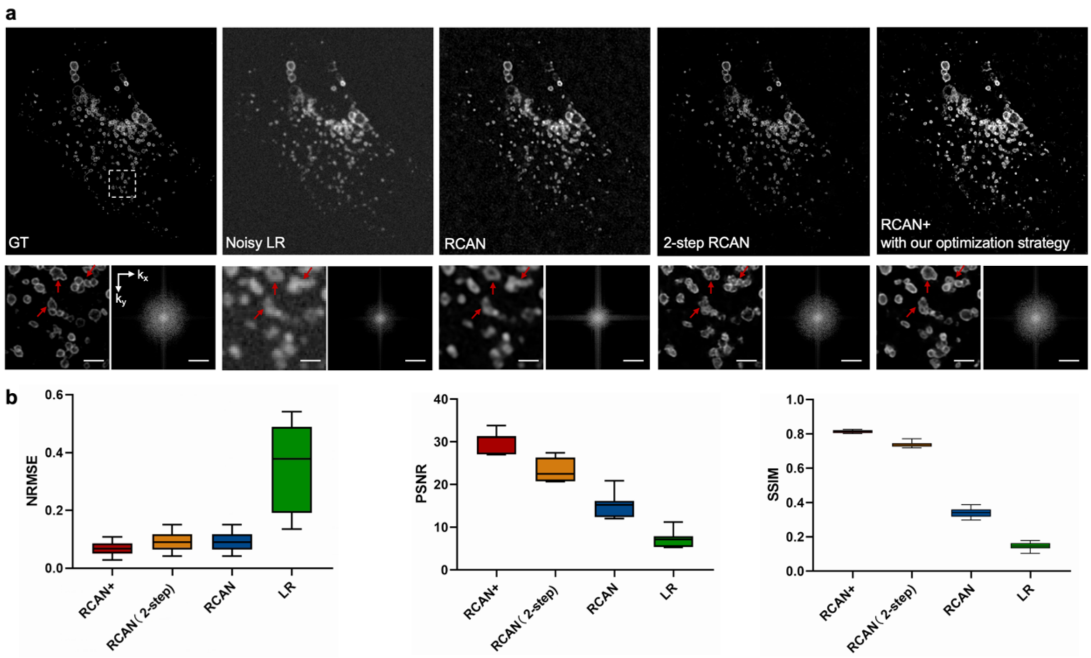

**Supplementary Figure 7. Our DPO strategy efficiently improves the reconstruction fidelity for 3D-RCAN. a,** The SR outputs of lysosomes reconstructed from noisy LR inputs using RCAN, 2-step RCAN (Denoising+SR) and RCAN+ (RCAN model optimized using our DPO strategy). The enlarged view indicated by white dotted box in upper row is shown in the bottom left corner of each panel. The red arrows indicate the noticeable errors in RCAN and 2-step RCAN results whereas being accurately resolved by RCAN+. The corresponding Fourier spectrum is shown in bottom right corner of each panel. Scale bar, 10 μm (top in each panel), 2 μm (bottom left), 1/280 nm$^{-1}$ (bottom right). **b,** The reconstruction fidelity of RCAN, 2-step RCAN and RCAN+ are demonstrated by calculating the normalized root mean square error (NRMSE), peak signal to noise ratio (PSNR) and structure similarity (SSIM), with using GT as reference. n$\geqslant$15 volumes are used for each analysis. n$\geqslant$8 volumes are used for each analysis. The center line represents the median, the box limits represent the lower and upper quartiles, and the whiskers represent the min and max value.

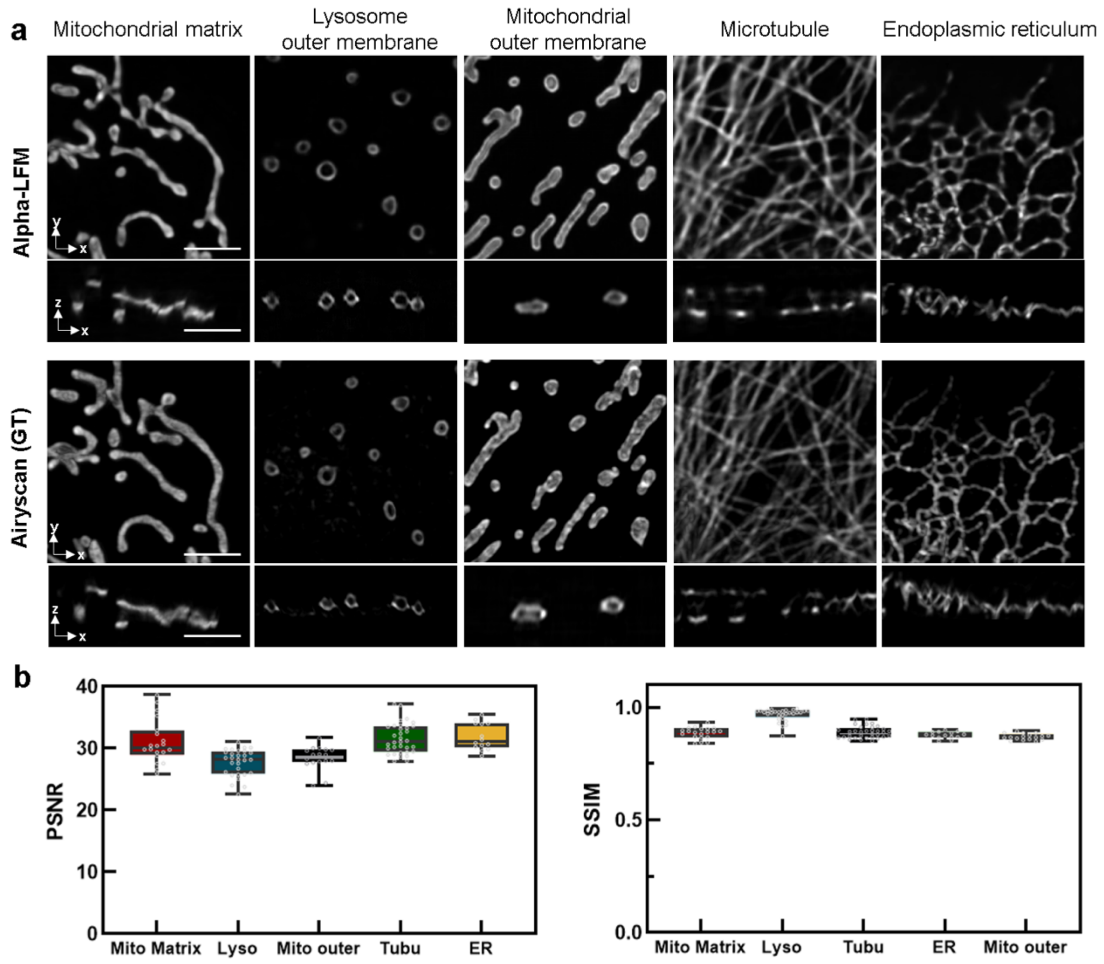

**Supplementary Figure 8. The fidelity of Alpha-Net on diverse subcellular structures. a,** The xy and xz MIPs of the Alpha-Net results and enhanced Airyscan data (GT) of diverse subcellular structures, including mitochondrial matrix (Mito Matrix), outer membrane of lysosome (Lyso), outer membrane of mitochondria (Mito outer), microtubule, endoplasmic reticulum (ER). Scale bar, 3μm.**b,** The reconstruction fidelity of Alpha-Net are demonstrated by calculating the peak signal to noise ratio (PSNR) and structure similarity (SSIM), with using Airyscan data as reference. n⩾15 volumes are used for each analysis. The center line represents the median, the box limits represent the lower and upper quartiles, and the whiskers represent the min and max value. The spots on the box indicates all calculated values.

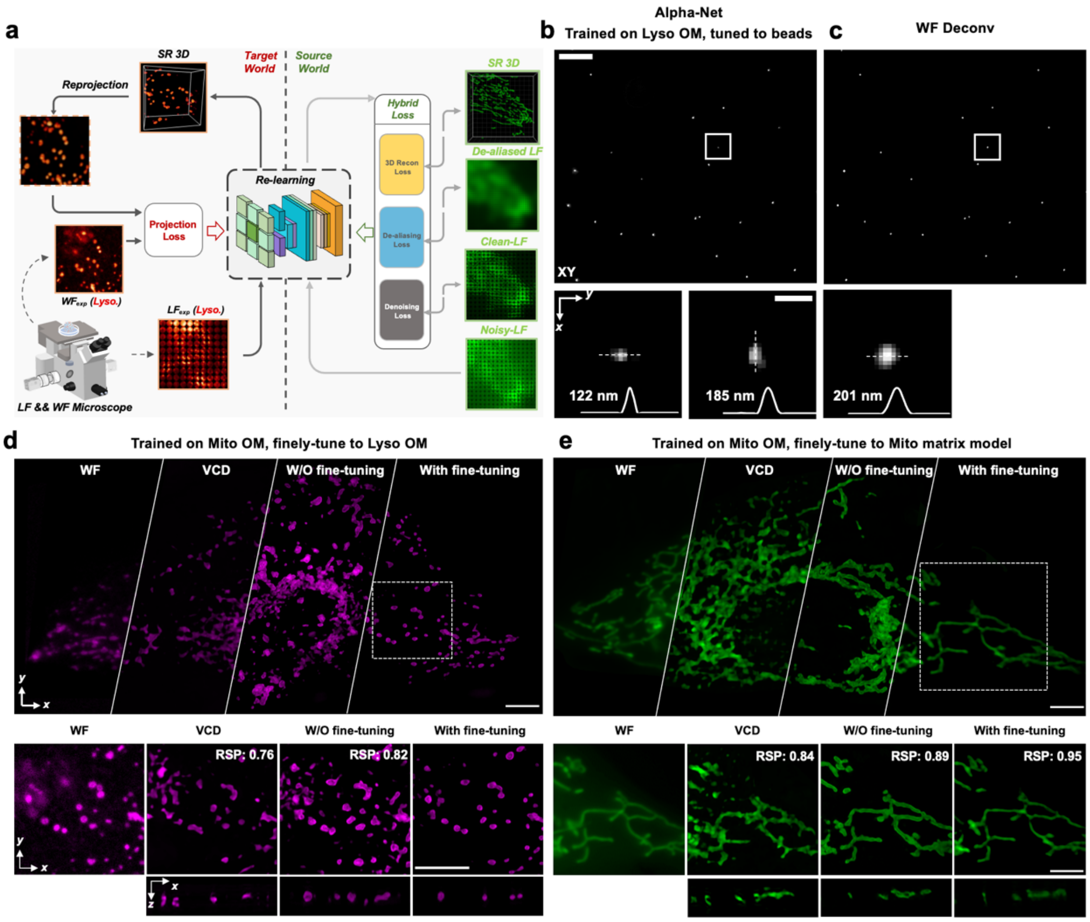

**Supplementary Figure 9. The fine-tuning strategy in Alpha-Net for improved generalization on diverse samples. a,** The schematic illustrating the adaptive-tuning strategy. **b,** The inference results of Alpha-Net on LF image of fluorescent beads. Alpha-Net were initially trained on outer membranes of lysosome (Lyso OM) and adaptively-tuned using 2D wide-field deconvolution results of fluorescent beads. **c,** 2D wide-field imaging results of in-situ fluorescent beads with deconvolution. **d, e** The Alpha-Net network originally trained on outer membrane of mitochondria (Mito OM) has been adaptively tuned to reconstruct the accurate 3D structure of unseen data: either Lyso OM (**d**) or matrix (**e**) with the guidance of few corresponding WF measurements. The comparison between WF images and reconstructions of VCD-Net and Alpha-Net with and without (W/O) finely-tuned model has verified the strong effect of fine-tuning strategy. The resolution-scaled Pearson coefficient (RSP) values quantifying the SR accuracy of the reconstructions were calculated by SQUIRREL analysis using in-focus WF images as reference. Scale bar, 10 µm.

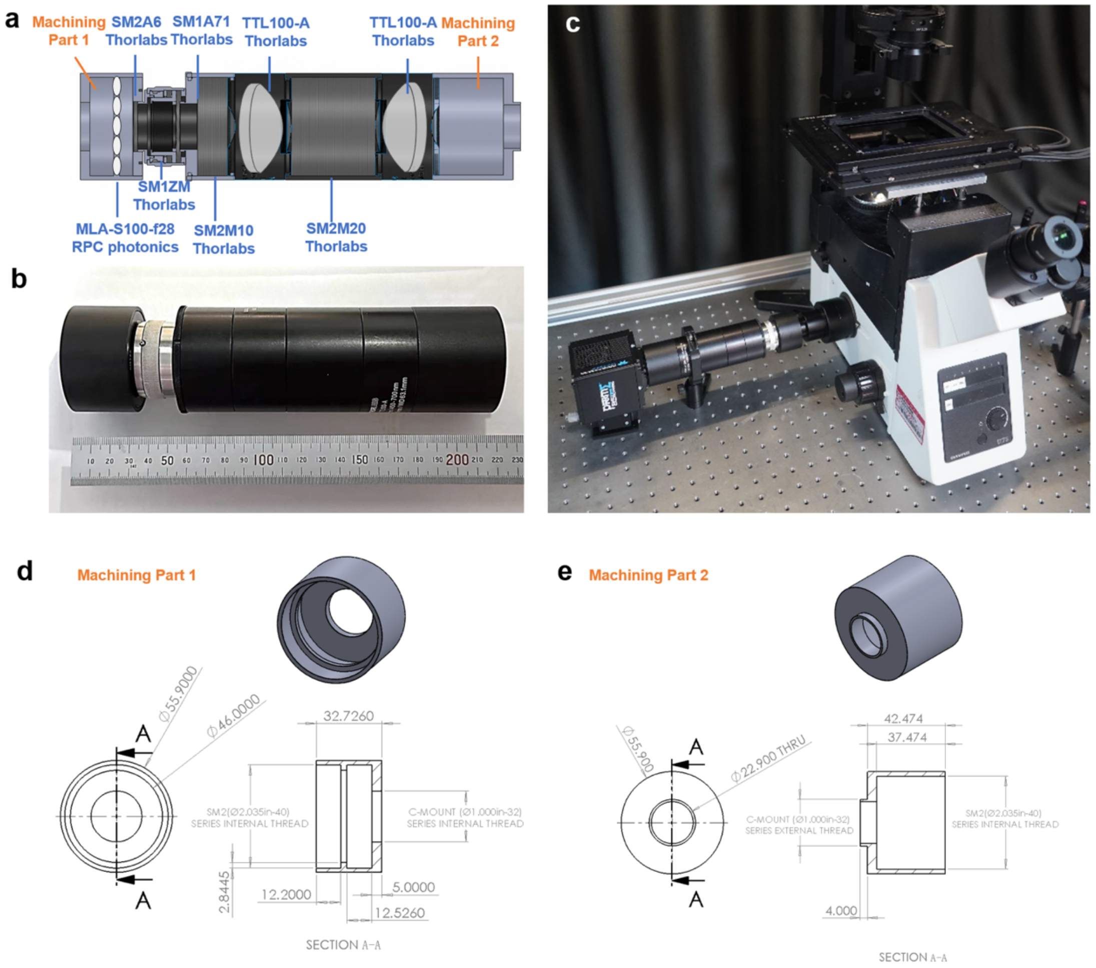

**Supplementary Figure 10. Light-field microscopy based on the simple retrofit of an inverted fluorescence microscope using our compact light-field add-on device. a,** The layout of light-field add-on device majorly containing off-the-shelf optical and mechanical elements (blue) and two customized machining parts (orange). **b,** The photograph of the whole device measured ~220 mm in length and ~50 mm in diameter. **c,** The picture of a commercial inverted microscope (Olympus, IX73) mounted with our light-field add-on, which together with the Alpha-Net program, can immediately upgrade ordinary 2D diffraction-limited wide-field imaging into advanced 3D super-resolution light-field imaging. **d, e** The designs of two customized mechanical parts.

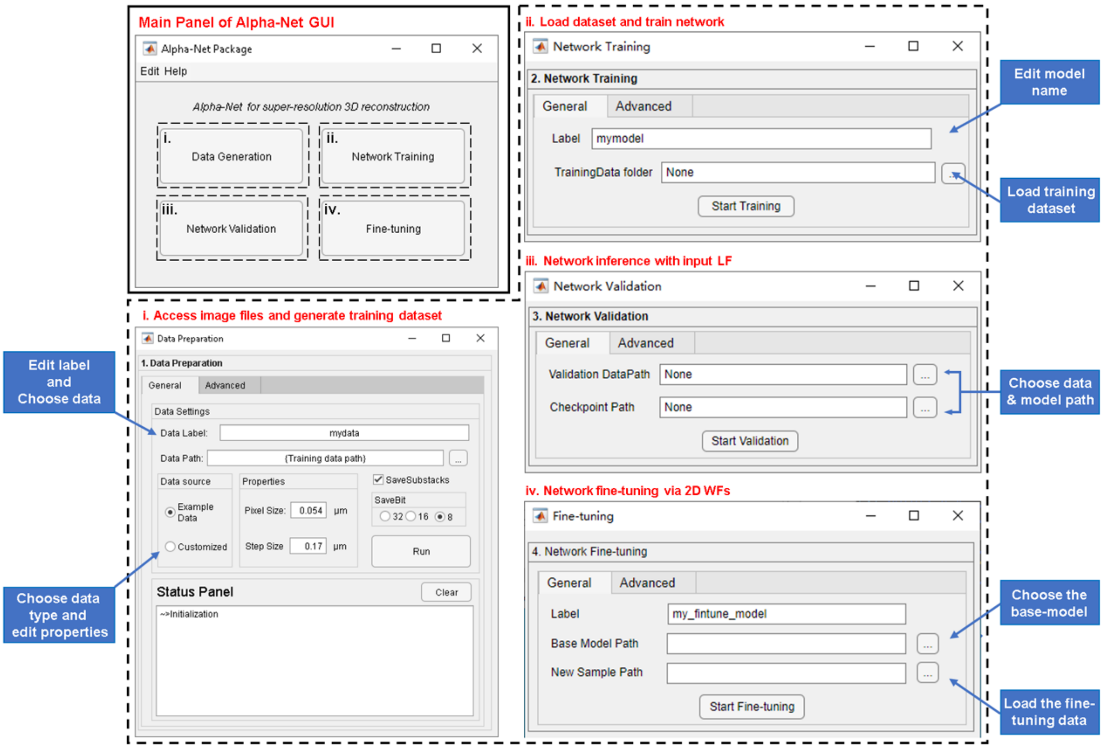

**Supplementary Figure 11. Open-source Alpha-Net program with graphic user interface (GUI) provided.** The Alpha-Net GUI contains 4 main parts: Data preparation, Network training, Network validation and Fine tuning. Users can customize their settings in the “General settings” panel and “Advanced Settings” panel. For fast implementation of Alpha-Net, users only need to set the “General settings”: (i) Input the training data’s name and choose the path of high-resolution stacks. Then, choose the data’s source and edit the properties of input data. Finally, click “Run” to start dataset preparation; (ii) Edit the name of network model and load training dataset. (iii) After the network optimization lasting several hours, a trained Alpha-Net model would be automatically saved. Then, users need to choose the path of saved models and LFs to start fast network inference; (iv) To enhance the model ability on different samples, a trained Alpha-Net model can be finely tuned using paired LF and 2D WF measurements, to make itself adaptive to the unseen new samples. Users can choose the path of base-model and paired LF-WF dataset for further network fine-tuning. A detailed tutorial has been provided to facilitate the new users to get started.

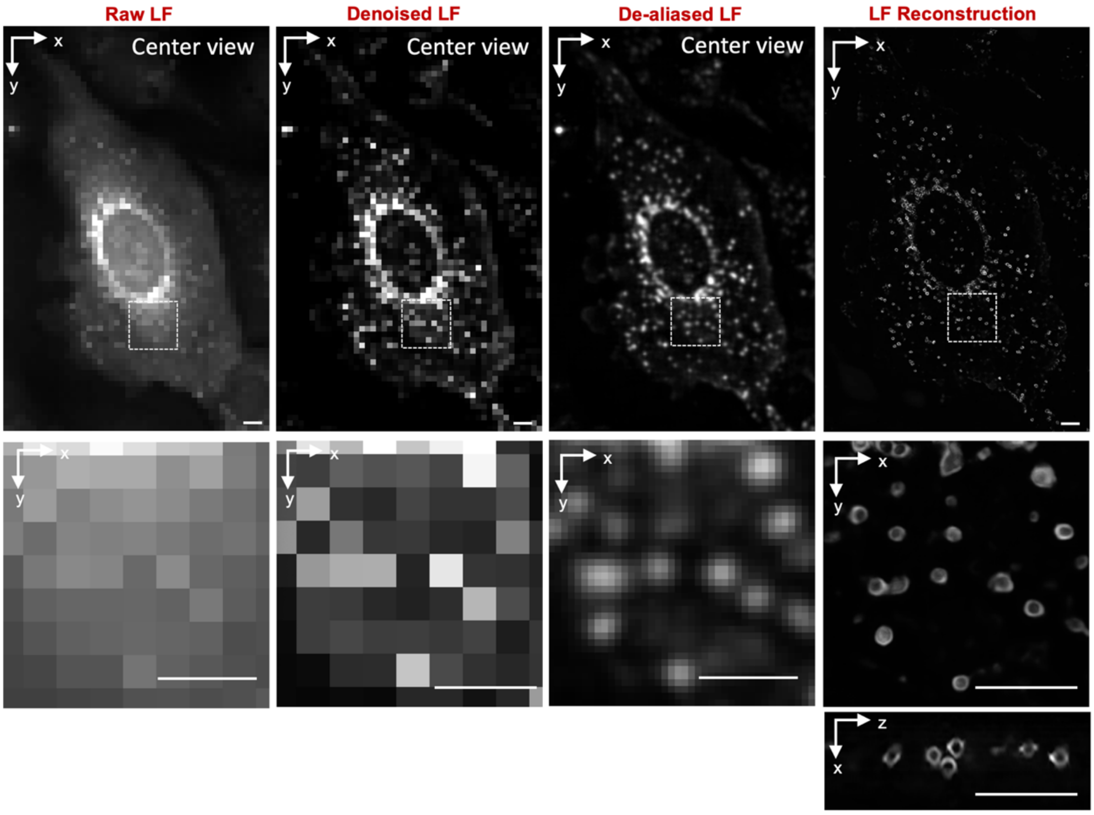

**Supplementary Figure 12. The enhancement achieved by the sub-networks of Alpha-Net.** The center views of raw LF and corresponding denoised, de-aliased and reconstruction results of Alpha-Net are shown in the top row in each panel. The enlarged view indicated by white dotted box in upper row is shown in the bottom left corner of each panel. The corresponding Fourier spectrum is shown in bottom right. Scale bar, 10 µm (top in each panel), 2 µm (bottom left), 1/280 $nm^{-1}$ (bottom right).

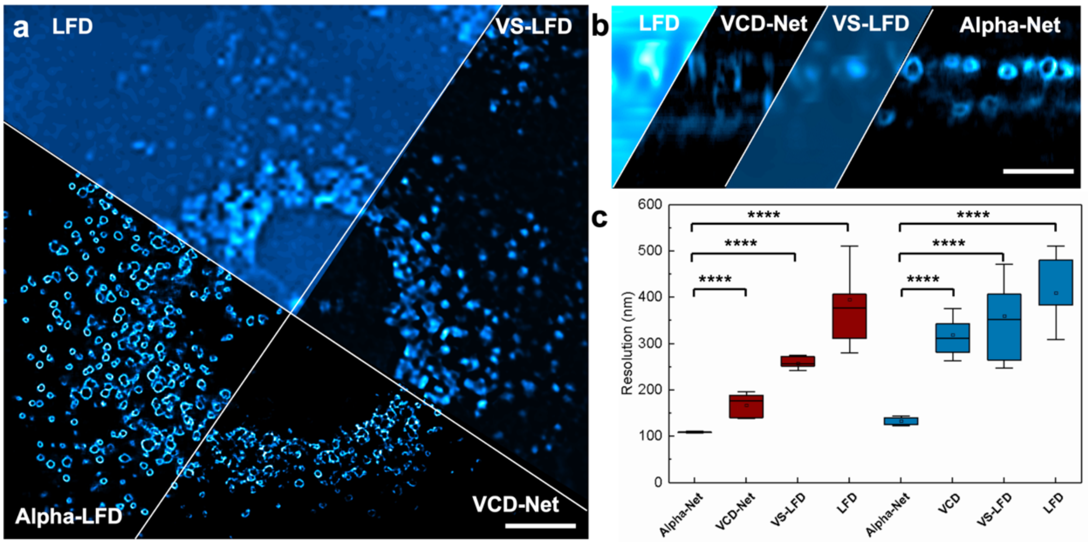

**Supplementary Figure 13. Reconstruction comparison of Alpha-LFM with LFD, VS-LFM and VCD-LFM.** The LF image acquired using microlens array with a pitch size of 45.5 µm and a focal length of 1.6 mm under a ×60/ NA 1.3 objective (Olympus UPlanSApo60XS2). **a, b** The orthogonal MIPs of the results reconstructed by LFD, VS-Net, VCD-Net and Alpha-Net. **c,** Decorrelation analysis quantifying the lateral and axial resolution of LFD, VS-Net, VCD-Net and Alpha-Net (n=9 volumes). The center line indicates the median, the box limits denote the lower and upper quartiles, and the whiskers represent the minimum and maximum values. Scale bar, 10 µm.

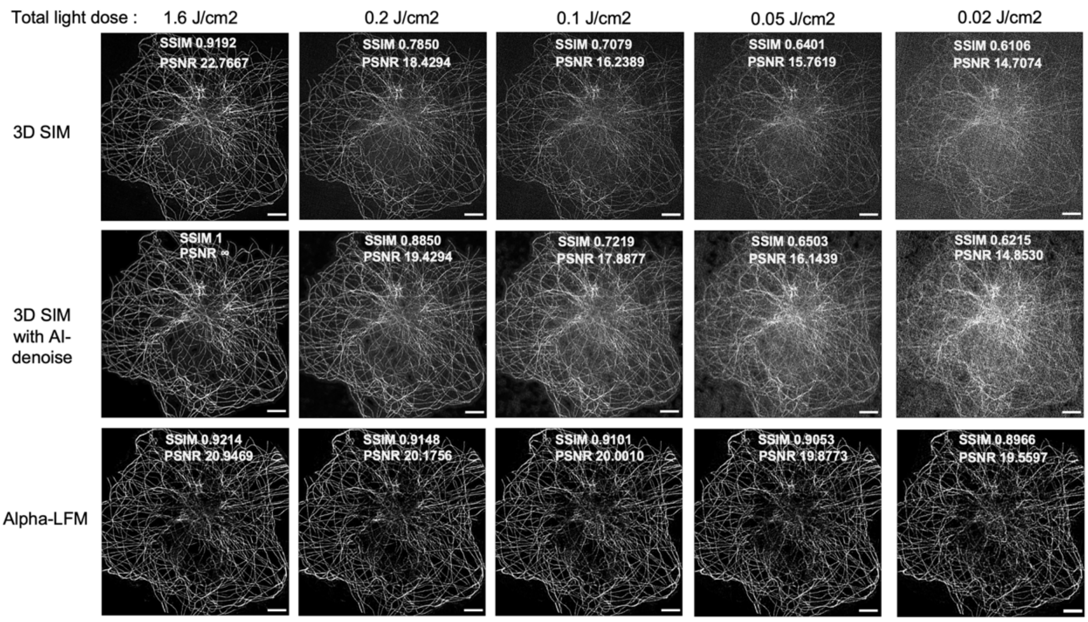

**Supplementary Figure 14. The performance comparisons between Alpha-LFM and 3D SIM under varying light dose used to acquire a volume.** MIPs of microtubules (tagged with Sec61β-EGFP) in a fixed COS-7 cell acquired by 3D-SIM, 3D-SIM with AI-denoising[4] and Alpha-LFM under varying total light dose (1.6 - 0.02 J/cm$^2$) used to capture a volume. The fidelity of 3D-SIM, AI-denoised 3D-SIM and Alpha-LFM are demonstrated by calculating SSIM and PSNR with using AI-denoised 3D-SIM data under the total light dose of 1.6 J/cm$^2$ as reference. Scale bar, 5 μm .

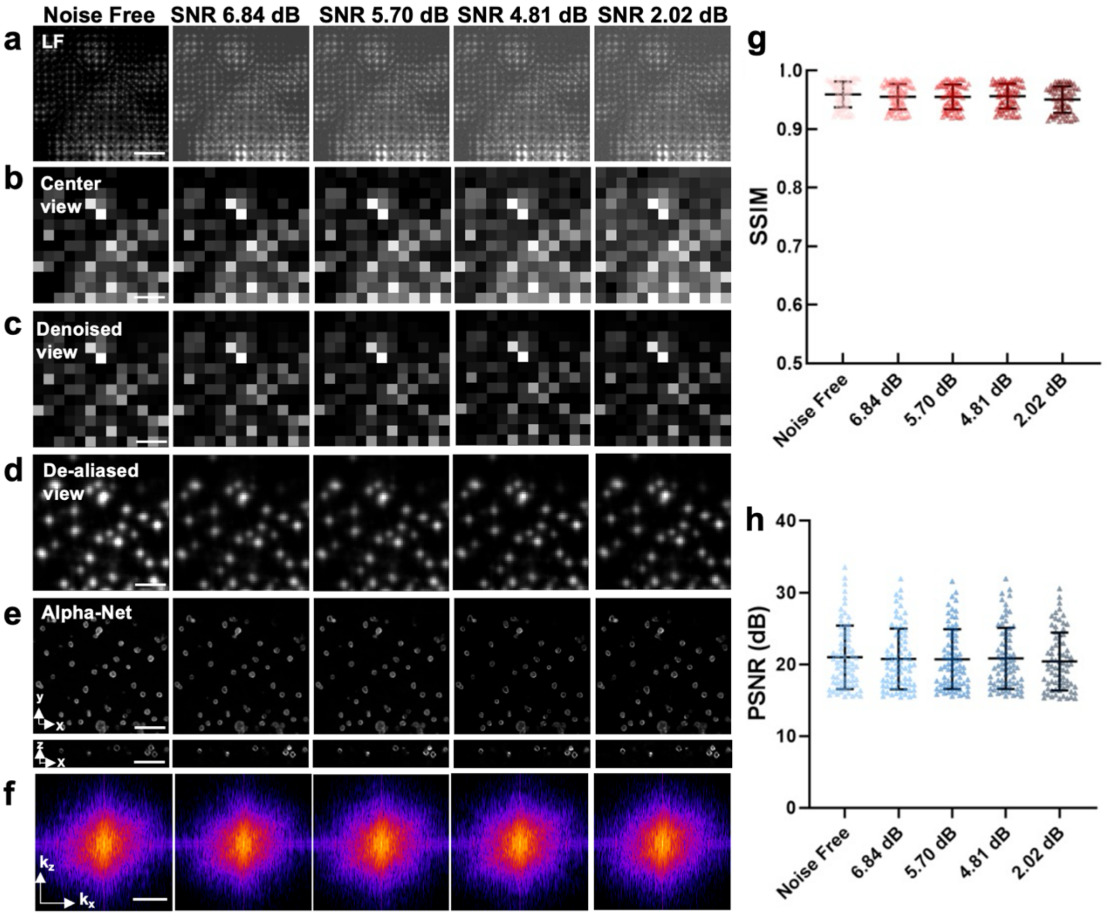

**Supplementary Figure 15. The robust denoising performance of Alpha-Net on LF images with different SNRs. a, b,** The semi-synthetic LF images (**a**) and their center views (**b**) with different SNRs ranging from 6.84 dB to 2.02 dB. **c, d,** The corresponding center view of denoised and de-aliased LF reconstructed by Alpha-Net. **e,** The orthogonal MIPs of the Alpha-Net reconstructions for these noisy and low-resolution LF inputs. **f,** The corresponding Fourier spectrums of x-z MIPs of Alpha-Net reconstructions in **e**. **g, h** The reconstruction fidelity of Alpha-Net are demonstrated by calculating the SSIM and PSNR, with using Airyscan data as reference. n=75 are used for each analysis. The data are shown as mean with standard diviation (SD), and the spots on the box indicates all calculated values. Scale bar, 3 μm (**a-e**), 10 $\mu m^{-1}$ (**f**).

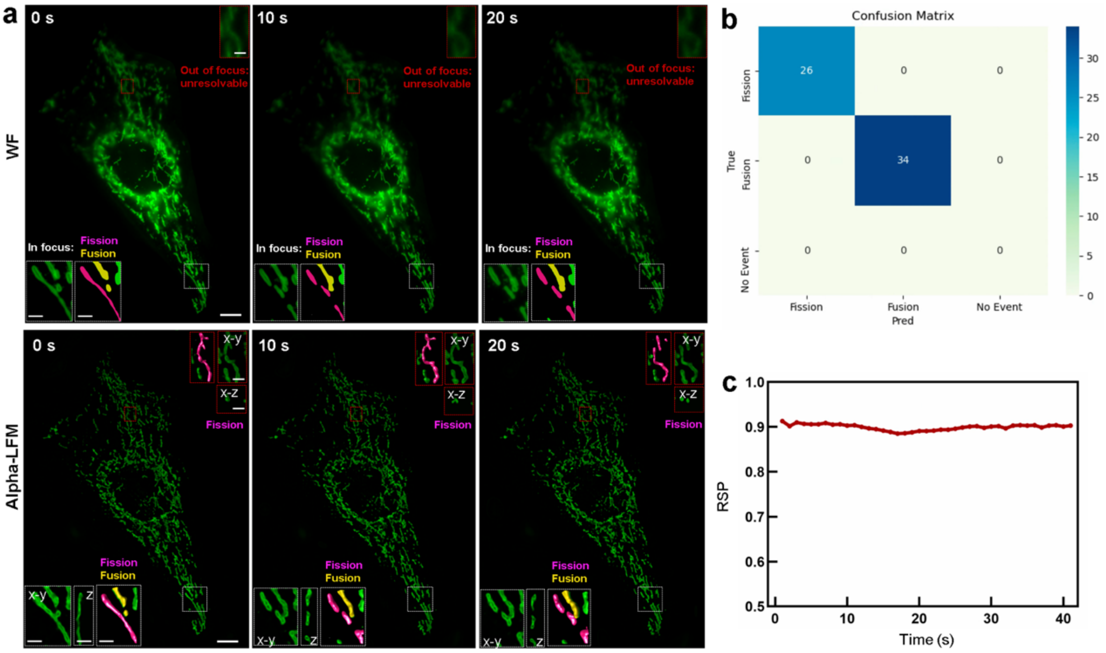

**Supplementary Figure 16. The fidelity validation of the biological discoveries by in-situ imaging the phenomena using hybrid LF-WF system in live samples. a,** The WF and Alpha-Net imaging results of mitochondrial outer membrane in live U2OS cell at three time points are shown. The insets show the magnified views indicated by dashed boxes. The white boxes indicate the region of interest (ROI) within the focus range of WF, while red boxes represent areas that out of focus in WF. The segmentations of the ROIs are shown in the insets with magenta representing fission events and yellow representing fusion events. Scale bar, 10 μm and 2 μm (insets). **b,** The confusion metrics quantifying the accuracy of identifying biological phenomenon (including the mitochondrial fission, fusion and no event) based on the results of our Alpha-LFM. By using the identification results of in-focus WF images as ground truth, our prediction results demonstrate an accuracy of 100%. The color bar represents the number of events. **c,** The RSP values quantifying the SR accuracy of the reconstructions at different times using in-focus WF images as reference.

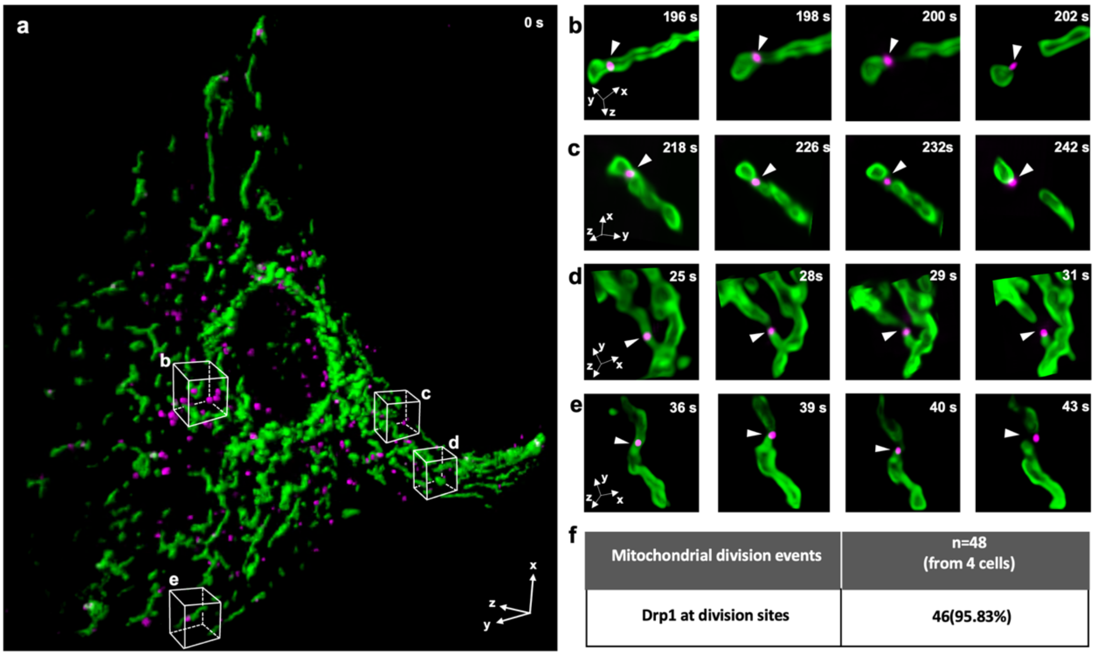

**Supplementary Figure 17. Accuracy validation of mitochondrial fission events identified by Alpha-LFM using live cells co-expressing Drp1 oligomers and mitochondria. a,** Dual-color volume rendering of a live U2OS cell co-expressing outer membranes of mitochondria (tagged with Tomm20-EGFP) and Drp1 oligomers (mCherry-Drp1) acquired using Alpha-LFM. **b-e,** Time-lapse images of Drp1-marked mitochondrial fission events occurred in ROIs indicated in **a**. **f,** Percentage of mitochondrial fission events marked by Drp1 (48 fission events from 4 cells).

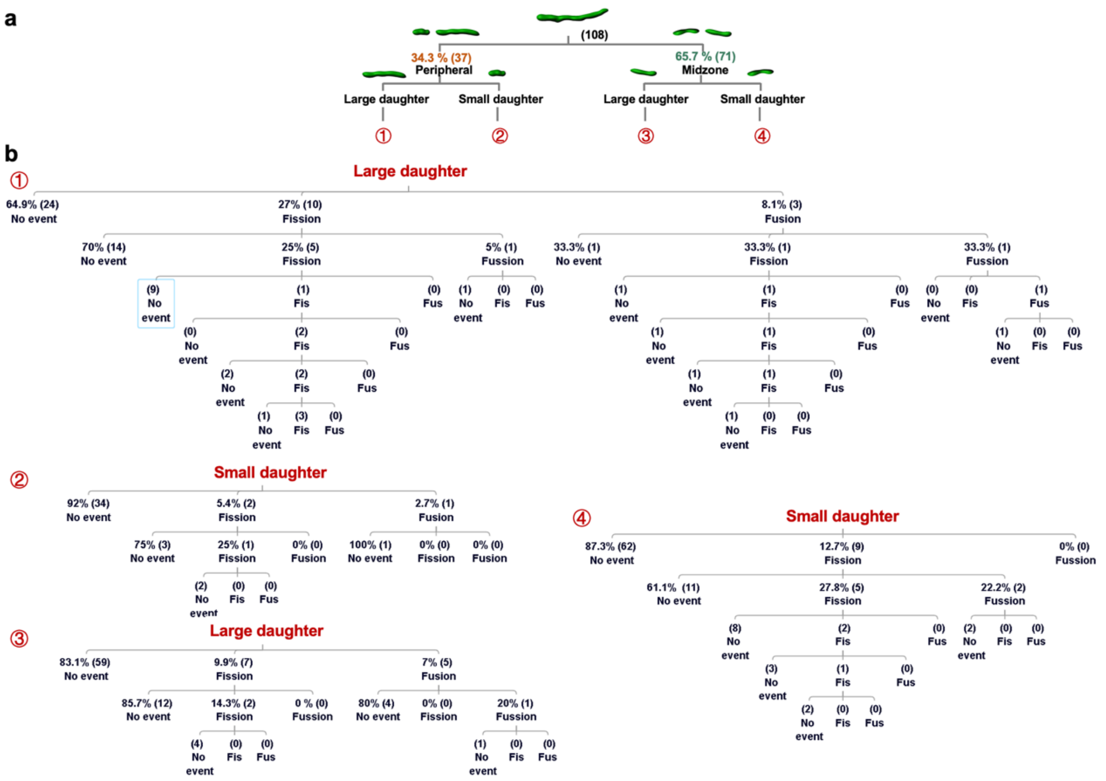

**Supplementary Figure 18. The fates of daughter mitochondria from peripheral and midzone fissions.** Large-scale lineage tracing diagram that depicts the detailed fates of 7-generation daughter mitochondria during 30 minutes, revealing the diverse fates of the daughter mitochondria from peripheral and midzone fissions.

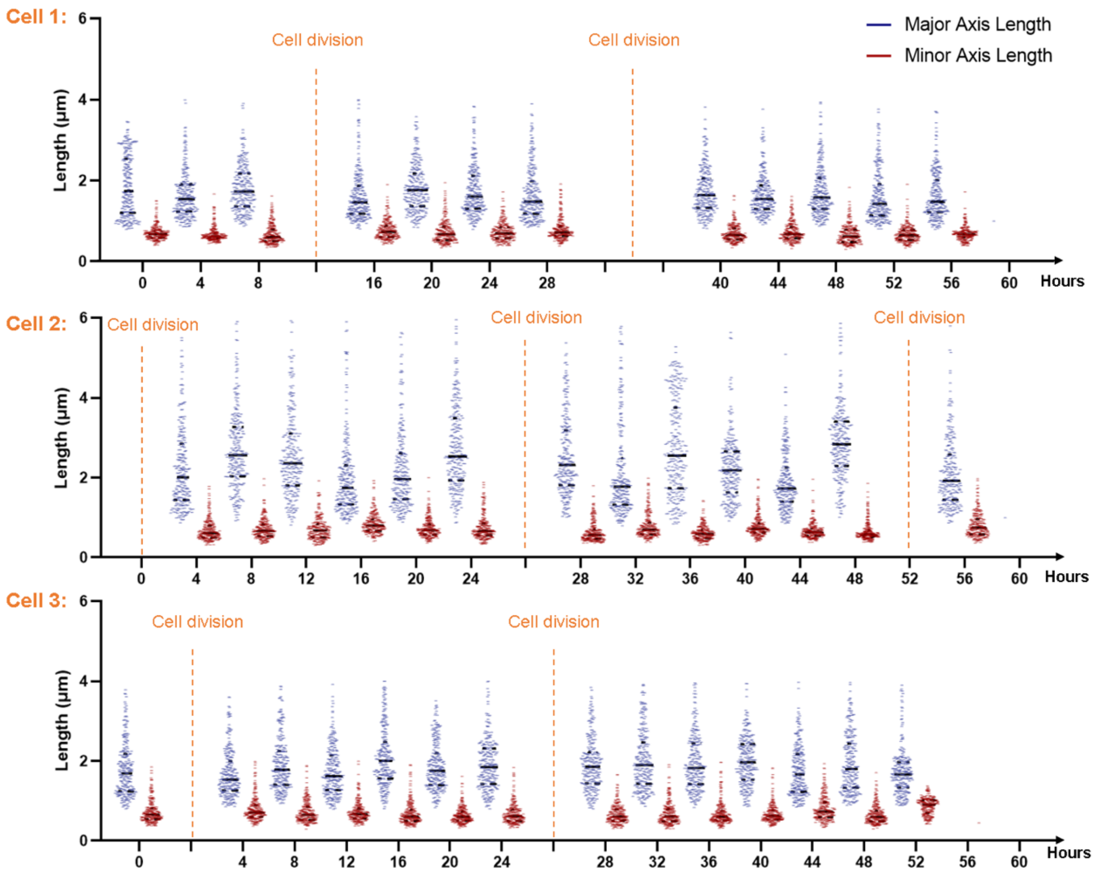

**Supplementary Figure 19. Quantification of morphology changes of 768 mitochondria in 3 living cells across 60 hours.** With the high-quality 4D SR image by Alpha-LFM, we successfully traced the length changes in the major and minor axes (blue and red) of mitochondria in three living cells throughout ~60 hours, a sufficiently long period encompassing at least one cell cycle. The orange dotted line indicates the timepoint of cell division.

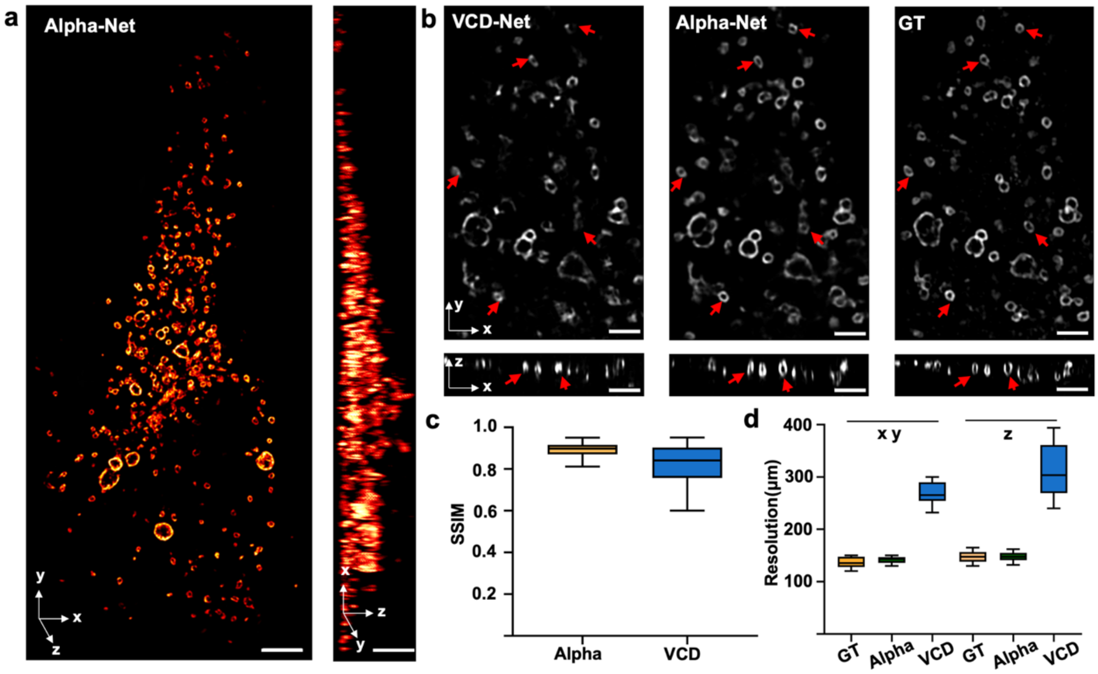

**Supplementary Figure 20. The performance demonstration of Alpha-Net using raw Airyscan data as training data. a,** The volume rendering of the Alpha-Net reconstruction results of semi-synthetic lysosome data using raw Airyscan data as training data. **b,** The orthogonal MIPs of the VCD-Net, Alpha-Net reconstruction results and raw Airyscan data (GT). **c,** SSIM metric quantitatively comparing the fidelity of Alpha-Net and VCD-Net using raw Airyscan data (GT) as reference (n=40 volumes). **d,** Decorrelation analysis quantifying the lateral and axial resolution of Alpha-Net, VCD-Net and GT results (n=40 volumes). The center line indicates the median, the box limits denote the lower and upper quartiles, and the whiskers represent the minimum and maximum values. Scale bar, 5 μm in **a**, 2 μm in **b**.

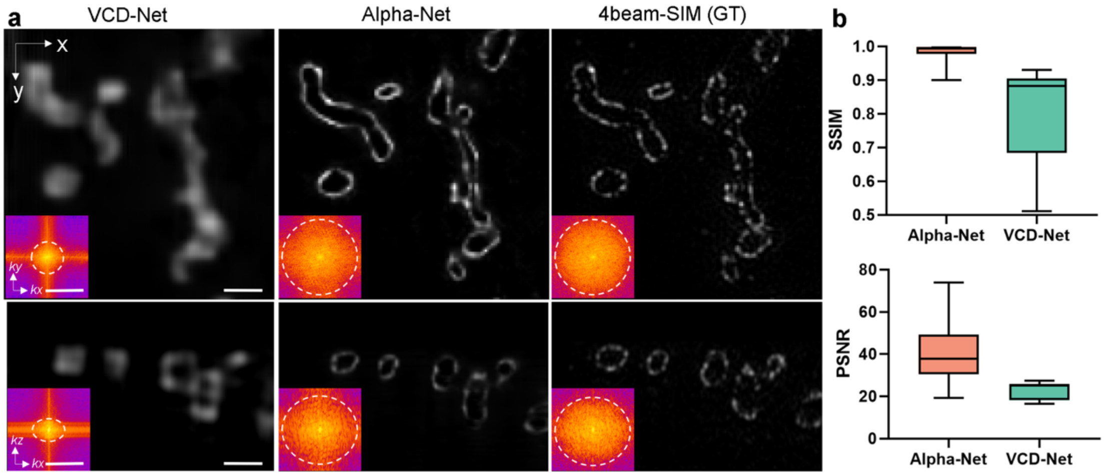

**Supplementary Figure 21. The performance of Alpha-Net using the data acquired from 4beam-SIM for training. a,** The 3D SR data of mitochondrial outer membrane were acquired from 4beam-SIM on fixed U2OS cells. We generated hierarchical training data and trained using our Alpha-Net framework and VCD-Net. The x-y and x-z MIPs of the reconstruction results of VCD-Net and Alpha-Net were shown with 4beam-SIM results as reference. The insets show the corresponding Fourier spectrums. The resolution of VCD-Net, Alpha-Net and 4beam-SIM are quantified to be 291 nm, 121 nm and 119 nm at lateral and 417 nm, 152 nm and 155 nm at axial. Scale bar, 10 µm (top in each panel), 2 µm (bottom left), 1/280 $nm^{-1}$ (bottom right). **b,** The reconstruction fidelity of Alpha-Net and VCD-Net are demonstrated by calculating the PSNR and SSIM, with using 4beam-SIM data as reference. n=30 volumes are used for each analysis. The center line represents the median, the box limits represent the lower and upper quartiles, and the whiskers represent the min and max value.

### Supplementary Notes

#### 1. Adaptive-learning physics-aware LFM (Alpha-LFM) framework

##### 1.1. Consideration in Light-field 3D Reconstruction

###### a. The complex inverse problem in light-field reconstruction

Light-field microscopy (LFM) is a computational imaging method that encodes spatial and angular information in a single 2D snapshot during the imaging process and 3D signals will be reconstructed by post-processing algorithms.

The LF encoding process from 3D spatial signals into a 2D snapshot is accompanied with the frequency aliasing and dimension degradation by the modulation of optical system and noise impacts from detectors or non-signal emission. This process can be formulated as $y = Hx + n + b$, where the vector $y$ represents the light-field image, the vector $x$ is the 3D distribution of signals, and $H$ is the 5D PSF (Point Spread Function) of LFM[5] with variables containing 3D spatial coordinates and 2D angular coordinates. n and b represent the Gaussian & Poisson noise in fluorescent detection[6] and background from dark current or defocus emission.

The LF reconstruction process is an inverse process of this forward imaging model to seek an accurate solution of 3D scene from this highly compressed two-dimensional (2D) input priors. However, the inference of high-dimensional (HD) images from low-dimensional (LD) images is an ill-posed problem since there exist infinite HD images that can be down sampled to the same LD images, the space of possible inversing solutions can be very large depending on the complexity of degradation model[7]. Hence, when dealing with the inverse problem of LF reconstruction, the space of candidate mapping function could be significantly large as the degradation encodes multidimensional information. In our study, light-field reconstruction beyond diffraction limit needs to inverse this compression with a space-bandwidth product (SBP) expansion over 600 times (Method). This highly ill-posed inversion problem poses a challenge on current LFM approaches.

###### b. The challenges in classical model-based approaches

Light field deconvolution (LFD) approach[5] adopted first-order, ideal imaging model to produce 3D results in an alternative projection manner. The optimization process can be described as minimizing a cost function (commonly formulated as the l2 norm: $|Hx - y|_2$; while such l2 norm would change termed as negative log-likelihood with Poisson noise assumption[8]) according to update all voxels of volume in image domain. The accurate prediction of this classical model-based approach largely relies on precise prediction of physical model. However, the variation of background and noise in LF images can significantly impact the construction of physical model, leading to a degradation in the quality of inductive results which is manifested as ringing artifacts and blur. Therefore, current deconvolution-based approach failed to break the large resolution gap between LF images and 3D SR image caused by intrinsic deficiency (aliased & uneven sampling pattern) in LFM imaging model[5], resulting a limited reconstruction resolution up to diffraction limit.

##### 1.2. Consideration in deep-learning approaches

Deep learning (DL) approaches have demonstrated superior performance in a wide range of image restoration tasks. Instead of relying on the precise physical modeling in classical model-based approach to solve the inverse problem, DL is a data-driven method that enables higher image enhancement capability through incorporating enormous high-quality label data and learning the nonlinear relationship between the source and label data.

However, these data-driven approaches encounter constraints in term of both enhanced capability and generalization ability. When dealing with complex inverse problems, ensuring accuracy is challenging. For instance, reconstructing a 3D super-resolution (SR) image from a 2D under-sampled light-field image with spatial bandwidth compressed by around 600 times, requires the recovery of various degradations brought by noise, resolution, and dimensionality reduction. This presents a highly intricate ill-posed problem with a huge solution space. Finding precise SR solutions in such a huge space is difficult and will result in in either limited resolution or reduced fidelity. Another typical challenge that supervised learning has to face is the generalization. This arises from the network's predominant focus on learning high-dimensional features of specific training samples, thereby limiting their applicability to the new samples. Here, we will specifically discuss the factors that influence the network fitting capability and generalization ability and analyze the strategies to enhance the fitting and generalization ability.

##### a. Enhancing capability of network

The network enhancing capability is evaluated by assessing its accuracy in reconstructing untrained data, referred to as the "generalization error" (less is better):

Let $E(f) = \mathbb{E}[\mathcal{L}_f(f(x), y)]$ is the generalization errors and $\hat{\mathrm{E}}(f)$ is the empirical loss (the loss computed from the limited training dataset). $\mathcal{X}$ and $\mathcal{Y}$ are all LF images and 3D ground truth in training dataset. $x \subseteq \mathcal{X}$ are input LF images while $y \subseteq \mathcal{Y}$ are target 3D stacks. $f$ is desired function (DL model) that would map $\mathcal{X}$ to $\mathcal{Y}$. $\mathcal{L}_f$ is the loss function that mapping from $\mathcal{X} \times \mathcal{Y}$ to [0, $C$] in which $C \subseteq R$. Based on Rademacher complexity[9], for any $\delta > 0$, with probability at least $1 - \delta$, the generalization error $E(f)$ satisfies for all $f \in \mathcal{H}$.s

$$E(f) \leq \hat{\mathrm{E}}(f) + 2\hat{R}_Z(\mathcal{H}) + 3C\sqrt{\frac{1}{2N} log\left(\frac{1}{\delta}\right)}$$

where $\hat{R}_Z$ is the empirical Rademacher complexity and $N$ is the number of samples. Let the $B(f)$ be the generalization bound of 3D SR reconstruction:

$$B(f) = 2\hat{R}_Z(\mathcal{H}) + 3C\sqrt{\frac{1}{2N} log\left(\frac{1}{\delta}\right)}$$

Based on above formulas, to reduce the generalization error, we need to reduce the empirical loss (related to the fitting ability of network) and the Rademacher complexity (related to the model complexity and objective function design) under a limited amount of training data.

Specifically, in our study, when solving the complex problem of inverting LF image to 3D SR image with a large space-bandwidth product ratio of ~600 times, the VCD model failed to fit the training data due to its limited model complexity and loss constraints. The reconstruction results showed high errors and degraded resolution (Fig. 2, Supplementary Fig. 6). To solve this complex inversion problem, one plausible way is to enlarge the function space to cover the optimal solution by increasing the model parameters. However, the increased model size produced higher Rademacher complexity, which "loosened" the generalization bounds. As shown in Supplementary

Note Figure 1.1, we trained three VCD net with different model size ("VCD-S", "VCD-M" and "VCD-L"). The training errors decreased with continuous increasement of the quantities of model parameters (Empirical loss: VCD-S < VCD-M < VCD-L). However, the increased model complexity would magnify the Rademacher complexity simultaneously. Therefore, based on the formula of upper bound of generalization error, the reduction of empirical training loss would be counteracted with the larger Rademacher complexity. The evaluation results in Supplementary Note Figure 1.1 also showed the variation trends of generalization errors: For the VCD-S, the small model size made network hard to fit the external morphology of mitochondria, manifested as the smaller width of intensity plots; For the VCD-M, although the larger model size facilitate the reduction of training errors while reconstructed the morphologies, it failed to resolve the hollow structure of membranes; For the VCD-L, though the further enlarged parameters reduced the training errors again, the validation results exhibited the same false structure with VCD-S one because of the counteraction of model complexity and empirical loss. On the contrary, our proposed Alpha-Net maintained the outer-morphology of mitochondria and resolved the hollow structure of membrane. We attributed this difference to the incorporation of physics priors into network design, data generation and optimization objective function.

Further, the more trainable parameters also complicated the optimization problem. For example, extending the depth and width of network would produce higher training errors than shallower model[10]. This optimization problem in solving highly ill-posed 3D reconstruction task impedes the accuracy of inductive results.

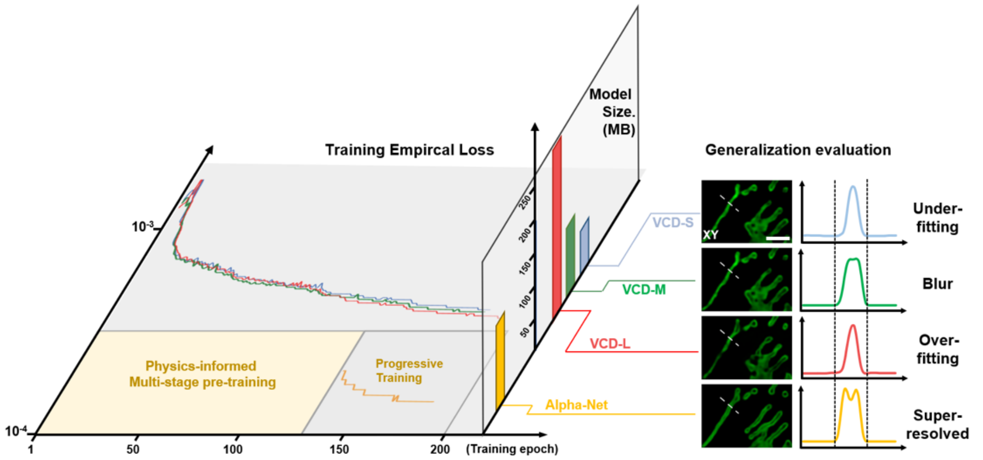

**Supplementary Note Figure 1. 1 Model comparison in training loss, network parameters and generalization evaluation.** Left: The plots of training errors of VCD-S (gray curve), VCD-M (green curve), VCD-L (red curve) and Alpha-Net (orange curve). Middle: The model size of four networks. Right: The validation results of mitochondria outer-membrane by the four trained model. Scale bar: 2 μm.

**b. Network generalization across different samples**

The data-driven learning algorithm could inverse the image distortion by learning the high-dimensional information from enormous label data. However, It inevitably suffered the errors when facing the large domain gap between the inference data and training data[11]. Specifically, in light-field 3D reconstruction, current VCD approaches[12, 13] couldn't be generalized to other samples with different structures. For example, we trained a VCD model on mitochondria outer-membrane, termed $model_{mito\ om}$. This model successfully resolved the structure of mitochondria with higher spatial resolution and fidelity. However, when applied $model_{mito\ om}$ on lysosome, the results exhibited obvious structural errors, such as the line structure artefacts. Such errors derived from the inductive assumption of data structure in previous model, which were termed as data bias. To reduce the errors in cross-domain validation, the transfer learning was proposed to apply the related but different knowledge containing in trained DL model to improve the performance of target learners[14, 15]. This process involves pre-training a model with massive paired data and fine-tuning the model with a small set of target data. Usually, the data in pre-training stage is easy accessible while the latter target data is hard to be obtained[15]. In light-field 3D reconstruction, a practical way is to train a model with paired SR-3D & LF dataset of organelle A and tune the model with paired SR-3D & LF dataset of organelle B. However, when observing the dynamics of a new sample with LFM, such transfer-learning paradigm requires to construct a complex in-situ SR & LFM microscope to capture the high-resolution details of new sample.

#### 1.3. Development of Alpha-LFM framework

As discussed in previous section, although data-driven DL methods exhibit superior enhancement capabilities compared to physical-model-based approaches, its reconstruction fidelity when solving complex inverse problem and generalization ability on unseen samples pose significant challenges. In addition to the data bias, incorporating more physical bias in the data-driven DL methods will improve the reconstruction fidelity and generalization ability of the network[16-19]. Since the model complexity and data priors are highly related to the generalization error of network (as discussed in *section 1.2)*, we established a physics-aware deep learning framework to incorporates physical bias in the model design and training data, therefore reducing the generalization error.

To appropriately enhance the model complexity and incorporate various data prior, we decomposed the complex light field inverse problem into multiple subtasks and designed a progressive multi-stage network to solve the subtasks with hierarchical data guided according to the physics model of the light-field encoding process. To permit the implementation of this framework to inverse the highly ill-posed LF problem, we specifically integrated physics priors into data synthesis, model design and objective function: **(i)** physics-embedded decomposed strategy; **(ii)**hierarchical data synthesis strategy; **(iii)** light-field aware multi-stage network; **(iv)** decomposed-progressive optimization strategy; **(v)** Adaptive tuning in a weak supervised manner, thereby enabling generalizable super resolution 3D reconstruction of LF images.

**a. Physics-embedded decomposed strategy of the framework**

The first challenges to achieve 30-fold super resolution (SR) from extremely under-sampled LF images lie in the decomposed learning strategy of the image inversion and the corresponding training data generation approach. The idea of breaking down the complex super-resolution light-field reconstruction procedure into multiple tasks like denoising, VCD reconstruction and 3D super resolution is quite straightforward, but it can't perform well. This is because such a procedure only utilizes spatial features and fails to incorporate angular constraints, which are essential for ensuring the accuracy of light-field reconstruction. Meanwhile, the final super-resolution module in 3D is highly computational demanding, leading to low inference speed of several minutes per volume. In the Supplementary Fig. 1, we compared the design and performance of VCD-Net, the easy idea of (Denoising+VCD+3D SR) and our Alpha-Net with (Denoising+De-aliasing+SR VCD). Alpha-Net shows notably enhanced reconstruction fidelity (Supplementary Fig. 1b, c) by incorporating angular constraint in the LF De-aliasing network, and achieves nearly four-order-of-magnitude higher inference speed by avoiding frequent use of complex 3D blocks (Supplementary Fig. 1b, c).

**b. Physics-embedded hierarchical data synthesis**

To permit the implementation of the decomposed framework, we developed a physics-embedded hierarchical data synthesis pipeline to generate corresponding hierarchical training data, including the raw LF, denoise LF, de-aliased LF and 3D SR. The pipeline is developed to synthesize the raw LF, denoise LF and de-aliased LF from 3D SR data based on the physical model without the requirement of a complex system to in situ acquire SR 3D and LF data (Supplementary Fig. 2).

The main challenge of the data synthesis lies in the absence of a method for generating de-aliased light-field images that can serve as ground-truth guidance for the de-aliasing task. We address this issue by pioneering a sub-aperture shifted light-field projection (SAS-LFP) method. As the discussion in Section 1.1, the formation of LF captures underwent the spatial-angular modulation and noisy & background contaminations. The first term could be factorized to limited-angle projection and spatial aliasing by low-pass filters (microlens). Inspired by the subpixel super-resolution in light-field photograpy[20, 21], we developed a sub-aperture shifted light-field projection (SAS-LFP) strategy to generate the de-aliased LF images served as the "de-aliasing priors". As shown in (*Step ii~iii*, Supplementary Fig. 2), because the low spatial resolution of extracted view image majorly caused by the low sampling rate of period arrangement of microlens array (severely below the Shannon-Nyquist limit), we shifted the SR 3D volumes by the distance less than the size of a single lenslet along horizontal and vertical direction, and projected the shifted 3D volumes based on light-field forward model, yielding multiple light field projections (LFPs) that contain "subpixel" information. In our setups, there were 15 pixels behind a single lenslet (after light-field image rectification) and the shift-times were 25 (5 x-shift and 5 y-shift). The shifted LFPs are then realigned to generate De-aliased LF images according to the image-shift trajectory, which enables recovering the lossy spatial resolution caused by aliasing.

This SAS-LFP method enables the generation of semi-synthetic de-aliased LFPs from SR image stacks and also serves as the cornerstone of our physics-embedded hierarchical data synthesis strategy (Supplementary Fig. 2). It is only through the integration of SAS-LFP that we can effectively implement the new decomposition model design of Alpha-Net to achieve superior performance (Supplementary Fig. 1).

Subsequently, we choose the non-shifted LFP as Clean LF to act the GT in denoising task. And then Gaussian & Poisson noise and background were added to the Clean LFs to mimic experimental LF captures: The intensity of Clean LFs were first rescaled to the same range of experimental LFs, and various levels of noise was applied on the rescaled LFs, according to the SNR (signal-to-noise ratio) variation of experimental LFs during long term observation. Then, the fluorescent background was added on such noisy images, according to the SBR (signal-to-background ratio) in raw LFs. As shown in Supplementary Note Figure 1.2, the high fidelity between Synthetic LFs and Experimental LFs validated the rationality of our synthetic pipeline.

With above data synthesis strategy, the generated hierarchical data could accurately describe the formation of experimental LFs and serve as the data prior in each local network.

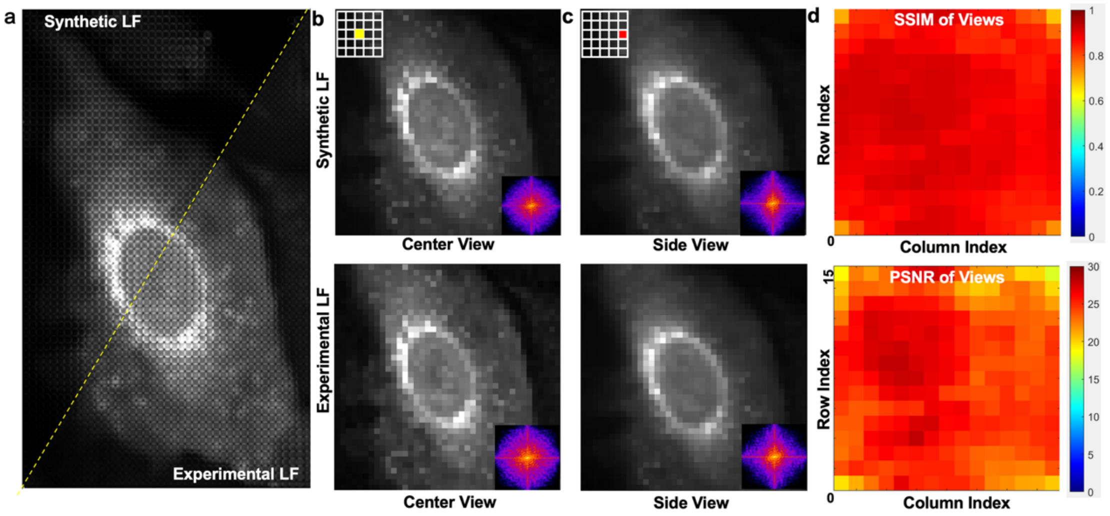

**Supplementary Note Figure 1. 2. The rationality verification of the data synthesis strategy. a,** The LF image synthesized using our synthetic pipeline (Synthetic LF) and acquired using our LFM systems (Experimental LF) are shown. **b,** The center and side views of Synthetic and Experimental LF images. The inset show the corresponding spectrum in Fourier domain. **c,** The fidelity map of views in Synthetic LF image are demonstrated by calculating SSIM and PSNR, with using Experimental LF image as reference. The color represents the SSIM and PSNR value.

#### c. Network design of Alpha-Net

The encoded LF images contains both 2D spatial and 2D angular information, which is highly correlation with the lateral and axial resolution in classical model based results. So, how to convey the 4D features without information loss and utilize the correlation between angular and spatial components in the process of network inference is key to our physics-informed network design. ***For LF denoising task***, the SNR (signal-to-noise ratio) variation among different LF views impeded the implementation of conventional denoised network only based on images denoising in spatial domain, like CARE[22], RCAN[23]. Inspired by the weighted deconvolution for 3D reconstrution[24] and dual attention strategy[25], we adopt learnable weights to wisely balance the contribution of various views. Specifically, a view-attention denoise module was introduced into conventional RCAN model (Supplementary Fig. 3) and adaptively captured discriminative noise features across different views. This dual-attention mechanism helped to a substantial improvement of SNR and preserve the yield higher resolution compared to spatial denoising strategy. ***For LF de-aliasing task***, the geometry consistency across input views mut be preserved after images super-resolution. Based on the spatial && angular information of LF, we split the network into two branches, and adopted spatial and angular convolution[26] respectively to extract features. ***For 3D reconstruction***, a light field reconstruction module is revised based on our VCD[12] to reconstruct the SR LF images, which incorporated the angular and spatial feature extraction operation into primary feature extraction module. Such dual-domain features were then interpolated and fused before U-Net block. Besides, the multi-scale convolution blocks were added to the encoder layer of U-Net which efficiently enlarged the receptive field of network and enhanced the fitting ability for super-resolution 3D reconstruction task.

**d. Decomposed and progressive training**

Alpha-Net is composed of 3 sub-networks tailored to distinct optimization tasks. To balance the independence (preserve the accuracy of each sub-network) and collaboration (ensure the low errors of 3D reconstruction), we devised a decomposed-progressive optimization strategy that consists of sub-networks pretraining and joint-optimization on cascaded network, during Alpha-Net optimization process.

i. Sub-networks pre-training

Firstly, the view-attention based denoising network was used to convert the input noisy LFs into “clean” LFs. To train this model for each organelle, ~1000 pairs of simulated noisy LFs and corresponding noiseless label data with a patch size of 360 × 360 were required. We chose Adam optimizer with an initial learning rate of $1\times1e^{-4}$ and a batch size of 1. As for the weighted L2-L1 loss function, the ratio of L2 loss was set to 1.0 while 0.1 for L1 loss. During training, the learning rate was reduced by half per 10 epochs. The total training epoch was set to 51 to obtain model well-convergence.

Secondly, The LF de-aliasing network was applied for increasing the sampling rate of extremely down sampled LF views. We used ~1000 pairs of noiseless LFs and scanning-LFs to train this model. The lateral upsampling factor was 5 in our setups and the patch size of input LF was 360 × 360. The weight ratio of parallax loss was set to 0.1. For training parameters, the learning rate was $5\times1e^{-4}$ and training epoch was set to 51.

Thirdly, the 3D reconstruction module took the upscaled LFs as inputs and predicted corresponding super-resolution stacks. With data augmentation (axial shift), ~1500 pairs of scanning-LFs (patch size: 1800 × 1800) and SR 3D stack (patch size: 720 × 720 × 161) were obtained. Then, this model was trained by Adam optimizer for 151 epochs with an initial learning of $5\times1e^{-4}$. During training, the learning rate would decay by half per 50 epochs.

ii. Decomposed progressive optimization

After pre-training finished, 3 trained sub-networks were obtained. To mitigate the mismatch among these models during the optimization of cascaded networks, we adopted a grouped joint-optimization strategy to achieve well converge. The total number of training pairs was ~1500. The patch size of each data was the same with the size used in pre-training phase. Based on the progressive restoration strategy, we deployed 3 Adam optimizers to update networks parameters. The $1^{st}$ optimizer was dedicated to updating the parameters of LF denoise network based on the denoising loss function. The $2^{nd}$ optimizer used the weighted denoising & de-aliasing loss function to update the parameters of LF denoising & de-aliasing networks. The $3^{nd}$ optimizer was employed for joint optimization on cascaded networks with the weighted denoising & de-aliasing && 3D reconstruction loss function. The initial learning rate was set to $5\times1e^{-4}$. During training, all of these optimizers shared the same learning rate and sequentially update corresponding network parameters over a training epoch of 101. The learning rate was halved per 50 epochs. To avoid model over-fitting, we used early-stop strategy to save the best epoch (calculated on validation data) during training. Compared with conventional one-step optimization strategy (equal to the $3^{rd}$ optimizer), our proposed progressive optimization achieved better reconstruction quality and network convergence (Supplementary Fig. 6).

##### e. Workflow of adaptive network tuning

As shown in Supplementary Fig. 9, taking the "mitochondria-to-lysosome" as an example, the captured LF and WF of lysosomes act the lateral constraints to prevent the structural errors in network prediction while the synthetic mitochondria data provides the volumetric priors to regularize the model with the preservation of 3D SR reconstruction ability.

To get the LF-WF training pairs, the users only need to mount an additional camera on the right optical port and therefore can easily acquire *in situ* LF-WF data of the new sample. Due to the minor misalignment between two modalities, the geometry relation of captured LF-WF data needed to obtained before network training. To register the WF images with the LF images, the reconstructed results of trained samples and corresponding WF can used to apply registration to get the affine transformation matrix. This step can be conducted using "*Registration*" plugin in Fiji. Since the register transformation between WF and LF are fixed once the system has been built, the register matrix can be applied to all the acquired LF-WF pairs. In our experiment, the adaptive tuning strategy only needs the paired data captured from15 cells. For live cells, the users can capture 5 *in situ* WF and LF frames of the dynamic organelles from 3 cells for training. For fixed cells, the users can acquire 15 *in situ* WF-LF pairs for adaptive training.

After the data acquisition, preprocessing of these data was needed to start model fine-tuning: 1) Subtract the fluorescent background of WFs and choose the in-focus region of WF captures. 2) Deconvolve the WFs with simulated PSF (derived from "PSF Generator" in Fiji), which was equal to the maximum projection of the signals in the limited DoF of WF. 3) Make affine transformation on WFs based on the registration parameters obtained in data acquisition phase. 4) Crop these paired data to small patches to generate training dataset. Then, this dataset was fed into the network to fine tune the trained model. During training, the mean-square-errors (MSE) between the down-sampled maximum projection of network's 3D prediction and input WF was calculated as lateral structural constraint. It's worth noting that only part of the slices of the network's 3D prediction within the DOF of the WF are projected. Furthermore, to maintain the mapping function between the 3D reconstructions and LF and prevent the over fitting by the lateral constraint, few 3D training data used for the raw large model should also be incorporated into the training process, acting as the volumetric constraint (more precisely, volumetric regularization term). In our experiments, we adopt alternate optimization strategy between two datasets.

#### 2. Evaluation of Alpha-Net

##### 2.1. Fidelity evaluation of network prediction

Currently, CNN shows superior capability of learn the mapping function from the low-resolution images to high-resolution images with high fidelity. However, when facing the complex inverse problem in LFM, finding precise SR solutions is difficult for end-to-end VCD network and result in in limited resolution or reduced fidelity (as validated in Fig. 2).

To reduce the solution space, Alpha-Net include the physical model as prior into the model design in a progressive supervised manner, instead of just include onefold data prior as classical

CNN. Through the physics-embedded design, we demonstrate that Alpha-Net can accurately infer the 3D distribution of signals with sub-diffraction-limited resolution from 2D light-field image by quantitatively validating the fidelity with compared to GT on both test dataset and experimental data and demonstrating the accuracy of judging the biological phenomenon.

Firstly, we quantified the fidelity and resolution of the reconstruction results during the network's convergence process by calculating the SSIM with GT and cut-off frequency. While VCD can not find a precise SR solution through end-to-end supervision, our Alpha-Net rapidly approach the global optimum (Figure 2a, b), indicating a high structure similarity with ground truth data (super resolution 3D volume) and super resolution. The well convergence of Alpha-Net demonstrated the fidelity of network prediction.

Secondly, to further demonstrate the fidelity of network prediction in real world, we conducted in-situ 3D SR and LF imaging experiments to evaluate the fidelity of Alpha-Net. Through quantifying SSIM of the network inference result and enhanced Airyscan data in different cells and regions, we demonstrated that the Alpha-Net can accurately infer the 3D distribution of signals with sub-diffraction-limited resolution from 2D experimental LF (Fig 2i-k).

Finally, we evaluated the fidelity of network inference when conducting biological downstream analysis. As shown in Supplementary Fig. 16, we conducted in-situ imaging of mitochondrial fission and fusion using our hybrid wide-field and LFM system. As depicted in the figure, although the wide-field imaging results exhibit excessive out-of-focus blurs in areas with dense signals (red boxes), the in-focus signals verify the fidelity of Alpha-LFM (white boxes). We have identified fission and fusion activities in the focus area of the wide-field image, which are consistent with the findings in the results of Alpha-LFM. Meanwhile, the blurred area in 2D wide-field image can be clearly resolved in 3D Alpha-Net reconstruction, verifying the advantage of our approach for live-cell imaging. We further conducted statistical analysis and determined the confusion matrix for the detection of mitochondrial fission and fusion event, showing a high accuracy of 100% of Alpha-LFM.

In conclusion, through validating on both structure similarity and accuracy of identifying biological event, our Alpha-Net shows its robust performance of reconstructing HiFi 3D SR distributions.

#### 2.2. Model uncertainties estimation

Besides of the resolution and localization accuracy evaluation on single model prediction, we also conducted analysis on model uncertainties.

A neural network is a highly non-linear function $f_\theta$ parameterized by model parameters $\theta$ that maps from input measurable set $\mathcal{X}$ to output measurable set $\mathcal{Y}$. In supervised learning, the neural function is obtained via optimizing an objective loss function computed on the finite training dataset $D \subseteq \mathcal{D} = \mathcal{X} \times \mathcal{Y}$. For a new data sample $x^* \subseteq \mathcal{X}$, the network $f_\theta$ trained on $D$ can predict corresponding target $y^* = f_\theta(x^*)$. During this process, there exist data uncertainties and model uncertainties[27]. The former one is caused by the randomness during data collection and measuring system. For example, in regression tasks, the noise in training dataset $D$ results in the data uncertainties. The latter one is the lack of knowledge about model, which caused by the model shortcomings[27]. For example, the randomly initialized network parameters would produce slightly different predictions when repeatedly training network.

In this study, the data uncertainties were derived from the random noise contamination in fluorescent imaging, denoted by the various SNR of captured light-field images. To investigate the impact of input randomness on final 3D reconstruction, we test the robustness of Alpha-Net under different SNRs on lysosome data (Supplementary Fig. 15). The results show that our method successfully reconstructed the hollow structure of lysosome and achieve high fidelity under different SNRs (2.02 dB ~6.84 dB), which revealed our method enabled the uncertainties reduction in collected data.

To evaluate the model uncertainties, we adopted the model ensemble approach[28] that computes the variation and mean of multiple ensemble members. Specifically, we independently trained Alpha-Net 4 times to obtain different models while each model was trained with randomly initialized parameter and data order. Then we calculated the per-pixel mean value and standard deviation from these network predictions:

$$Mean = \frac{1}{N} \sum_{n=1}^{N} f_{\theta_n}(x)$$

$$SD = \sqrt[2]{\frac{1}{N} \sum_{n=1}^{N} \left(f_{\theta_n}(x) - Mean\right)^2}$$

where $f_{\theta_n}$ is the n$^{th}$ model parameterized by network parameters $\theta_n$ and $x$ is the input LF image (each model adopted the same input). $N$ is the number of trained models. Then, we computed the coefficient of variation (CV) by:

$$V_\sigma = \frac{SD}{Mean}$$

The larger $V_\sigma$ means the higher uncertainties of network. We conducted the analysis on the 3D reconstructions of lysosome as shown in **Supplementary Note Figure 2**. The results shown higher similarity between different network predictions and less model uncertainty (CV is 0.0238 for axial projections while 0.0086 for lateral projections).

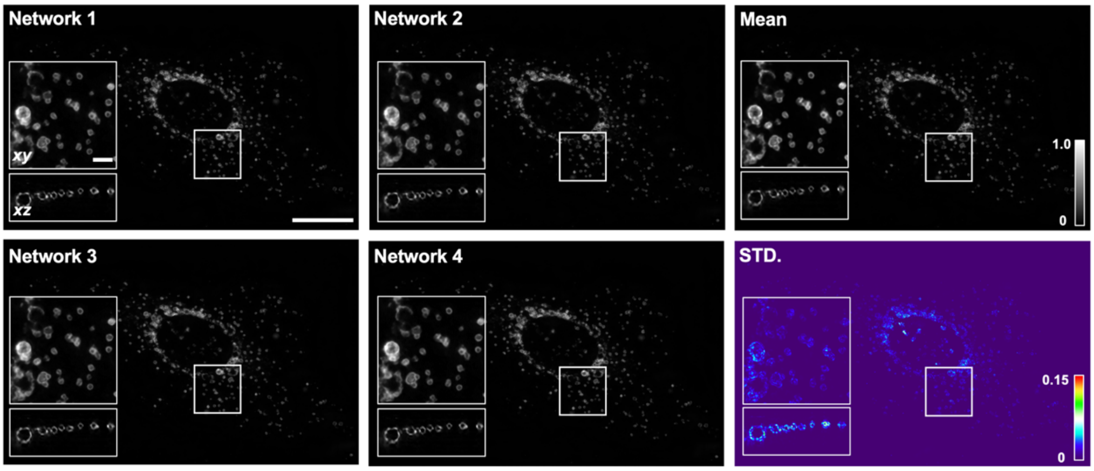

**Supplementary Note Figure 2. Model uncertainty of Alpha-Net assayed with lysosome 3D reconstructions.** From left to right, we shown the four network predictions, the pixel-wise mean and the pixel-wise of standard deviation (SD). The insets of each panel depicted the magnified view of the average projection of lateral planes and axial planes of 3D volumes, which revealed less disagreement among the hollow structure of four network reconstructions. The coefficient of variation (CV) is 0.0086 for the lateral projections while 0.0238 for the axial ones, indicating small variation among different networks with random initialization. Scale bar: 20 μm in whole-FOV projections while 2 μm in insets.

### Supplementary Videos

**Supplementary Video 1. Principle and pipeline of Alpha-Net for 3D super-resolution reconstruction.** The first part of video shows the physics-embedded hierarchical data synthesis process, including 3D SR, De-aliased LF, Clean LF and Noisy LF. The second part of the video shows the network architecture of light-field-aware Alpha-Net and corresponding optimization process. The last part of video shows the principle of network fine-tuning based on trained Alpha-Net.

**Supplementary Video 2. Comparisons of LF reconstruction results of mitochondrial outer membranes in live U2OS cells by VCD, VS-LFM and Alpha-Net.**

**Supplementary Video 3. Low-photobleaching imaging of lysosomes (tagged with Rab7-EGFP) in a live U2OS cell encompassing 40000 volumes.**

**Supplementary Video 4. Imaging of peroxisomes (tagged with SKL-mApple) in a live U2OS cell at 100 vps.**

**Supplementary Video 5. Imaging of endoplasmic reticulum (ER, tagged with Sec61β-EGFP) in a live COS-7 cell at 100 vps.**

**Supplementary Video 6. The time-lapse simultaneous dual-color 3D SR imaging of the outer membranes of mitochondrial (tagged with Tomm20-EGFP) and lysosomes (tagged with Rab7-mCherry) in a live U2OS cell via Alpha-LFM reveals the typic lysosomes mediated mitochondrial fusion and fission events.**

**Supplementary Video 7. The quantitative analysis of the velocity of mitochondrial fission and fusion with or without lysosomes contact enabled by 5D SR imaging of Alpha-LFM.**

**Supplementary Video 8. Long-term imaging of mitochondria (tagged with Cox4-EGFP) and chromosomes (tagged with H2B) via Alpha-LFM throughout 60 hours.**

**Supplementary Video 9. The time-lapse imaging of mitochondria (tagged with Cox4-EGFP) during 30 minutes via Alpha-LFM and detailed lineage tracing of the selected single mitochondrion with 7 generations traced.**

### Supplementary References

1. Wang, Y. et al. in Computer Vision – ECCV 2020. (eds. A. Vedaldi, H. Bischof, T. Brox & J.-M. Frahm) 290-308 (Springer International Publishing, Cham; 2020).
2. Wang, Z. et al. Real-time volumetric reconstruction of biological dynamics with light-field microscopy and deep learning. *Nat Methods* **18**, 551-556 (2021).
3. Weigert, M. et al. Content-aware image restoration: pushing the limits of fluorescence microscopy. *Nat Methods* **15**, 1090-1097 (2018).
4. Chen, J. et al. Three-dimensional residual channel attention networks denoise and sharpen fluorescence microscopy image volumes. *Nat Methods* **18**, 678-687 (2021).
5. Broxton, M. et al. Wave optics theory and 3-D deconvolution for the light field microscope. *Opt Express* **21**, 25418-25439 (2013).
6. Zhang, Y. et al. in 2019 IEEE/CVF Conference on Computer Vision and Pattern Recognition (CVPR) 11702-11710 (2019).
7. Ledig, C. et al. in Proceedings of the IEEE conference on computer vision and pattern recognition 4681-4690 (2017).
8. Sage, D. et al. DeconvolutionLab2: An open-source software for deconvolution microscopy. *Methods* **115**, 28-41 (2017).
9. Mohri, M., Rostamizadeh, A. & Talwalkar, A. Foundations of machine learning. (MIT press, 2018).
10. He, K., Zhang, X., Ren, S. & Sun, J. in Proceedings of the IEEE conference on computer vision and pattern recognition 770-778 (2016).
11. Belthangady, C. & Royer, L.A. Applications, promises, and pitfalls of deep learning for fluorescence image reconstruction. *Nat. Methods* **16**, 1215-1225 (2019).
12. Wang, Z. et al. Real-time volumetric reconstruction of biological dynamics with light-field microscopy and deep learning. *Nat. Methods* **18**, 551-556 (2021).
13. Yi, C. et al. Video-rate 3D imaging of living cells using Fourier view-channel-depth light field microscopy. *Communications Biology* **6**, 1259 (2023).
14. Zhuang, F. et al. A comprehensive survey on transfer learning. *Proceedings of the IEEE* **109**, 43-76 (2020).
15. Liao, T. et al. A super-resolution strategy for mass spectrometry imaging via transfer learning. *Nature Machine Intelligence* **5**, 656-668 (2023).
16. Yanny, K., Monakhova, K., Shuai, R.W. & Waller, L. Deep learning for fast spatially varying deconvolution. *Optica* **9**, 96-99 (2022).
17. Huang, L., Chen, H., Liu, T. & Ozcan, A. Self-supervised learning of hologram reconstruction using physics consistency. *Nature Machine Intelligence* **5**, 895-907 (2023).
18. Li, Y. et al. Incorporating the image formation process into deep learning improves network performance. *Nature Methods* **19**, 1427-1437 (2022).
19. Guo, Y. et al. in Proceedings of the IEEE/CVF conference on computer vision and pattern recognition 5407-5416 (2020).
20. Bishop, T.E. & Favaro, P. The Light Field Camera: Extended Depth of Field, Aliasing, and Superresolution. *IEEE Transactions on Pattern Analysis and Machine Intelligence* **34**, 972-986 (2012).
21. Chan, W.-S., Lam, E.Y., Ng, M.K. & Mak, G.Y. Super-resolution reconstruction in a

computational compound-eye imaging system. *Multidimensional Systems and Signal Processing* **18**, 83-101 (2007).

22. Weigert, M. et al. Content-aware image restoration: pushing the limits of fluorescence microscopy. *Nat. Methods* **15**, 1090-1097 (2018).
23. Chen, J. et al. Three-dimensional residual channel attention networks denoise and sharpen fluorescence microscopy image volumes. *Nat. Methods* **18**, 678-687 (2021).
24. Lu, Z. et al. Phase-space deconvolution for light field microscopy. *Opt Express* **27**, 18131-18145 (2019).
25. Mo, Y., Wang, Y., Xiao, C., Yang, J. & An, W. Dense dual-attention network for light field image super-resolution. *IEEE Transactions on Circuits and Systems for Video Technology* **32**, 4431-4443 (2021).
26. Wang, Y. et al. in Computer Vision–ECCV 2020: 16th European Conference, Glasgow, UK, August 23–28, 2020, Proceedings, Part XXIII 16 290-308 (Springer, 2020).
27. Gawlikowski, J. et al. A survey of uncertainty in deep neural networks. *Artificial Intelligence Review* **56**, 1513-1589 (2023).
28. Lakshminarayanan, B., Pritzel, A. & Blundell, C. Simple and scalable predictive uncertainty estimation using deep ensembles. *Advances in neural information processing systems* **30** (2017).
